# Comparative mitogenomic analysis of subterranean and surface amphipods (Crustacea, Amphipoda) with special reference to the family Crangonyctidae

**DOI:** 10.1101/2023.04.27.538449

**Authors:** Joseph B. Benito, Megan L. Porter, Matthew L. Niemiller

## Abstract

Mitochondrial genomes play important roles in studying genome evolution, phylogenetic analyses, and species identification. Amphipods (Class Malacostraca, Order Amphipoda) are one of the most ecologically diverse crustacean groups occurring in a diverse array of aquatic and terrestrial environments globally, from freshwater streams and lakes to groundwater aquifers and the deep sea, but we have a limited understanding of how habitat influences the molecular evolution of mitochondrial energy metabolism. Subterranean amphipods likely experience different evolutionary pressures on energy management compared to surface-dwelling taxa that generally encounter higher levels of predation and energy resources and live in more variable environments. In this study, we compared the mitogenomes, including the 13 protein-coding genes involved in the oxidative phosphorylation (OXPHOS) pathway, of surface and subterranean amphipods to uncover potentially different molecular signals of energy metabolism between surface and subterranean environments in this diverse crustacean group. We compared base composition, codon usage, gene order rearrangement, conducted comparative mitogenomic and phylogenomic analyses, and examined evolutionary signals of 35 amphipod mitogenomes representing 13 families, with an emphasis on Crangonyctidae. Mitogenome size, AT content, GC-skew, gene order, uncommon start codons, location of putative control region (CR), length of *rrnL* and intergenic spacers differed between surface and subterranean amphipods. Among crangonyctid amphipods, the spring-dwelling *Crangonyx forbesi* exhibited a unique gene order, a long *nad5* locus, longer *rrnL* and *rrnS* loci, and unconventional start codons. Evidence of directional selection was detected in several protein-encoding genes of the OXPHOS pathway in the mitogenomes of surface amphipods, while a signal of purifying selection was more prominent in subterranean species, which is consistent with the hypothesis that the mitogenome of surface-adapted species has evolved in response to a more energy demanding environment compared to subterranean amphipods. Overall, gene order, locations of non-coding regions, and base-substitution rates points to habitat as an important factor influencing the evolution of amphipod mitogenomes.

## Introduction

Caves and other subterranean habitats, such as groundwater aquifers and superficial subterranean habitats (SSHs; Culver and Pipan 2009), represent some of the most challenging environments that exists on Earth. The primary characteristic of all subterranean habitat is the lack of light and associated photosynthesis (Culver and Pipan 2009; Soares and Niemiller 2020). Though some subterranean ecosystems are supported by chemoautotrophic production by microbial communities (Engel *et al*. 2004; Porter *et al*. 2009), chemoautotrophy rarely provides enough energy to support several trophic levels in most subterranean ecosystems (Poulson and White 1969; Culver and Pipan 2009). The primary source of energy input for many cave systems is the organic matter transferred from the surface hydrologically or by animals that frequently enter and exit caves (Simon and Benfield 2001; Culver and Pipan 2009), which drive the structure and dynamics of subterranean communities (Huppop 2000; Graening and Brown 2003; Huntsman *et al*. 2011). Although most subterranean ecosystems are largely thought to be energy-limited (Venarsky *et al*. 2014), food availability can be highly variable both among and within cave systems (Culver *et al*. 1995; Juan *et al*. 2010). Previous studies have shown that many subterranean organisms living in such energy-limited habitats have undergone several physiological and metabolic adaptations to sustain themselves during extended food shortages (Hervant *et al*. 1997; Issartel *et al*. 2010). Among these troglomorphic traits, low metabolic rate is a key adaptation that occurs in both terrestrial and aquatic fauna of the subterranean communities (Bishop *et al*. 2014; Nair *et al*. 2020).

Mitochondria are the primary sites of energy production in cells, generating ∼95% of the adenosine triphosphate (ATP) required for everyday activities of life through oxidative phosphorylation (OXPHOS; Das 2006; Shen *et al*. 2010; Yang *et al*. 2014). The mitochondrial genome—mitogenome—encodes 13 essential proteins including two ATP synthases (*atp6* and *atp8*), three cytochrome oxidases (cox*1, cox2,* and *cox3*), seven NADPH reductases (*nad1*, *nad2*, *nad3*, *nad4*, *nad4l*, *nad5*, and *nad6*), and cytochrome *b* (*cytb*) subunits. All mitochondrial protein-coding genes (PCGs) play a vital role in the electron transport chain (Boore 1999; Burger *et al*. 2003; Xu *et al*. 2007). Due to the unique characteristics of mitochondria, including maternal inheritance, small genomic size, absence of introns, and their surplus availability in cells, the use of mitochondrial DNA (mtDNA) loci and mitogenomes has a long history in population genetics, phylogenetics, and molecular evolution studies (e.g., Ballard and Pichaud 2014; Bourguignon *et al*. 2018; Zou *et al*. 2018). Previous studies have demonstrated a close association between mitochondrial loci and energy metabolism (Shen *et al*. 2009, 2010; da Fonseca *et al*. 2008; Zhang *et al*. 2013). Although considered to largely evolve under purifying selection, there is growing evidence that mitogenomes may undergo episodes of directional selection in response to shifts in physiological or environmental pressures (Botero-Castro *et al*. 2018; Sun *et al*. 2020) leading to improved metabolic performance under new environmental conditions (da Fonseca *et al*. 2008; Garvin *et al*. 2011; Welch *et al*. 2014). For example, previous studies that investigated varying selective pressures acting on mitochondrial PCGs of insects and mammals have revealed significant positive selective constraints at several loci that have comparatively increased energy demands (Shen *et al*. 2010; Yang *et al*. 2014; Li et *al*. 2018).

Similarly, other studies have shown the various adaptive mitochondrial responses of organisms surviving in extreme environments including the deep sea and Tibetan Plateau (Mu *et al*. 2018; Li *et al*. 2018; Sun *et al*. 2020). However, these adaptations can occur at different metabolic levels, not just mitochondrial metabolism (Beall 2007; Scott *et al*. 2011). Thus, variation in mitogenomes of species inhabiting different environments may reflect only a small portion of these adaptive metabolic changes. Despite this limitation, previous studies have detected signals of directional selection in the mitogenomes of organisms dwelling in contrasting habitats with varying energy demands (Peng *et al*. 2011; Yang *et al*. 2015; Li *et al*. 2016).

Amphipods (Class Malacostraca: Order Amphipoda) are one of the most ecologically diverse crustacean groups including over 10,000 species (Arfianti *et al*. 2018; Horton *et al*. 2020), occurring in a diverse array of aquatic and even terrestrial environments globally, from aphotic groundwater aquifers and hadal depths to freshwater streams and lakes in temperate and tropical forests, among other habitats (Bousfield 1983; Barnard and Karaman 1991). Several studies have demonstrated the genetic basis of subterranean adaptation in several taxa, including dytiscid diving beetles (Hyde *et al*. 2018), cave dwelling-cyprinid fishes (Wu *et al*. 2010; Dowling *et al*. 2002), anchialine cave shrimps (Guo *et al*. 2018), and cave isopods (Protas *et al*. 2011).

However, we still have a limited understanding of the mechanisms of subterranean adaptations in amphipods. Although physiological adaptations to challenging environments like cave and groundwater ecosystems have been well-studied in amphipods (e.g., Hervant *et al*. 1997; Nair *et al*. 2020), no studies to date have addressed the selective pressures and the molecular evolution mechanisms of mitochondrial energy metabolism loci in amphipods occupying caves and other subterranean habitats. Subterranean amphipods likely experience different evolutionary pressures on energy management due to lower levels of predation, lower food resources, and more stable environments compared to surface-dwelling taxa that generally experience higher levels of predation and energy resources (Qiu *et al*. 2012; Qu *et al*. 2013).

In this study, we compared the mitogenomes of surface and subterranean amphipods, including the 13 mitochondrial PCGs involved in the OXPHOS pathway to understand the potential molecular mechanisms of energy metabolism in this diverse crustacean group. Our aims were to test whether the mitochondrial PCGs showed evidence of adaptive evolution in subterranean environments in amphipods. We tested the hypothesis that the mitogenome of surface-adapted amphipods will be imprinted by mitogenomic adaptations to the energy demanding environment with greater signal of directional selection when compared to their subterranean counterparts.

We compared base composition, codon usage, gene order rearrangement, conducted comparative mitogenomic and phylogenomic analyses, and examined evolutionary signals using publicly available amphipod mitogenomes. In particular, we focused on the amphipod family Crangonyctidae, a diverse family that comprises species inhabiting a variety of surface and subterranean habitats and for which several mitogenomes have been sequenced and annotated recently (e.g., Aunins *et al*. 2016; Benito *et al*. 2021).

## Materials and methods

We generated new mitogenomes recently for the following crangonyctid species: *Stygobromus pizzinii*, *S. tenuis potomacus*, *Bactrurus brachycaudus*, *Stygobromus allegheniensis*, and *Crangonyx forbesi* (Benito *et al*. 2021). We then downloaded from GenBank the annotated mitogenomes of 30 additional amphipod taxa that occupy aquatic habitats, including groundwater and springs, and three isopods that served as outgroups for comparative analyses.

### Mitogenome analyses

Nucleotide composition, amino acid frequencies, and codon usage were calculated in PhyloSuite v1.1.15 (Zhang *et al*. 2020). The web-based program CREx (http://pacosy.informatik.uni-leipzig.de/crex, Bernt *et al*. 2007) was used to perform pair-wise comparison of the gene orders in the mitogenome to determine rearrangement events. CREx comparisons were based on common intervals, and it considers common rearrangement scenarios including transpositions, reversals, reverse transpositions, and tandem-duplication-random-losses (TDRLs). In addition, phylograms and gene orders were visualized in iTOL (https://itol.embl.de/, Letunic and Bork 2021) using files exported from PhyloSuite. AT and GC skew of entire mitogenomes and individual loci were calculated in PhyloSuite using the formulae: AT-skew = (A - T)/(A + T) and GC-skew = (G-C)/(G + C). Welch two sample t-tests were performed between the surface and subterranean amphipods for different mitogenomic features, including mitogenome length, AT content, AT and GC skew, and rRNA length using R (R Core Team 2021). Visualization of AT-skew, GC-skew, AT-content, and amino acid frequencies were generated in R.

### Phylogenetic inference

The amino acid sequences of 13 PCGs of the five new mitogenomes (Benito *et al*. 2021), 30 previously published amphipod mitogenomes, and three isopod mitogenomes were aligned using MAFFT version 7 (Katoh and Standley 2013). Poorly aligned regions were eliminated using Gblocks version 0.91b (Castresana 2000). The best partitioning strategy and best-fit evolutionary models for each partition were inferred using PartitionFinder v2.1.1 (Lanfear *et al*. 2012).

Phylogenetic relationships of the 35 amphipod mitogenomes and three isopod mitogenomes using the concatenated 13 PCG alignment were determined using Bayesian inference in MrBayes v3.2 (Ronquist *et al*. 2012). The analyses contained two parallel runs with four chains each and were conducted for 5,000,000 generations (sampling every 100 generations). After dropping the first 25% “burn in” trees to ensure stationarity after examination of log-likelihood values for each Bayesian run using Tracer v1.7 (Rambaut *et al*. 2018), the remaining 37,500 sampled trees were used to estimate the consensus tree and the associated Bayesian posterior probabilities. All analyses were conducted within PhyloSuite.

### Positive selection and selection pressure analyses of mitochondrial PCGs

We performed base-substitution analyses on entire mitogenomes as well as for each of the 13 PCGs individually to compare surface versus subterranean amphipod taxa. The non-synonymous to synonymous rate ratio (dN/dS or ω) for each PCG was estimated using the free-ratio model using the CodeML program implemented in PAML v4.8a (Xu and Yang 2013). The ω values were estimated for surface and subterranean species separately and visualized in R for comparison. To estimate the probability of positively selected sites in each PCG across all amphipod species, we implemented site models (M1 and M2, M8a and M8), where ω was allowed to vary among sites (Yang 2007). To further investigate if some lineages and sites in particular lineages have undergone positive selection, we conducted maximum likelihood analyses on all PCG using the branch model and branch-site model in EasyCodeML v1.21, a visual tool for analysis of selection using CodeML (Gao *et al*. 2019). For branch models, the ‘one-ratio’ model (model 0), and ‘two-ratios’ model were implemented in the combined dataset of 13 PCG as well as on each PCG to identify if selective pressure differed between an amphipod lineage of interest (foreground branch) and other amphipod lineages (background branches). A likelihood ratio test (LRT) was conducted for each PCG to test whether ω was homogenous across all lineages. In the branch model, the null hypothesis assumes that the average ω values of branches of interest (ωF) is equal to that of other branches (ωB), whereas the alternative hypothesis assumes the opposite ωF≠ωB. If the alternative hypothesis is supported and ω > 1, the foreground lineage is under positive selection. The branch-site model allows ω to differ among codon sites in a foreground lineage when compared to background lineages. We implemented the branch-site model to identify sites on specific lineages regulated by positive selection. Selected sites were considered positively selected only if they passed a Bayes Empirical Bayes (BEB) analysis with a posterior probability of >95%.

We performed selection pressure analyses on the concatenated 13 PCGs dataset aligned using the codon mode as well as on each PCG with the Bayesian topology (see Figure 4) as the guidance tree using several approaches available from the Datamonkey Webserver (Weaver *et al*. 2018). First, we implemented aBSREL (Adaptive Branch-Site Random Effects Likelihood), an improved version of the commonly used “branch-site” models, to test if positive selection has occurred on a proportion of branches (Smith *et al*. 2015). We implemented BUSTED (Branch-site Unrestricted Statistical Test for Episodic Diversification) to test for gene-wide (not site-specific) positive selection by querying if a gene has experienced positive selection in at least one site on at least one branch (Murrell *et al*. 2015). Finally, we implemented RELAX (Wertheim *et al*. 2015) to test whether the strength of selection has been relaxed or intensified along a specified set of test branches.

## Results and discussion

We compared the mitogenomes for 35 surface and groundwater amphipods, including seven recently sequenced mitogenomes of one spring-dwelling and six groundwater species in the family Crangonyctidae by Aunins *et al*. (2016) and Benito *et al*. (2021), to determine whether subterranean species show evidence of adaptive evolution in subterranean habitats. Our study included a phylogenetic analysis, examined whether features of mitogenomes, such as base composition, codon usage, and gene order, differed in relation to dominant habitat (surface vs. subterranean), and conducted selection analyses to infer the evolutionary forces potentially shaping mitogenome evolution in amphipods, with an emphasis on crangonyctid species.

### Mitogenome length and content

Mitogenome sizes ranged from 14,113 to 18,424 bp for all amphipods and 14,661 to 15,469 bp for crangonyctid amphipods (Table 1). Mean mitogenome size of surface amphipods (15,770 ± 1206 bp; mean ± 1 standard deviation) was higher than that of the subterranean amphipods (14,716 ± 297 bp) (t-test: t = −3.31, df = 15.3, p-value = 0.005; Supplementary Figure S1). All crangonyctid amphipod mitogenomes possessed 13 PCGs, two rRNA genes, 22 tRNA genes, a control region, and intergenic spacers of varying number and lengths (Supplementary Figure S2, annotations of the genomes are presented in Supplementary Table S1) like other arthropods (Clary and Wolsetenholme 1985). The length of the crangonyctid mitogenomes is similar to lengths reported for other amphipod families including Allocrangonyctidae, Caprellidae, Eulimnogammaridae, Gammaridae, Hadziidae, Lysianassidae, Metacrangonyctidae, Micruropodidae, Pallaseidae, Pontogeneiidae, Talitridae. Variation in mitogenome length within Crangonyctidae appears to be related to length variation variation in the *nad5, rrnL*, and *rrnS* loci, which were all notably longer in the *C. forbesi* mitogenome.

**Table 1.**
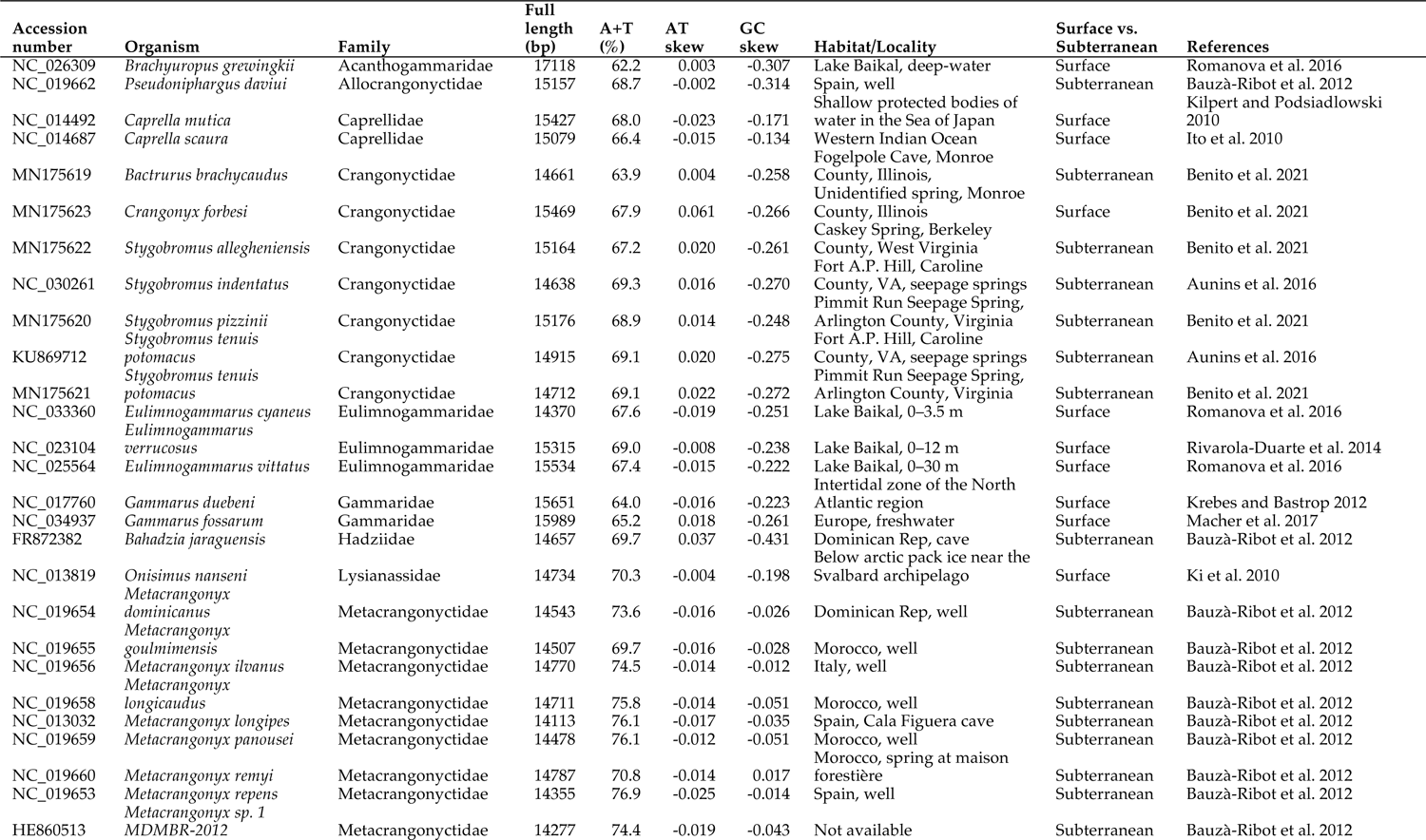
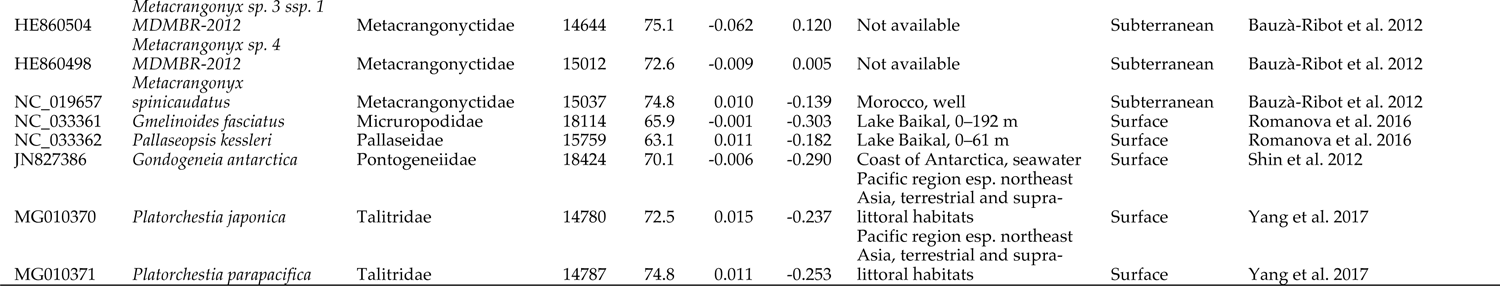
Summary of mitogenomic characteristics, location, and habitat of subterranean and surface amphipods included for comparative mitogenome analyses.

### Base composition and AT- and GC-skews

Mitogenome AT% in all amphipods ranged from 62.2 to 76.9% (Table). Mean AT% of the subterranean amphipods (71.8 ± 3.6%) was higher than that of the surface amphipods (67.6 ± 3.4%) (t-test: t = 3.49, df = 31.2, p-value = 0.001). Mean AT% of all 13 PCG of the subterranean amphipods was significantly higher than that of the surface amphipods (Supplementary Figure S3a). Variation in AT% across crangonyctid amphipod taxa ranged 63.9–69.3%, with a mean of 67.9 ± 1.93% (Table 1). Across loci, AT% ranged from a minimum of 60.0% at the *cox1* locus and a maximum of 75.5% at the *nad4l* locus (Figure 1A). Variation in AT% across all PCGs combined ranged from 61.9% (*B. brachycaudus*) to 69.0% (*S. indentatus*). Genes encoded on the negative strand had a slightly higher AT-content values than those on the positive strand. The *nad6* locus showed the greatest variation in AT-content across species. *Bactrurus brachycaudus* displayed the outlier lower AT% values for most of the PCG (Table 2, Figure 1A). Similarly, *Bactrurus brachycaudus* had the lowest AT content (63.9%) among the crangonyctid mitogenomes, while all other mitogenomes had comparatively typical AT content reported for other arthropods (Crozier and Crozier 1993; Dotson and Beard 2001). This could indicate that the evolution of the *B. brachycaudus* mitogenome has occurred under different evolutionary pressures (adaptive or non-adaptive) than other subterranean crangonyctids.

**Figure 1.**
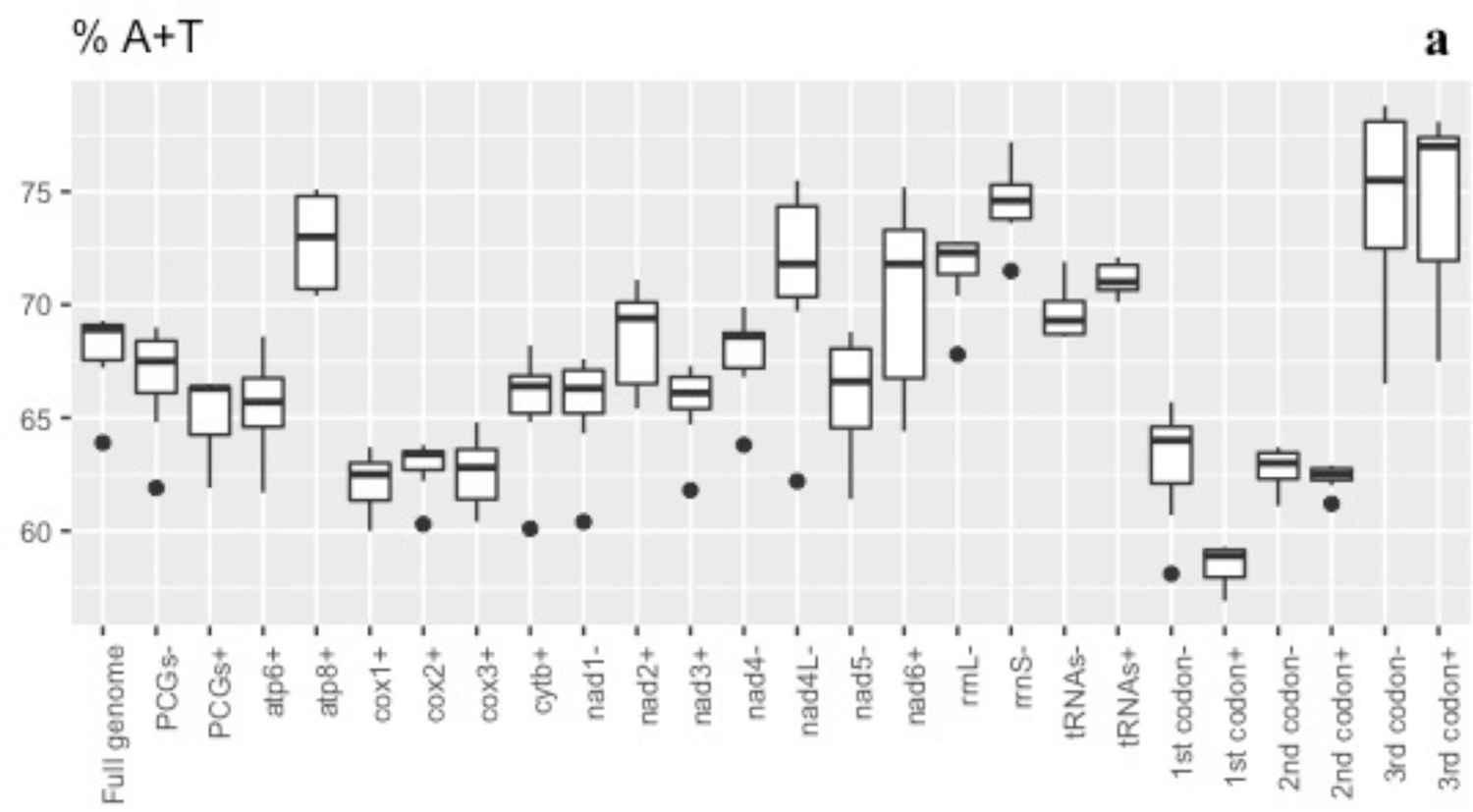
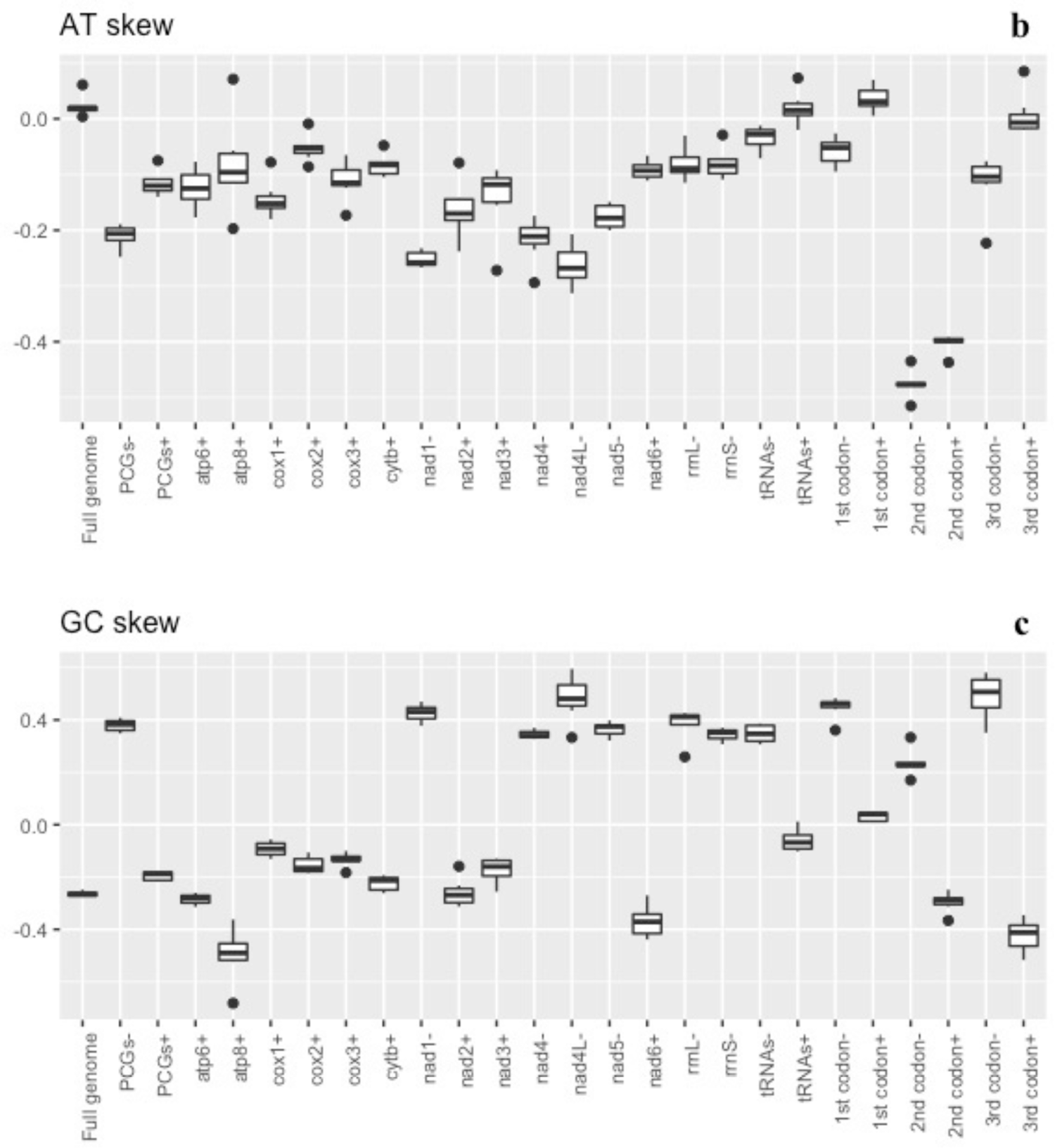
Crangonyctidae mitochondrial nucleotide composition. Box plots showing values of nucleotide composition (A + T percentage) **(a)**, AT-skew **(b)**, and GC-skew **(c)** across mitogenomes, protein coding genes (PCG), ribosomal (rRNA), and transfer ribosomal (tRNA). The same features are shown for each protein-coding gene and pooled by codon position and coding strand. Genes coded on the (-) strand are represented by a “-“ sign and genes coded on the (+) strand are represented by “+” sign at the end of the gene label.

**Table 2.**
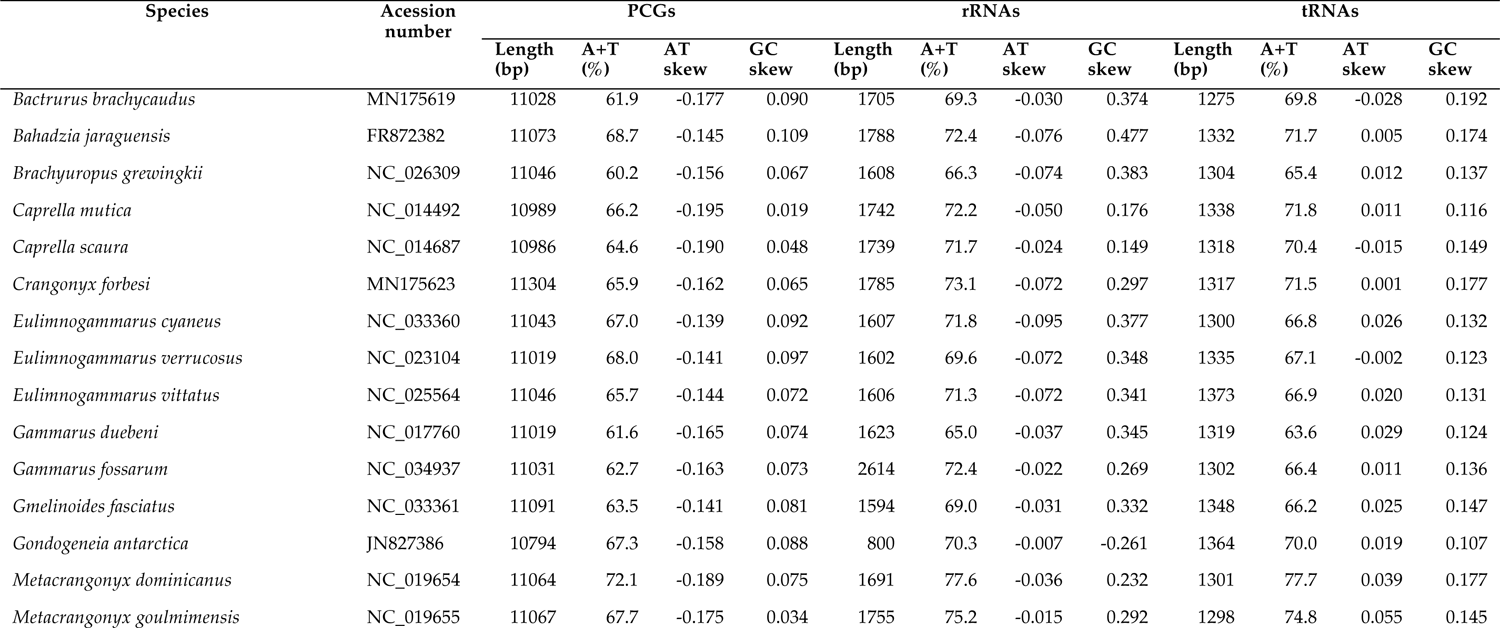
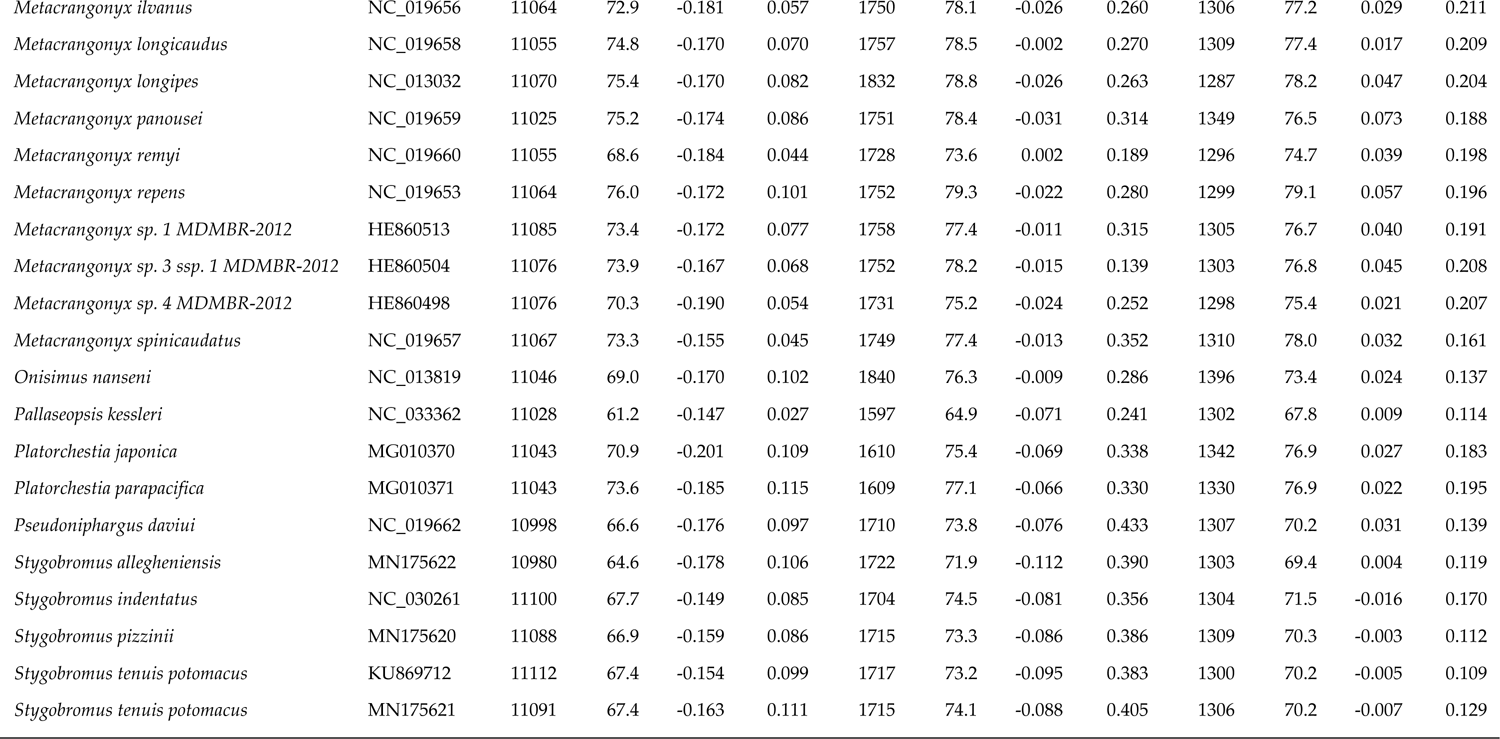
Comparison of mitogenomic characteristics of 35 amphipods discussed in this study

Mitogenome AT-skew in all amphipods ranged from −0.062 to −0.037. Mean AT-skew of the surface amphipods (0.001 ± 0.02) was positive and slightly higher than that of the subterranean amphipods (−0.004 ± 0.02) (t = −0.63101, df = 31.117, p-value = 0.5326). Mean AT-skew of four PCG (*cox1*, *cox2*, *nad2*, *nad3*) of surface amphipods was significantly higher than that of the subterranean amphipods, whereas the mean AT-skew of *nad4* of the subterranean amphipods was significantly higher than that of the surface amphipods (Supplementary Figure S3b). Among crangonyctid amphipods, mean AT-skew was 0.022 ± 0.02 (range 0.004 to 0.061), with all mitogenomes displaying positive skew. Mitogenome GC-skew ranged from −0.431-0.120. Mean GC-skew of the subterranean amphipods (−0.129 ± 0.15) was negative and higher than that of the surface amphipods (−0.236 ± 0.05) (t = 3.01, df = 24.3, p-value = 0.006). Mean GC-skew of seven PCG (*atp6*, *atp8*, *cox1*, *cox2*, *cox3, nad2*, *nad3*) of subterranean amphipods was significantly higher than that of the surface amphipods, whereas the mean GC-skew of *nad4* of the surface amphipods was significantly higher than that of the subterranean amphipods (Supplementary Figure S3c). Among crangonyctid amphipods, mean GC-skew was −0.264 ± 0.01 (range −0.275 to −0.248) with all mitogenomes displaying negative skew (Table 1). Strand asymmetry is commonly observed in mitogenomes (Reyes *et al*. 1998; Wei *et al*. 2010), however, at times it can hinder phylogenetic reconstruction and yield false phylogenetic artefacts especially when unrelated taxa display inverted skews (Hassanin *et al*. 2005; Zhang *et al*. 2019). *Bactrurus brachycaudus* exhibited the lowest AT skew among the crangonyctid mitogenomes (0.004), while *S. tenuis* had the lowest GC skew (−0.275). Crangonyctid amphipod mitogenomes exhibited positive GC-skew values for genes encoded on the (-) strand and negative GC-skew for genes encoded on the (+) strand (Figure 1C), whereas all PCGs exhibited negative AT-skew values (Figure 1B). Except the six loci (*nad1, nad4, nad4L, nad5, rrnL,* and *rrnS*) which were encoded on the (-) strand, most PCG had negative GC skews. Such strand bias is typical for most mitochondrial genomes in metazoan (Ki *et al*. 2010; Krebes and Bastrop 2012). This is consistent with the hypothesis that strand asymmetry is caused by spontaneous deamination of bases in the leading strand during replication (Reyes *et al*. 1998). All other mitogenomes had comparatively typical AT and GC skew values like other amphipod species (Pons *et al*. 2014; Romanova *et al*. 2016). The only outlier to this pattern was the positive GC skew value of tRNAs encoded on the (+) strand of B*. brachycaudus* (0.012). In general, crangonyctid mitogenomes exhibited relatively consistent skews.

Higher AT-content was observed at first (t = 3.80, df = 67.8, p-value < 0.001), second (t = 5.13, df = 67.9, p-value < 0.001), and third codon positions (t = 4.26, df = 60.7, p-value < 0.001) of PCGs on both the (+) and (-) strands in subterranean amphipods compared to surface amphipods (Supplementary Figure S3a). Among crangonyctid amphipods, a contrasting pattern was observed in the nucleotide composition per codon position (Figure 1A). AT-content was higher at third positions on both strands (74.5 ± 4.1% on (+) strand; 74.6 ± 4.4% on (-) strand) compared to first (58.5 ± 0.9% on (+) strand; 63.0 ± 2.7% on (-) strand) and second positions (62.4 ± 0.6% on (+) strand; 62.8 ± 0.9% on (-) strand). AT skew was near zero at the first codon position, whereas a T nucleotide-enrichment (about −0.4 AT skew value on average) in loci of both strands was observed at the second codon position. Intermediate negative AT skews was observed at the third codon position (Figure 1B). GC skew was positive for the first codon position, negative or close to zero for the second codon position and showed greater variation at third codon positions between loci on the positive and negative strands (Figure 1C).

### Rearrangements of mitochondrial genome

Conserved gene blocks in crangonyctid mitogenomes include (1) cox1*-*tRNA*-L2-cox2-*tRNA*-K,D-atp8-atp6-cox3-nad3-*tRNA*-A,S1,N,E,R,F-nad5* and (2) tRNA*-H-*nad4-nad4l and (3) *nad6-cytb-*tRNA*-S2* and (4) tRNA*-L1-rrnL* and (5) *rrnS-*tRNA*-I* and (6) tRNA-*Y,Q* (Figure 2B). The gene orders in subterranean species (genera *Stygobromus* and *Bactrurus*) are identical except for the transposition of tRNA-*G,W*. However, a few unique gene order arrangements were observed in the spring-dwelling *C. forbesi*. The gene order of *C. forbesi* differs from the four subterranean species in the locations of the conserved gene blocks (tRNA*-H-*nad4-*nad4l* and *nad6-cytb-* tRNA*-S2* and tRNA*-L1-rrnL* and *rrnS-*tRNA*-I* and tRNA*-Y,Q*), seven tRNAs (*P,T,M,V,G,C,* and *W*), and two protein-coding loci: *nad1* and *nad2*. Compared to the conserved mitogenome gene orders of other crangonyctid mitogenomes, another unique feature in the rearranged *C. forbesi* mitogenome was the presence of at least two long (∼50 and 70 bp) non-coding regions (Supplementary Table S1). The locations of rRNA genes in all crangonyctid mitogenomes are mostly similar compared to the pancrustacean ground pattern except for *C. forbesi* where the rRNA genes had altered positions (Figure 2A and 2B). Rearrangements in the mitogenome is common especially when it involves only tRNA-coding genes (Matsumoto *et al*. 2009). In case of ribosomal RNA genes or PCGs, rearrangements occur much less frequently, and they are commonly referred to as major rearrangements, as they might potentially affect the differential regulation of replication and transcription of mitogenomes (Rawlings *et al*. 2001).

**Figure 2.**
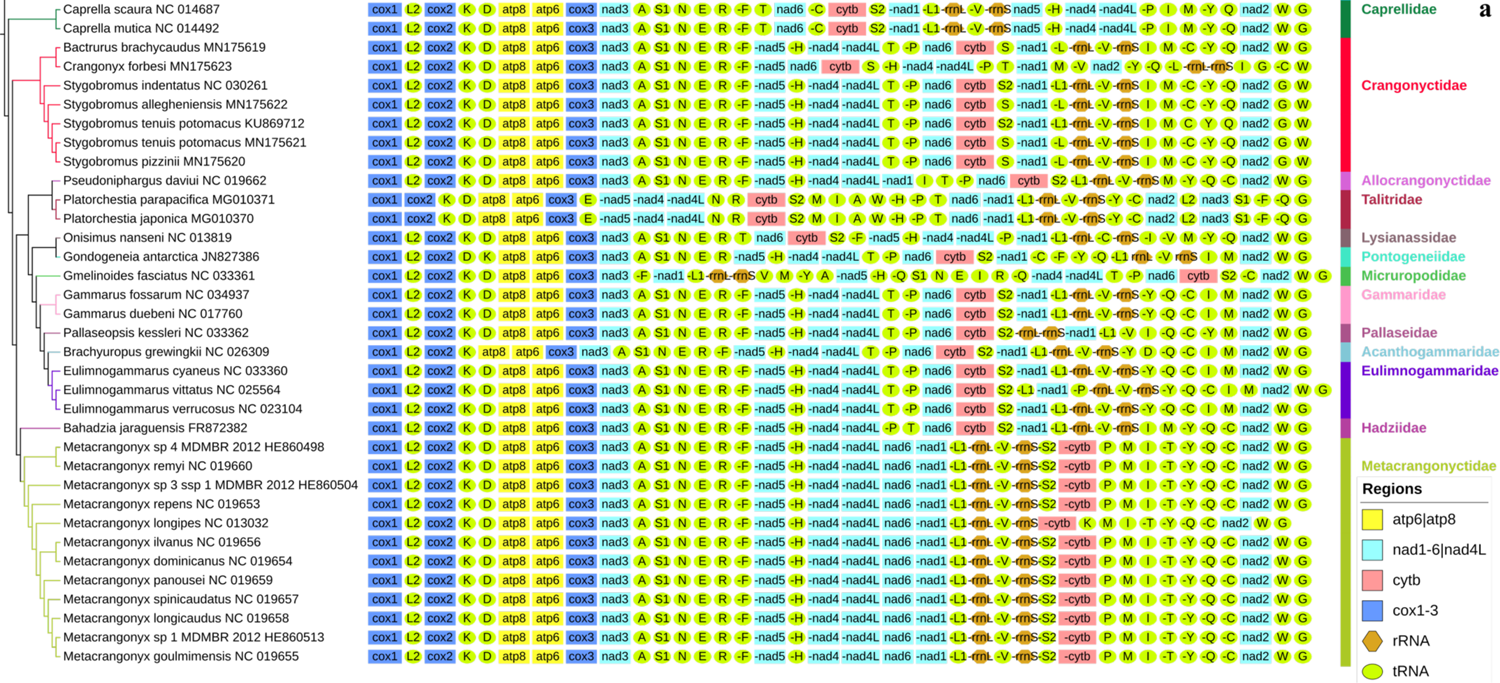
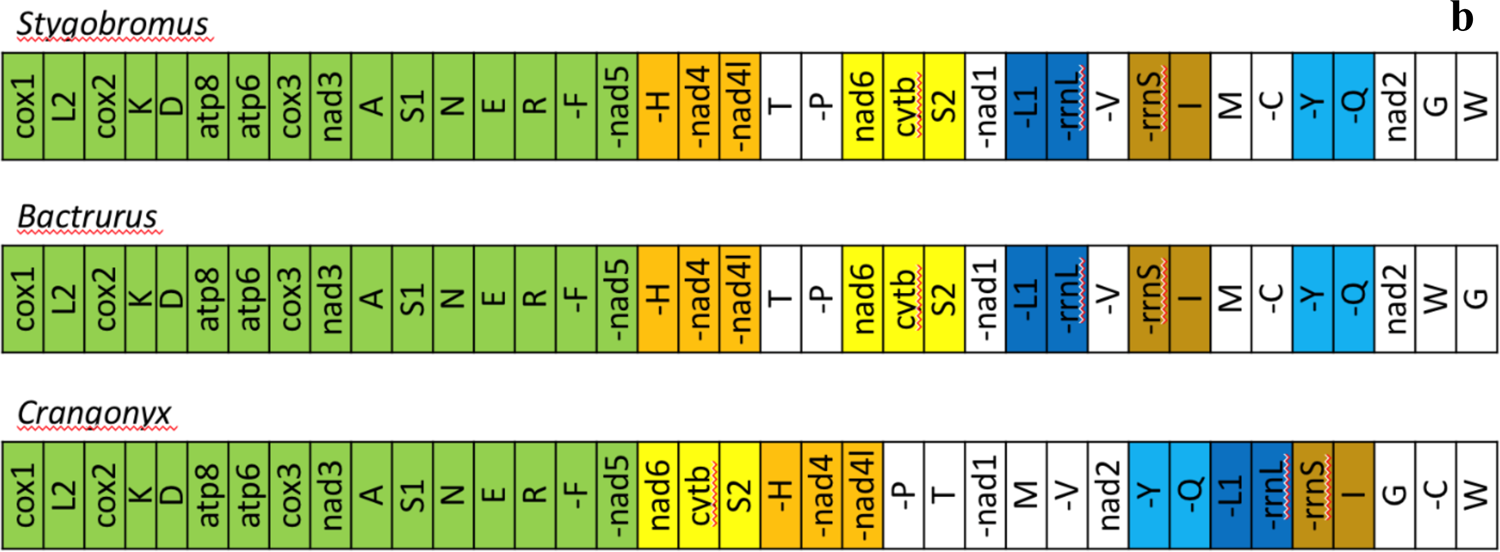
Mitochondrial phylogenomics and gene orders: (a) Bayesian phylogram inferred using amino acid sequences of all mitochondrial PCGs (left) and gene orders (right). Three isopod outgroups are not shown. GenBank accession numbers are included as suffix next to the species names; (b) gene orders of mitochondrial genomes in three genera of crangonyctid amphipods, including *Stygobromus*, *Bactrurus*, and *Crangonyx*. Conserved gene clusters are indicated by different colors.

CREx analysis indicated that transpositions and TDRL may have been responsible for the evolution of mitogenomes in crangonyctid amphipods. Two transpositions of tRNA-*R,N,S1,E* and two steps of TDRL from the ancestral pan-crustacean pattern were needed to generate the gene order observed in *Stygobromus* species. In addition to the same two transpositions, one TDRL, and a transposition within a second TDRL from the ancestral pattern were required to generate the gene order in *Bactrurus*. However, four different transpositions (tRNA-*N,S1* and tRNA-*T,P* and tRNA-*W,C* and gene block tRNA*-H-nad4-nad4L*-tRNA-*P,T*-*nad6-cytb*-tRNA-*S2*) and three steps of TDRL from the ancestral pattern were needed to generate the gene order observed in *C. forbesi* (Supplementary Figure S4).

Similar to *C. forbesi*, other surface amphipods including *Gmelinoides fasciatus* (Micruropodidae) and *Onisimus nanseni* (Lysianassidae) exhibited a highly rearranged gene order. Other surface amphipods that exhibited a moderate to highly rearranged gene order include *Gondogeneia antarctica* (Pontogeneiidae), *Platorchestia parapacifica* and *P. japonica* (Talitridae), *Pallaseopsis kessleri* (Pallaseidae), and the two basal amphipods *Caprella scaura* and *C. mutica* (Caprellidae). Interestingly, a subterranean amphipod *Pseudoniphargus daviui* (Allocrangonyctidae) also exhibited a moderate rearranged gene order. The stark contrast between the highly conserved gene order in most subterranean amphipods and the highly volatile gene order in many of the surface amphipods may support the hypothesis that evolution of mitogenomic architecture could be highly discontinuous. A prolong period of inactivity in gene order and content could have been dictated by a rearrangement event resulting in a destabilized mitogenome, which is much more likely to undergo an exponentially accelerated rate of mitogenomic rearrangements (Zou *et al*. 2017). Thus, it is appealing to examining mitogenomes of surface amphipod families represented by just a single taxon in our dataset.

### Codon usage and amino acid frequencies

In addition to the regular start codons (ATA and ATG) and uncommon start codons (ATT, ATC, TTG, and GTG), surface amphipods, particularly *Caprella scaura*, possessed one rare start codon CTG, whereas subterranean amphipods possessed three rare start codons including CTG, TTT, and AAT to initiate the mitochondrial PCGs (Supplementary Table S2). Codon usage analysis of the five crangonyctid amphipods mitogenomes identified the existence of all codon types typical for any invertebrate mitogenome. In addition to the regular start codons (ATA and ATG), uncommon start codons (ATT, ATC, TTG, and GTG) were also present to initiate the mitochondrial PCG. Such unusual start codons have been reported previously in other arthropods (Lessinger *et al*. 2000; Boore *et al*. 2005). A few PCG in the crangonyctid mitogenomes possessed truncated or incomplete stop codons (TA- and T--) that have been described in other crustaceans (Supplementary Table S1). These are presumably completed after a post-transcriptional polyadenylation (Ojala *et al*. 1981; Castresana *et al*. 1998; Nardi *et al*. 2001).

Among the crangonyctid mitogenomes, the most frequently used codons are TTA (Leu2; 5.64% to 8.49%) and TTT (Phe; 5.94% to 6.78%). Other frequently used codons include ATT (Ile; 4.92% to 6.85%) and ATA (Met; 4.13% to 5.34%) (Supplementary Table S3). These four codons are also among the most abundant in non-crangonyctid amphipods included in this study. This bias towards the AT-rich codons is quite typical for arthropods (Wilson *et al*. 2000). Among crangonyctid amphipod mitogenomes, relative synonymous codon usage (RSCU) values, which is the measure of the extent that synonymous codons depart from random usage, showed a high prevalence of A or T nucleotides at third codon positions (Figure 3). This trend was also observed in other subterranean and surface amphipods. This positive correlation between RSCU and AT content at third codon positions has been reported in mitochondrial genomes of the abalone and oyster (Ren *et al*. 2010; Xin *et al*. 2011).

**Figure 3.**
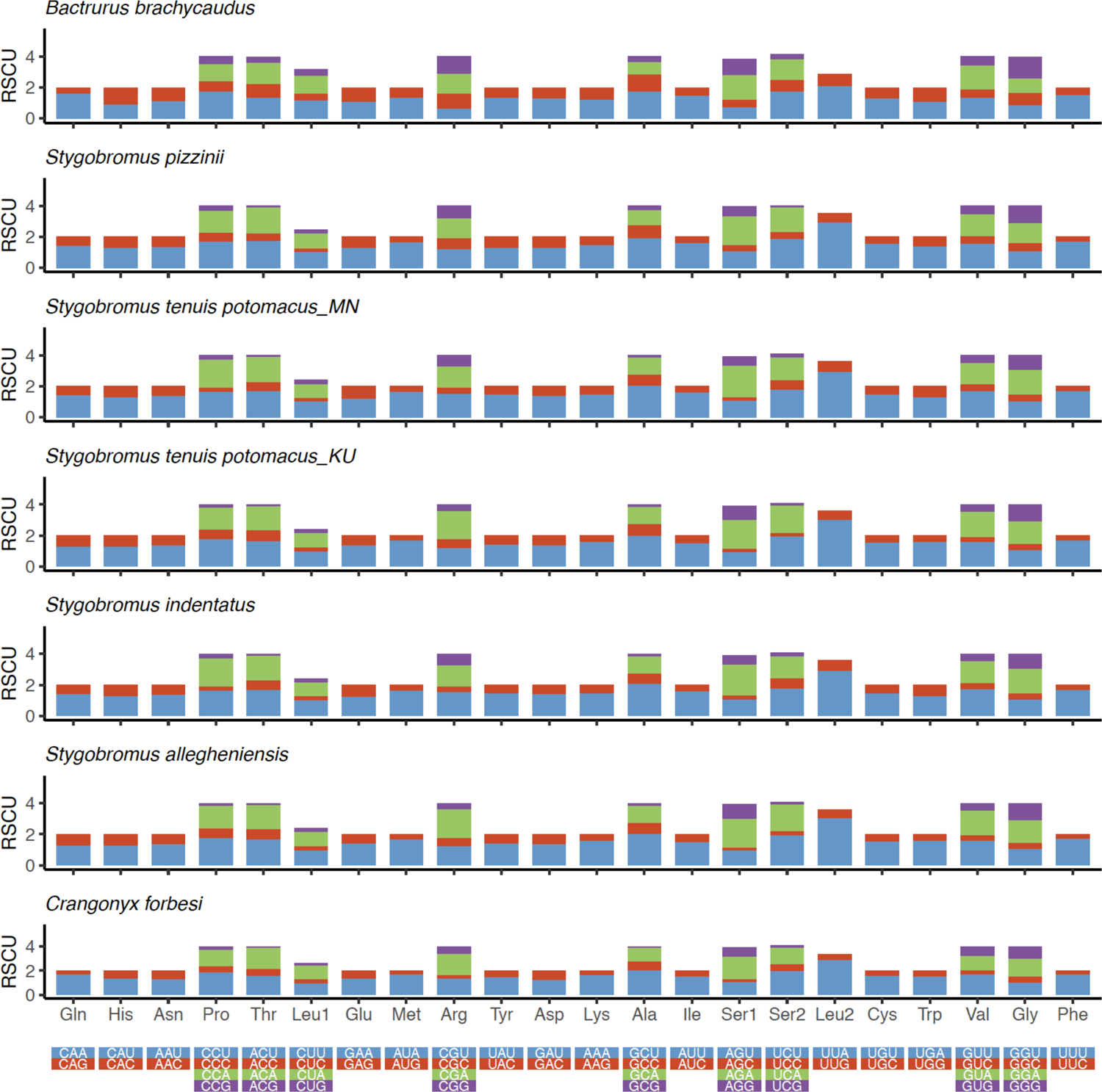
The relative synonymous codon usage (RSCU) of the mitogenome of all crangonyctid amphipods. The RSCU value are provided on the Y-axis and the codon families are provided on the X-axis.

In PCG, the second copy of leucine (8.86–10.01%) and cysteine (0.95–1.17%) are the most and the least used amino acids, respectively. Amino acid frequency analysis of both surface and subterranean amphipods indicated that five amino acids (leucine, phenylalanine, isoleucine, methonine, and valine) account for more than half of the total amino acid composition and exhibited greater variation among species (Supplementary Figure S5; Supplementary Table S4).

### Transfer RNA genes

All 22 tRNA genes were identified in the mitogenomes of crangonyctid amphipods. However, the locations of tRNA genes are highly variable among these mitogenomes, and they also displayed altered positions relative to the pancrustacean ground pattern (Figure 2; Supplementary Figure S3). The secondary structures of all mitogenome-encoded tRNAs belonging to crangonyctid amphipods were predicted and ranged in length from 50 to 66 bp. Most of the tRNAs displayed the regular clover-leaf structures, however, a few displayed aberrant structures. The tRNA-Ser1 (UCU) lacked the DHU arm in all crangonyctid species. Similarly, the tRNA-Ser2 (UGA) lacked the DHU arm in all crangonyctid species except *S. allegheniensis* where tRNA-Ser2 (UGA) possessed the DHU arm. The DHU arm was also missing in the tRNA-Cys and tRNA-Arg of *B. brachycaudus* and tRNA-Arg of *C. forbesi*. The tRNA-Gln lacked the TψC arm in all crangonyctid species except *C. forbesi* where tRNA-Gln possessed the TψC arm. In addition to lacking the TψC arm, tRNA-Gln of *B. brachycaudus* lacked the acceptor stem as well (Supplementary Figure S6). The presence of aberrant structures in tRNAs have been observed in several other crustaceans and invertebrates (Okimoto *et al*. 1990; Bauza-Ribot *et al*. 2009; Ito *et al*. 2010; Lin *et al*. 2012), which may be the result of replication slippage (Macey *et al*. 1997) or selection towards minimization of the mitogenome (Yamazaki *et al*. 1997).

### Ribosomal RNA genes

The length of *rrnL* genes in all amphipods ranged 604–1,137 bp and that of *rrnS* genes ranged 196–1,631 bp. *rrnL* length of the subterranean amphipods (1,055 ± 26 bp) was higher than that of the surface amphipods (971 ± 108 bp) (t-test: t = −2.94, df = 15.2, p-value = 0.001). *rrnS* length of the surface amphipods (694 ± 290 bp) was slightly higher than that of the subterranean amphipods (684 ± 16 bp) (t-test: t = 0.13, df = 14.1, p-value = 0.896). The length of *rrnL* genes in crangonyctid amphipods ranged 1,034–1,090 bp and that of *rrnS* genes ranged 671–695 bp. The length of rRNA genes in crangonyctid amphipods is similar to that of other amphipod mitogenomes except *C. forbesi*, which had long overhangs (∼50 bp and ∼25 bp) on the 5’ end of the *rrnL* and *rrnS* genes, respectively. AT content ranged 67.8–72.8% in the *rrnL* genes and 71.5–77.2% in the *rrnS* genes of crangonyctid species, respectively. GC-skew values for rRNA genes were positive (0.259 to 0.426) and comparable to that of PCGs encoded on the (-) strand (Supplementary Table S5).

### Control region and intergenic spacers

In the mitogenome of *S. pizzinii* the putative control region (CR) was identified as a 1,021 bp sequence between the *rrnS* gene and the *trnl-trnM-trnC-trnY-trnQ-nad2* gene cluster. Similarly, CR was observed in the other crangonyctid mitogenomes, including *S. tenuis* (556 bp), *S. allegheniensis* (991 bp), *B. brachycaudus* (531 bp), *S. indentatus* (535 bp), and *S. tenuis potomacus* (773 bp). The CR was similarly located between the *rrnS* and *nad2* genes in some of the other mitogenomes of non-crangonyctid amphipods, including *G. duebeni (*Krebes and Bastrop 2012), *O. nanseni* (Ki *et al*. 2010), *G. antarctica* (Shin *et al*. 2012), *P. daviui* (Bauzà-Ribot *et al*. 2012), and for the pancrustacean ground pattern. However, the adjacent tRNA genes were often different. In *G. fasciatus,* the CR region was located between the *rrnS* and *nad5* genes (Romanova *et al*. 2016). In contrast, a control region 843 bp was observed in *C. forbesi* which is located between the *nad1* and *trnM-trnV-nad2* gene cluster and separated by few intergenic spacers was identified as the CR (Supplementary Figure S1; Supplementary Table S1). The only other surface amphipod that had a similar CR location to *C. forbesi* was *P. kessleri* with the CR located between *nad1* and *nad2* genes, although the adjacent tRNA genes were different (Romanova *et al*. 2016). Thus, the variable location of the CR in *C. forbesi* is in concordance with few other surface amphipods, while the subterranean amphipods mostly followed the universal pattern between *rrnS* and *nad2* genes.

The non-coding regions or intergenic spacers identified in the crangonyctid mitogenomes varied in number and length. The number of intergenic spacers ranged from 7 to 17 and their lengths ranged from 1 to 99 bp (mean 13.0 bp ± 18.6). Two crangonyctid mitogenomes (*S. allegheniensis* and *C. forbesi*) possessed the largest intergenic spacers (a total of 220 bp and 249 bp, respectively; Supplementary Table S1). Among the non-crangonyctid amphipods, *G. fasciatus* and *G. antarctica* possessed relatively large non-coding intergenic spacers (a total of 3,863 bp and 4,354 bp, respectively; Shin *et al*. 2012; Romanova *et al*. 2016).

### Phylogenetic inference

The phylogenetic analyses of the 13 concatenated PCG from 35 amphipod species using Bayesian Inference (BI) resulted in a well-supported phylogeny, with the crangonyctid species forming a well-supported monophyletic group (Figure 4). Within Crangonyctidae, *Stygobromus* species formed a monophyletic group sister to *Bactrurus* + *Crangonyx*; however, few crangonyctid taxa were included in our analysis. A previous study based on the *cox1* gene found that *Stygobromus* was not monophyletic, but several relationships had low support (Niemiller *et al*. 2018). Likewise, *Stygobromus* as recovered as polyphyletic in a multilocus concatenated phylogenetic analysis of the Crangonyctidae by Copilas-Ciocianu et al. (2019). In addition, several well-supported clades were recovered within Crangonyctidae but relationships among these clades had low support. Other past studies have not supported monophyly of the widespread genera (i.e., *Crangonyx*, *Stygobromus*, and *Synurella*) in the family based on either morphological (Koenemann and Holsinger 2001) or molecular data (Kornobis et al. 2011, 2012). A comprehensive phylogenomic study with robust taxonomic sampling is greatly needed to better elucidate evolutionary relationships and test biogeographic and ecological hypotheses regarding the origin and diversification of this diverse family of subterranean and surface-dwelling amphipods.

### Selection in PCGs

Most of the energy required for active movement to escape predation and meet energy demands is supplied by the mitochondrial electron transport chain (Shen *et al*. 2009, 2010). Mitochondrial genes encode for all of the protein complexes related to oxidative phosphorylation except for succinate dehydrogenase (complex II) (Scheffler 1998; Carroll *et al*. 2009; McKenzie *et al*. 2009). Because of their importance in oxidative phosphorylation during cellular respiration, it is unsurprising that many studies have shown evidence of purifying (negative) selection in mitochondrial PCG (Meiklejohn *et al*. 2007; Hao *et al*. 2017; Sun *et al*. 2020). While we found strong evidence for purifying selection in amphipod mitochondrial PCG in our selection analyses, we also found signatures of positive selection in a few of the mitochondrial PCGs in the surface amphipods.

The one-ratio model (model 0) analyses conducted for all 13 PCG revealed that the *ω* values for each gene ranged from 0.021 to 0.130 and were significantly less than 1 (Table 3). Using a free-ratio model (model 1; Shen *et al*. 2009), we calculated the *ω* values for the 13 PCG for the terminal branches to estimate the strength of selection between different primary habitats (i.e., subterranean vs. surface). The *cox2* gene significantly differed in *ω* values between the amphipods of the two habitat types (*p* = 0.020), with higher *ω* values for the surface amphipods. Similarly, *cox1* and *cox3* genes also exhibited a similar trend (*p* = 0.095 and *p* = 0.057, respectively) (Figure 5). This could be because the rate at which slightly deleterious mutations (ω) responsible for the mitochondrial gene evolution accumulates comparatively faster in *cox* gene family of the surface lineages. However, this result is quite contradictory to previous studies showing higher functional constraint and conserved pattern in the genes coding for *cox* proteins than in other mitochondrial genes (Meiklejohn *et al*. 2007; Zapelloni *et al*. 2021).

**Figure 4.**
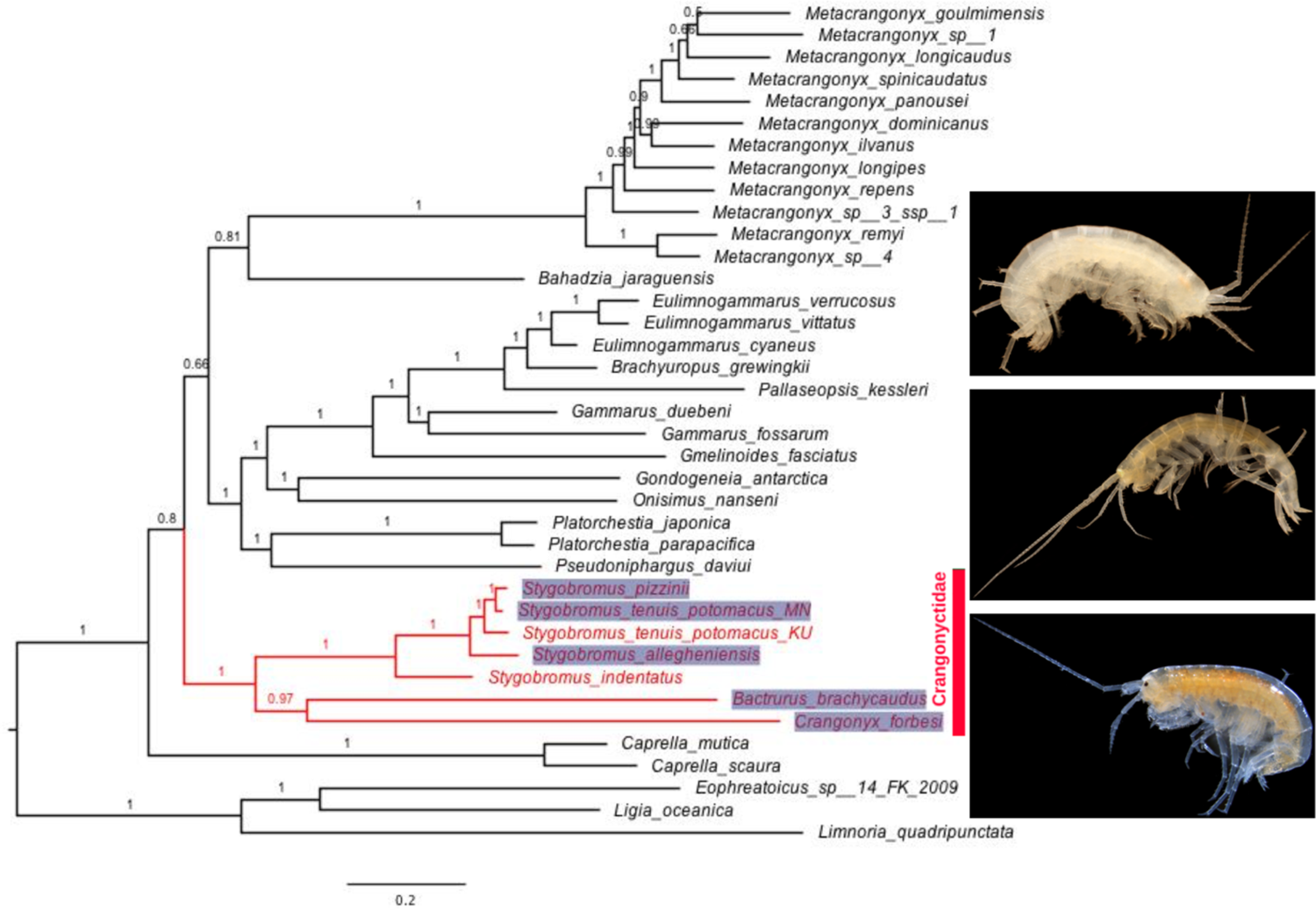
Bayesian phylogeny of aligned protein-coding loci (3,607 amino acids) for five new amphipod mitogenomes (*Stygobromus allegheniensis, S. pizzinii, S. tenuis potomacus, Bactrurus brachycaudus,* and *Crangonyx forbesi*) in addition to 30 additional amphipod mitogenomes available on Genbank. The three isopods *Ligia oceanica, Limnoria quadripunctata,* and *Eophreatoicus sp.14 FK-2009* are included as an outgroup to root the phylogeny. New mitogenomes generated in this study are highlighted. GenBank accession numbers are included as suffix next to the species names. Values at nodes represent posterior probabilities.

**Figure 5.**
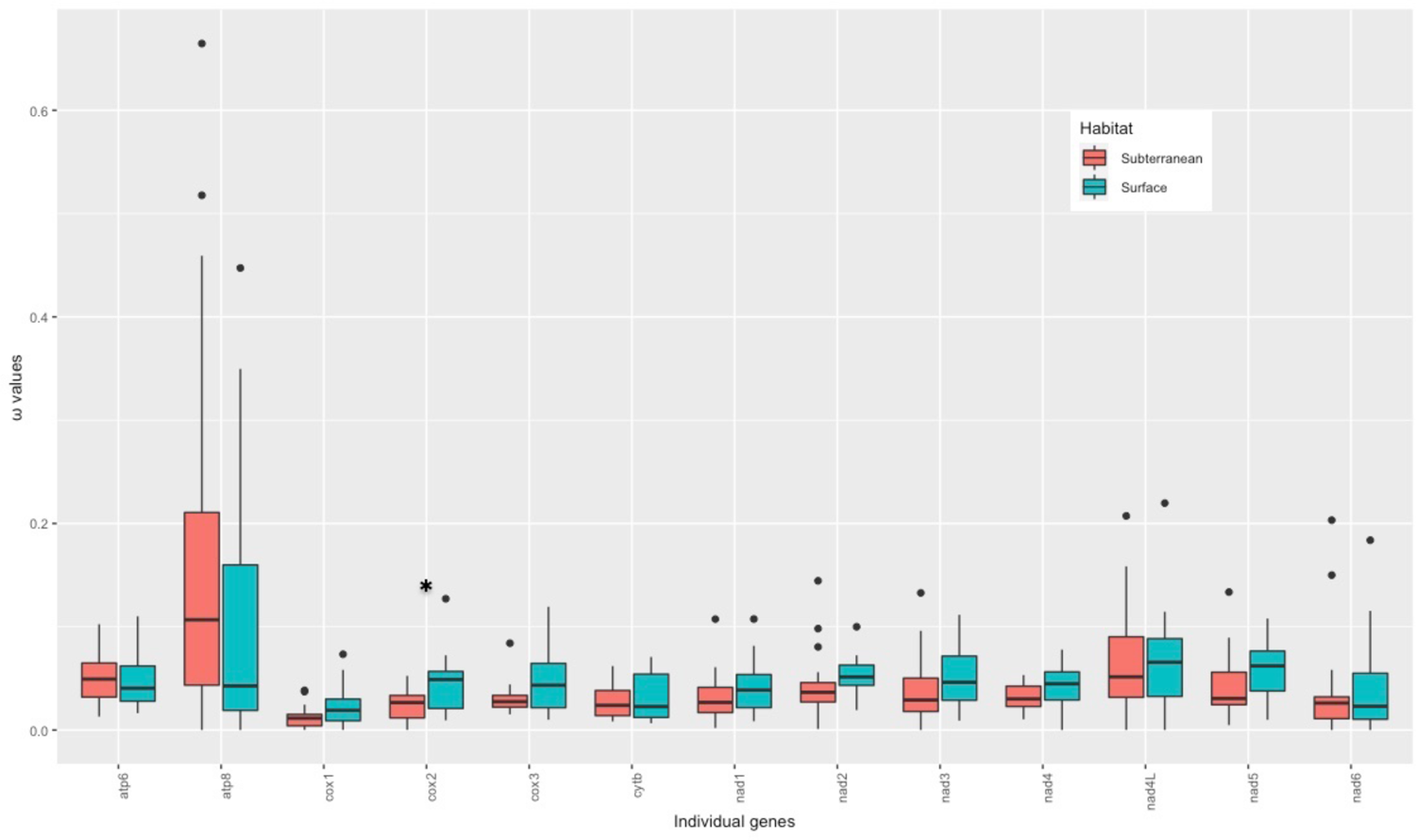
Ratio of non-synonymous to synonymous substitutions (ω) in the 13 PCGs of subterranean (coral color) and surface (cyan color) amphipods based on the free-ratio model. Boxes include 50% of values; ω is not significantly different between subterranean and surface amphipods for any gene except *cox2*\*. P values: *atp6* = 0.52; *atp8* = 0.27; *cox1* = 0.10; *cox2* = 0.02; *cox3* = 0.06; *cytb* = 0.48; *nad1* = 0.28; *nad2* = 0.33; *nad3* = 0.34; *nad4* = 0.18; *nad4l* = 0.93; *nad5* = 0.11; *nad6* = 0.65.

**Table 3.** Likelihood ratio tests of selective pressures (ω ratio) on mitochondrial PCGs between subterranean and surface amphipods. The terminal branches of surface amphipods are assigned as foreground branches and the terminal branches of subterranean amphipods are assigned as foreground branches. PCGs with significant LRT P-value < 0.05 are highlighted in red color

| Model | np | LnL | Estimates of parameters |  |  | Model compared | LRT P-value | Omega for Foreground Branch | Gene |
| --- | --- | --- | --- | --- | --- | --- | --- | --- | --- |
| Two ratio Model 2 | 70 | -15282.77 | $\omega$: | $\omega_0=0.05659$ | $\omega_1=0.04105$ | Model 0 vs. Two ratio Model 2 | 0.0184 | $\omega_1=0.04105$ | atp6 |
| Model 0 | 69 | -15285.55 | $\omega=$ | 0.05174 | | | | | |
| Two ratio Model 2 | 70 | -3926.02 | $\omega$: | $\omega_0=0.12582$ | $\omega_1=0.14171$ | Model 0 vs. Two ratio Model 2 | 0.7207 | $\omega_1=0.14171$ | atp8 |
| Model 0 | 69 | -3926.08 | $\omega=$ | 0.13019 | | | | | |
| Two ratio Model 2 | 70 | -24614.85 | $\omega$: | $\omega_0=0.01988$ | $\omega_1=0.02323$ | Model 0 vs. Two ratio Model 2 | 0.1009 | $\omega_1=0.02323$ | cox1 |
| Model 0 | 69 | -24616.19 | $\omega=$ | 0.02099 | | | | | |
| Two ratio Model 2 | 70 | -12959.59 | $\omega$: | $\omega_0=0.02831$ | $\omega_1=0.03729$ | Model 0 vs. Two ratio Model 2 | 0.0720 | $\omega_1=0.03729$ | cox2 |
| Model 0 | 69 | -12961.21 | $\omega=$ | 0.03033 | | | | | |
| Two ratio Model 2 | 70 | -16206.66 | $\omega$: | $\omega_0=0.04062$ | $\omega_1=0.04744$ | Model 0 vs. Two ratio Model 2 | 0.1739 | $\omega_1=0.04744$ | cox3 |
| Model 0 | 69 | -16207.58 | $\omega=$ | 0.04274 | | | | | |
| Two ratio Model 2 | 70 | -23399.90 | $\omega$: | $\omega_0=0.03742$ | $\omega_1=0.02615$ | Model 0 vs. Two ratio Model 2 | 0.0024 | $\omega_1=0.02615$ | cytb |
| Model 0 | 69 | -23404.50 | $\omega=$ | 0.03337 | | | | | |
| Two ratio Model 2 | 70 | -19418.37 | $\omega$: | $\omega_0=0.03385$ | $\omega_1=0.02885$ | Model 0 vs. Two ratio Model 2 | 0.2450 | $\omega_1=0.02885$ | nad1 |
| Model 0 | 69 | -19419.05 | $\omega=$ | 0.03239 | | | | | |
| Two ratio Model 2 | 70 | -19836.87 | $\omega$: | $\omega_0=0.04288$ | $\omega_1=0.04833$ | Model 0 vs. Two ratio Model 2 | 0.3826 | $\omega_1=0.04833$ | nad2 |
| Model 0 | 69 | -19837.25 | $\omega=$ | 0.04445 | | | | | |
| Two ratio Model 2 | 70 | -7403.62 | $\omega$: | $\omega_0=0.04379$ | $\omega_1=0.05134$ | Model 0 vs. Two ratio Model 2 | 0.4299 | $\omega_1=0.05134$ | nad3 |
| Model 0 | 69 | -7403.93 | $\omega=$ | 0.04560 | | | | | |
| Two ratio Model 2 | 70 | -29490.78 | $\omega$: | $\omega_0=0.04233$ | $\omega_1=0.04466$ | Model 0 vs. Two ratio Model 2 | 0.6063 | $\omega_1=0.04466$ | nad4 |
| Model 0 | 69 | -29490.91 | $\omega=$ | 0.04296 | | | | | |
| Two ratio Model 2 | 70 | -6866.17 | $\omega$: | $\omega_0=0.03397$ | $\omega_1=0.04991$ | Model 0 vs. Two ratio Model 2 | 0.1902 | $\omega_1=0.04991$ | nad4l |
| Model 0 | 69 | -6867.03 | $\omega=$ | 0.03788 | | | | | |
| Two ratio Model 2 | 70 | -42647.82 | $\omega$: | $\omega_0=0.06189$ | $\omega_1=0.05843$ | Model 0 vs. Two ratio Model 2 | 0.5037 | $\omega_1=0.05843$ | nad5 |
| Model 0 | 69 | -42648.04 | $\omega=$ | 0.06077 | | | | | |
| Two ratio Model 2 | 70 | -11163.67 | $\omega$: | $\omega_0=0.04176$ | $\omega_1=0.04888$ | Model 0 vs. Two ratio Model 2 | 0.4948 | $\omega_1=0.04888$ | *nad6* |
| Model 0 | 69 | -11163.91 | $\omega=$ | 0.04373 | | | | | |

To test if specific branches have undergone variable selective pressures, especially those amphipod branches adapted to surface habitats, we employed the two-ratio branch model (model 2). We evaluated the selective pressures acting on surface amphipods (foreground, *ω*1) and subterranean amphipods (background, *ω*0). LRT tests showed that the two-ratio model fits were significantly better than the one-ratio model for two genes: *atp6* (*p* = 0.018) and *cytb* (*p* = 0.002) (Table 3), indicating a divergence in selective pressure between surface and subterranean amphipods. In addition, a similar trend was observed for the *cox2* gene (*p* = 0.072). We then tried the same two-ratio model to estimate selection acting on each subterranean and surface amphipod lineage to further examine the difference between them. When the ω values for the individual PCG were compared between each amphipod lineage and the other 34 amphipod taxa, several genes in surface amphipod mitogenomes were found to be undergoing positive selection (ω1 > ω0; Figure 6; Supplementary Table S6). This suggests that many surface amphipods have experienced directional selection in their mitochondrial genes due to high energy demands and was accordance to the previous studies of insect orders (Yang *et al*. 2014; Mitterboeck *et al*. 2017; Li *et al*. 2018). In contrast, several genes in subterranean amphipod mitogenomes were undergoing purifying selection (ω1 < ω0). Surprisingly, a few genes in subterranean taxa displayed positive selection (ω1 > ω0; Figure 6; Supplementary Table S6). This indicated that the subterranean amphipods have undergone stronger evolutionary constraints to remove deleterious mutations to maintain efficient energy metabolism (Shen *et al*. 2009). Overall, this proves that molecular evolution of mitochondrial genes is correlated to the changes in the energy requirements.

**Figure 6.**
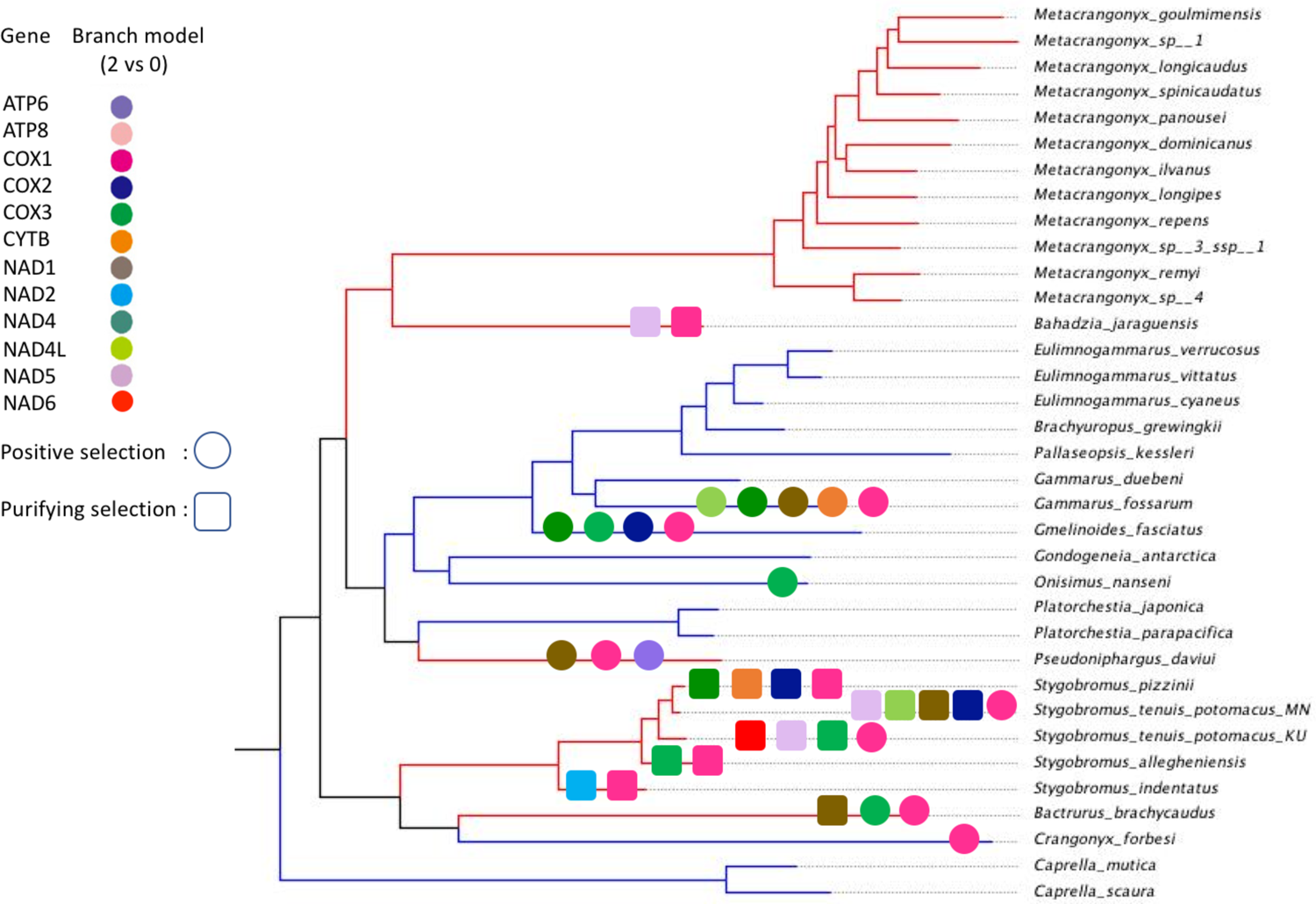
Results of selective pressure analysis of mitochondrial PCGs with LRT P-value < 0.05 in subterranean and surface-dwelling lineages of amphipods based on branch 2 vs. 0 model. Different colored shapes represent different mitochondrial genes. Squares represent purifying selection and circles represent positive selection. Surface amphipod branches are colored blue and subterranean amphipod branches are colored red.

To test if individual gene codons were subject to positive selection, we implemented two pairs of site models (M1a vs. M2a and M8a vs. M8). The M8 model identified one positively selected site on the *atp8* gene (37 N; *p* = 0) and one positively selected site on the *nad5* gene (482 Q; *p* = 0). Similarly, The M2a model identified two positively selected sites (37 N & 31 S; *p* = 0.0194) on the *atp8* gene (Table 4).

**Table 4.** Evidence of positive selection on the mitochondrial PCGs of subterranean and surface dwelling amphipods based on site models

| Model | np | Ln L | Estimates of parameters |  |  |  | Model compared | LRT P-value | Positive sites | Gene |
| --- | --- | --- | --- | --- | --- | --- | --- | --- | --- | --- |
| M2a | 72 | -4021.125381 | p: | 0.32097 | 0.56570 | 0.11334 | M1a vs. M2a | 0.019437929 | 31 S 0.977*, 37 N 0.997** | *atp8* |
| | | | $\omega$: | 0.09324 | 1.00000 | 2.44333 | | | | |
| M1a | 70 | -4025.065910 | p: | 0.37651 | 0.62349 |  |  |  |  |  |
| | | | $\omega$: | 0.09961 | 1.00000 | | | | | |
| M8 | 72 | -3950.635672 | p0=0.85323<br>(p1= 0.14677) | p=0.54776<br>$\omega= 1.00000$ | q=3.55619 | | M8a vs.M8 | 0.000013424 | 37 N 0.875 | |
| M8a | 71 | -3941.161040 | p0=0.97880<br>(p1= 0.02120) | p=0.73512<br>$\omega= 1.00000$ | q=3.74996 | | | | | |
| M2a | 72 | -42238.654620 | p: | 0.85660 | 0.08524 | 0.05815 | M1a vs. M2a | 1.000000000 | 482 Q 0.524 | *nad5* |
| | | | $\omega$: | 0.07211 | 1.00000 | 1.00000 | | | | |
| M1a | 70 | -42238.654620 | p: | 0.85660 | 0.14340 |  |  |  |  |  |
| | | | $\omega$: | 0.07211 | 1.00000 | | | | | |
| M8 | 72 | -40545.492521 | p0=0.98460<br>(p1= 0.01540) | p=0.53211<br>$\omega= 1.00000$ | q=7.22545 | | M8a vs.M8 | 0.000000000 | 482 Q 0.856 | |
| M8a | 71 | -40454.428911 | p0=0.99711<br>(p1= 0.00289) | p=0.50047<br>$\omega= 1.00000$ | q=6.79227 | | | | | |

Similar to the mitochondrial genes of the flying grasshoppers that have evolved to adapt to increased energy demands to maintain the flight capacity (Li *et al*. 2018), the mitochondrial genes of surface amphipods may have evolved mechanisms to meet increased energy demands due to predation, dispersal, etc. Although surface amphipods appear to be evolving under selective pressures different from those of the subterranean taxa and their mitochondrial genes have accumulated more nonsynonymous than synonymous mutations compared to subterranean taxa, the branch model tests did not clearly support positive selection on these branches, and we cannot rule out the influence of relaxed selection. Previous studies have demonstrated that positive selection will act on only a few sites for a short period of evolutionary time, and a signal of positive selection often is overwhelmed by continuous negative selection that sweeps across most sites in a gene sequence (Zhang *et al*. 2005).

In contrast to branch models where ω varies only among branches, branch-site models allow selection to vary both among amino acid sites and lineages. Thus, branch-site models are considered quite useful in distinguishing positive selection from relaxed or purifying selection (Zhang *et al*. 2005). Using the more stringent branch-site model, we detected positive selection in 14 branches and 12 genes with a total of 308 amino acid sites under positive selection. Among them, 80 amino acid sites in seven genes (*atp6*, *atp8*, *cox3*, *nad2*, *nad3*, *nad4*, and *nad5*) were identified on the subterranean branches, whereas 228 amino acid sites in 10 genes (*atp6*, *atp8*, *cox1*, *cox2*, *cytb*, *nad1*, *nad2*, *nad3*, *nad5*, and *nad6*) were identified on the surface branches. Nearly three times as many positively selected amino acid sites were detected on surface branches compared to subterranean branches. Most of the positively selected genes on surface branches were found in *C. forbesi* with 114 sites (Figure 7; Supplementary Table S7). In total, eight positive selected genes (*atp6*, *atp8*, *cox1*, *cox2*, *cytb*, *nad1*, *nad4*, and *nad5*) were identified by the branch-site model and by at least one other model on the surface branches, whereas only four positive selected genes (*atp6*, *atp8*, *cox3*, and *nad5*) were identified on the subterranean branches.

**Figure 7.**
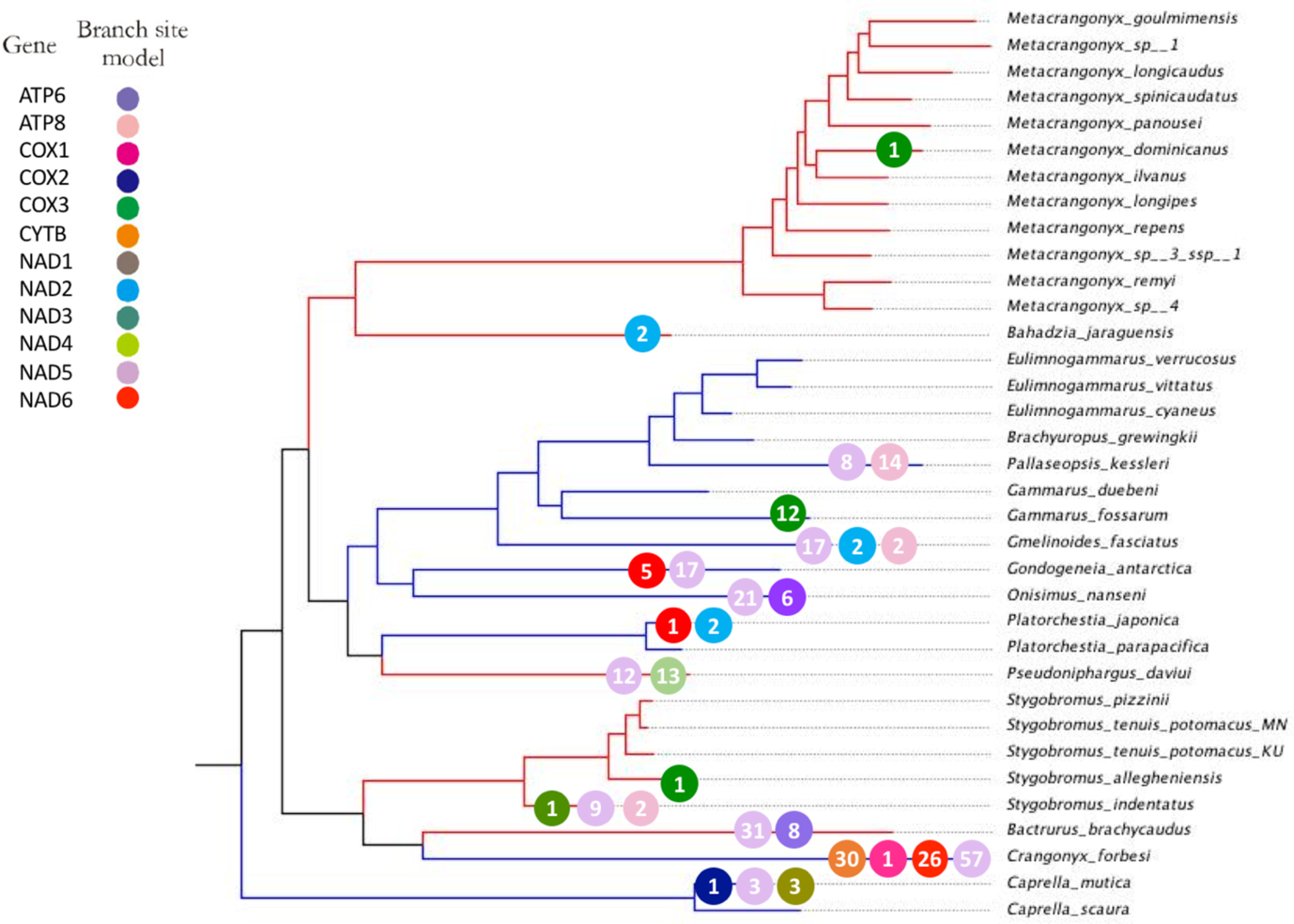
Evidence of positive selection on the mitochondrial PCGs with LRT P-value < 0.05 and positively selected site (BEB: P≥95%) in subterranean and surface-dwelling lineages of amphipods based on branch-site models. Different colored circles represent different mitochondrial genes. The number within each circle represents the number of positive selection sites detected for the gene. Surface amphipod branches are colored blue and subterranean amphipod branches are colored red.

The identification of many positively selected amino acid sites suggests that episodic positive selection has acted on mitochondrial PCGs of surface amphipods. In addition, we also identified a few positively selected sites on subterranean branches primarily in *B. brachycaudus* with 39 sites and *P. daviui* with 25 sites (Supplementary Table S7). *Bactrurus brachycaudus* is usually associated with springs and caves (Taylor and Niemiller 2016), whereas *P. daviui* associated with groundwater wells (Bauzà-Ribot *et al*. 2012).

### Direction and magnitude of selective pressures

Given the crucial role played by the mitochondrial genome in metabolic energy production (Hassanin *et al*. 2009), we hypothesized that the mitogenome of surface amphipods may show evidence of adaptation (directional selection) to life in surface habitats where energy demand is higher relative to subterranean habitats. We found support for directional selection in surface lineages based on three different selection analyses (RELAX, aBSREL, and BUSTED). In summary, all tests confirmed the existence of a moderate signal of positive or diversifying selection, as well as signal for significant relaxed purifying selection in the mitogenome of surface amphipods. This supports a previous study by Carlini and Fong (2017) who reported evidence for relaxation of functional evolutionary constraints (positive or diversifying selection) in the transcriptome of a subterranean amphipod *Gammarus minus*. The authors correlated the signal to the adaptation of this species to the subterranean environment.

We implemented aBSREL on the concatenated 13 PCG dataset comprising all 35 species as test branches and detected episodic diversifying selection in seven species: *P. daviui* (*p* = 0), *O. nanseni* (*p* = 0.0008), *G. fasciatus* (*p* = 0.0298), *G. fossarum* (*p* = 0.045), *B. jaraguensis* (*p* = 0.0016), *C. forbesi* (*p* = 0), and *B. brachycaudus* (*p* = 0.0001). We then used aBSREL to conduct independent tests for the crangonyctid species as the test branch and the remaining species as reference branches. We detected evidence of episodic diversifying selection in *C. forbesi* (*p* = 0) and *B. brachycaudus* (*p* = 0.0001) (Table 5). Using BUSTED, which provides a gene-wide test for positive selection, we detected evidence of episodic diversifying selection in three of the surface species: *C. forbesi* (*p* = 0.011), *G. fasciatus* (*p* = 0.033), *G. antarctica* (*p =* 0.009), whereas evidence of gene-wide episodic diversifying selection was found in just one of the subterranean species, *P. daviui* (*p* = 0.020) (Table 5). Using RELAX, which tests whether the strength of selection has been relaxed or intensified along a specified set of test branches, we detected selection evidence of relaxed selection in *C. forbesi* (*p* = 0) and other surface species, including *O. nanseni, G. fasciatus, G. fossarum, G. antarctica,* and *P. kessleri* with a *p* value of 0. Contrastingly, evidence of intensification of selection was detected in subterranean species including *S. tenuis* (*p* = 0), *S. allegheniensis* (*p* = 0.0025), *S. indentatus* (*p* = 0), and *S. pizzinii* (*p* = 0). Surprisingly, a few of the surface species including *C. mutica* (*p* = 0.015), *E. cyaneus* (*p* = 0), and *P. japonica* (*p* = 0) exhibited intensification of selection and subterranean species including *P. daviui* (*p* = 0) and *M. dominicanus* (*p* = 0.015) exhibited relaxation of selection (Table 5).

**Table 5.** Selection signals in the mitogenomes of amphipods inferred using aBSREL, BUSTED, and RELAX algorithms. The dataset comprising all 13 concatenated protein-coding genes with 3,607 amino acid sites in the alignment. K column: a statistically significant K > 1 indicates that selection strength has been intensified, and K < 1 indicates that selection strength has been relaxed. LR is likelihood ratio and D indicates the direction of selection pressure change: intensified (I) or relaxed (R), where * highlights a statistically significant (p < 0.05) result. Mitogenomes with significant LRT P-value < 0.05 are highlighted in red color

| Species | Genes-Sites | aBSREL | BUSTED | K | RELAX |  |  |
| --- | --- | --- | --- | --- | --- | --- | --- |
|  |  | p-value | p-value |  | p-value | LR | D |
| *Crangonyx forbesi* | 13PCG-3607 | 0.000 | 0.011 | 0.00 | 0.000 | 146.46 | R* |
| *Gmelinoides fasciatus* | 13PCG-3607 | 0.030 | 0.033 | 0.24 | 0.000 | 174.12 | R* |
| *Onisimus nanseni* | 13PCG-3607 | 0.001 | 0.500 | 0.25 | 0.000 | 80.59 | R* |
| *Gammarus fossarum* | 13PCG-3607 | 0.045 | 0.500 | 0.43 | 0.000 | 94.94 | R* |
| *Caprella mutica* | 13PCG-3607 | 0.500 | 0.500 | 1.12 | 0.015 | 5.97 | I* |
| *Eulimnogammarus cyaneus* | 13PCG-3607 | 0.500 | 0.448 | 2.20 | 0.000 | 77.75 | I* |
| *Platorchestia japonica* | 13PCG-3607 | 0.500 | 0.500 | 1.90 | 0.000 | 57.20 | I* |
| *Gondogeneia antarctica* | 13PCG-3607 | 0.500 | 0.009 | 0.28 | 0.000 | 49.83 | R* |
| *Pallaseopsis kessleri* | 13PCG-3607 | 0.500 | 0.500 | 1.00 | 0.000 | 15.68 | R* |
| *Brachyuropus grewingkii* | 13PCG-3607 | 0.500 | 0.500 | 1.03 | 0.624 | 0.24 | I |
| *Pseudoniphargus daviui* | 13PCG-3607 | 0.000 | 0.020 | 0.63 | 0.000 | 2721.06 | R* |
| *Metacrangonyx dominicanus* | 13PCG-3607 | 0.500 | 0.293 | 0.88 | 0.015 | 5.89 | R* |
| *Bahadzia jaraguensis* | 13PCG-3607 | 0.002 | 0.500 | 0.90 | 0.116 | 2.47 | R |
| *Bactrurus brachycaudus* | 13PCG-3607 | 0.000 | 0.500 | 0.90 | 0.119 | 2.43 | R |
| *Stygobromus tenuis potomacus_MN* | 13PCG-3607 | 0.500 | 0.500 | 48.44 | 0.000 | 66.82 | I* |
| *Stygobromus tenuis potomacus_KU* | 13PCG-3607 | 0.500 | 0.500 | 1.17 | 0.739 | 0.11 | I |
| *Stygobromus allegheniensis* | 13PCG-3607 | 0.500 | 0.500 | 49.37 | 0.025 | 5.02 | I* |
| *Stygobromus pizzinii* | 13PCG-3607 | 0.500 | 0.500 | 1.27 | 0.000 | 1852.16 | I* |
| *Stygobromus indentatus* | 13PCG-3607 | 0.500 | 0.500 | 1.28 | 0.000 | 16.42 | I* |

In addition to the concatenated 13 PCG dataset, we also conducted selection analyses for each PCG to determine which genes might be evolving under unique selection pressures. We found evidence of directional selection in *atp8* of *C. forbesi* (*p* = 0.026) and *nad3* of *S. pizzinii* (*p* = 0.041) using aBSREL and *cox3* of *B. brachycaudus* (*p* = 0.029) using BUSTED. *Atp8* of surface amphipod *C. forbesi* exhibiting strong evidence of directional selection is quite surprising as *atp8* is a small gene sometimes missing from metazoan mitogenomes and normally evolves under highly relaxed selection pressures (Gissi *et al*. 2008). Based on missing evidence for relaxation selection pressures in subterranean amphipods, we can confirm that *atp8* is indeed evolving under predominantly strong directional selection in surface amphipods which needs to be explored further. RELAX analyses uncovered five genes (*cox1, cox3, cytb, nad1,* and *nad3*) that exhibited relaxed selection and one gene (*atp6*) that exhibited intensification of selection in *C. forbesi*. Similarly, three genes (*cox3, nad5,* and *nad6*) in *B. brachycaudus* showed evidence of relaxed selection. Several genes in other subterranean species, including S. *tenuis, S. allegheniensis,* and *S. pizzinii*, exhibited varying levels of intensification of selection, whereas none exhibited relaxed selection (Table 6). Some of these outliers were expected, as *nad5* and *nad6* are known to evolve faster among the mitochondrial genes (Zhang *et al*. 2018). Also, evidence for relaxation of functional evolutionary constraints (positive or diversifying selection) has been reported in the *nad* family of subterranean *Gammarus* species adapted to the subterranean environment (Carlini and Fong 2017). Although this may explain outliers in the subterranean *B. brachycaudus* mitogenome, it remains unclear why *cox3* exhibited signatures of relaxed selection. This gene is generally one of the most conserved mitochondrial loci in animals (Oliveira *et al*. 2008; Xiao *et al*. 2011; Pons *et al*. 2014), and high levels of purifying selection has been observed in the *cox* family in other amphipod species (Sun *et al*. 2020). In *C. forbesi*, *atp6* showed signatures of positive selection, which contrasted most other PCGs in its mitogenome that exhibited relaxed selection. Overall, in accordance with the results obtained using the concatenated dataset, individual mitochondrial genes of subterranean amphipods mostly exhibited varying levels of purifying selection, whereas surface amphipods predominantly exhibited relaxed selection.

**Table 6.** Selection signals in the mitochondrial PCGs of crangonyctid amphipods sequenced in this study inferred using aBSREL, BUSTED, and RELAX algorithms. K column: a statistically significant K > 1 indicates that selection strength has been intensified, and K < 1 indicates that selection strength has been relaxed. LR is likelihood ratio and D indicates the direction of selection pressure change: intensified (I) or relaxed (R), where * highlights a statistically significant (p < 0.05) result. PCGs with significant LRT P-value < 0.05 are highlighted in red color

| Gene | Site<br>s | *Crangonyx forbesi* |  |  |  | *Bactrurus brachycaudus* |  |  |  | *Stygobromus tenuis*<br>*potomacus* |  |  |  | *Stygobromus allegheniensis* |  |  |  | *Stygobromus pizzinii* |  |  |  |
| --- | --- | --- | --- | --- | --- | --- | --- | --- | --- | --- | --- | --- | --- | --- | --- | --- | --- | --- | --- | --- | --- |
|  |  | aBSR<br>EL | BUS<br>TED | RELAX | D | aBSR<br>EL | BUS<br>TED | RELAX | D | aBSR<br>EL | BUS<br>TED | RELAX | D | aBSR<br>EL | BUST<br>ED | RELAX | D | aBSR<br>EL | BUST<br>ED | RELAX | D |
|  |  | p-value | p-value | p-value |  | p-value | p-value | p-value |  | p-value | p-value | p-value |  | p-value | p-value | p-value |  | p-value | p-value | p-value |  |
| *atp6* | 218 | 0.500 | 0.272 | 0.001 | I* | 0.500 | 0.357 | 0.648 | R | 1.000 | 0.500 | 1.000 | R | 0.069 | 0.500 | 0.000 | I* | 1.000 | 0.500 | 0.127 | I |
| *atp8* | 50 | 0.026 | 0.229 | 0.625 | I | 1.000 | 0.500 | 0.214 | I | 0.500 | 0.500 | 0.510 | R | 0.435 | 0.478 | 0.142 | R | 0.444 | 0.500 | 0.336 | R |
| *cox1* | 511 | 0.056 | 0.146 | 0.000 | R* | 0.105 | 0.500 | 1.000 | R | 1.000 | 0.500 | 0.000 | I* | 1.000 | 0.500 | 0.150 | I | 1.000 | 0.500 | 0.044 | I* |
| *cox2* | 222 | 0.500 | 0.476 | 0.634 | I | 0.188 | 0.280 | 0.262 | R | 1.000 | 0.500 | 0.017 | I* | 0.500 | 0.500 | 0.359 | I | 1.000 | 0.500 | 0.096 | I |
| *cox3* | 262 | 1.000 | 0.500 | 0.032 | R* | 0.199 | 0.029 | 0.023 | R* | 1.000 | 0.500 | 1.000 | R | 1.000 | 0.500 | 0.000 | I* | 0.367 | 0.500 | 0.018 | I* |
| *cytb* | 377 | 0.143 | 0.500 | 0.000 | R* | 0.500 | 0.500 | 0.112 | I | 1.000 | 0.500 | 0.080 | I | 0.230 | 0.500 | 0.672 | I | 1.000 | 0.500 | 0.536 | I |
| *nad1* | 305 | 0.093 | 0.500 | 0.000 | R* | 1.000 | 0.500 | 0.067 | I | 1.000 | 0.500 | 0.001 | I* | 0.151 | 0.500 | 0.034 | I* | 1.000 | 0.500 | 0.125 | I |
| *nad2* | 321 | 1.000 | 0.500 | 0.386 | R | 1.000 | 0.500 | 0.363 | R | 1.000 | 0.500 | 0.220 | I | 1.000 | 0.500 | 0.002 | I* | 1.000 | 0.500 | 0.141 | I |
| *nad3* | 116 | 0.414 | 0.308 | 0.000 | R* | 1.000 | 0.500 | 0.752 | R | 1.000 | 0.500 | 0.000 | I* | 0.500 | 0.500 | 0.118 | I | 0.041 | 0.500 | 0.149 | R |
| *nad4* | 435 | 1.000 | 0.146 | 0.068 | R | 1.000 | 0.500 | 0.482 | I | 1.000 | 0.500 | 0.076 | I | 1.000 | 0.500 | 0.105 | I | 1.000 | 0.500 | 0.008 | I* |
| *nad4l* | 94 | 1.000 | 0.500 | 0.978 | I | 1.000 | 0.500 | 0.854 | I | 1.000 | 0.500 | 0.018 | I* | 0.500 | 0.500 | 0.980 | I | 1.000 | 0.500 | 1.000 | I |
| *nad5* | 552 | 1.000 | 0.500 | 0.774 | I | 0.088 | 0.479 | 0.024 | R* | 1.000 | 0.500 | 0.005 | I* | 0.447 | 0.500 | 0.012 | I* | 1.000 | 0.500 | 0.003 | I* |
| *nad6* | 144 | 0.500 | 0.500 | 0.250 | R | 1.000 | 0.500 | 0.036 | R* | 1.000 | 0.500 | 0.643 | R | 1.000 | 0.500 | 0.517 | I | 0.500 | 0.500 | 0.680 | R |

To provide further evidence of positive selection, we implemented the RELAX, aBSREL, and BUSTED algorithms on the branch, branch-site, and site models. Eight genes (*atp8*, *cox1*, *cox2*, *cytb*, *nad1*, *nad4*, *nad5*, and *nad6*) all involved in the OXPHOS pathway were under positive selection in surface branches by at least two methods. The genes *nad1*, *nad4*, *nad5*, and *nad6* encode the subunits of NADH dehydrogenase also called Complex I that initiates the oxidative phosphorylation process. Complex I is the largest and complicated proton pump of the respiratory chain involved in electron transfer from NADH to ubiquinone to supply the proton motive force used for ATP synthesis (Wirth *et al*. 2016) and plays a key role in cellular energy metabolism by pumping gradient of protons across the mitochondrial membrane producing more than one-third of mitochondrial energy (Dröse *et al*. 2011). This would clarify the reason behind detecting more evidence of positive selection in complex I than in other complexes. Genes *cox1* & *cox2* which acts as a regulator encodes the catalytic core of Cytochrome c oxidase also called as Complex IV. Complex IV is directly involved in electron transfer and proton translocation (Zhang *et al*. 2013). Gene *atp8* encodes a part of ATP synthase also called as Complex V and plays a major role in the final assembly of ATPase (Zhang *et al*. 2013). In summary, our selection analyses revealed signals of positive selection in several mitochondrial genes of surface amphipods, which may be associated with increased energy demands in surface environments. In contrast, subterranean amphipods showed signatures of purifying selection, which may be related to maintaining efficient energy metabolism in subterranean habitats.

## Conclusion

In this study, we compared mitogenome features including AT/GC-skew, codon usage, gene order, phylogenetic relationships, and selection pressures acting upon amphipods inhabiting surface and subterranean habitats. We describe a novel mitochondrial gene order for *C. forbesi*. We identified a signal of directional selection in the protein-encoding genes of the OXPHOS pathway in the mitogenomes of surface amphipods and a signal of purifying selection in subterranean species, which is consistent with the hypothesis that the mitogenome of surface-adapted amphipods has evolved in response to a more energy demanding environment compared to subterranean species. Our comparative analyses of gene order, locations of non-coding regions, and base-substitution rates points to habitat as an important factor influencing the evolution of amphipod mitogenomes. However, the generation and study of mitogenomes from additional amphipod taxa, including other crangonyctid species, are needed to better elucidate phylogenetic relationships and the evolution of mitogenomes of amphipods. In addition, more evidence is needed to further validate our inferences, such as studying the effects of amino acid changes on three-dimensional protein structure and function. Nevertheless, our study provides a necessary foundation for the study of mitogenome evolution in amphipods and other crustaceans.

