## Supplementary Materials for "Comparative mitogenomic analysis of subterranean and surface amphipods (Crustacea, Amphipoda) with special reference to the family Crangonyctidae"

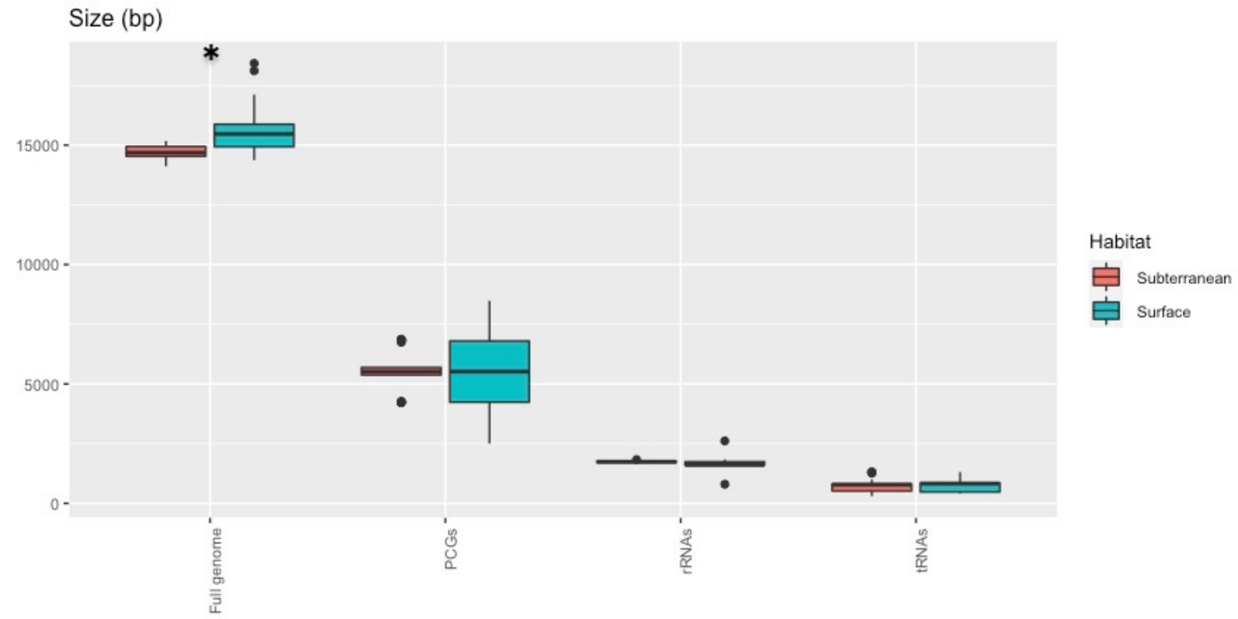


**Supplementary Figure S1.** Box plot showing size (i.e., length) in bp of mitogenomes, protein coding genes (PCG), ribosomal (rRNAs) loci, and transfer ribosomal (tRNAs) loci between subterranean and surface amphipods. Significant P-value < 0.05 is indicated using *.

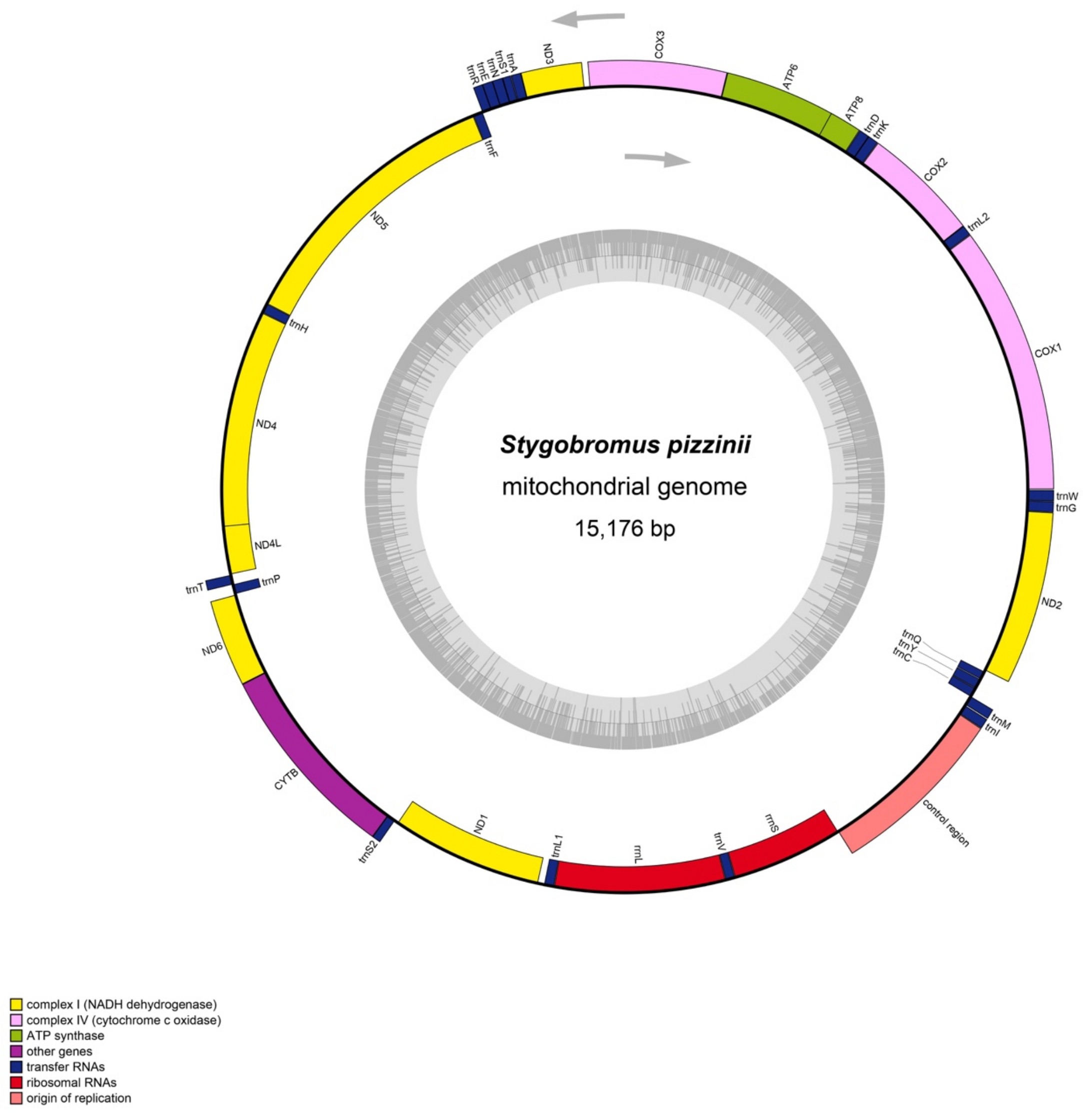

Stygobromus pizzinii
mitochondrial genome
15,176 bp
COX3
ATP6
ATP8
trnD
trnK
COX2
trnL2
COX1
trnW
trnG
ND2
trnQ
trnY
trnC
trnM
trnI
control region
rrnS
trnV
rrnL
trnL1
ND1
trnS2
CYTB
ND6
trnT
trnP
ND4L
ND4
trnH
ND5
trnF
trnR
trnE
trnN
trnS1
trnA
ND3
complex I (NADH dehydrogenase)
complex IV (cytochrome c oxidase)
ATP synthase
other genes
transfer RNAs
ribosomal RNAs
origin of replication

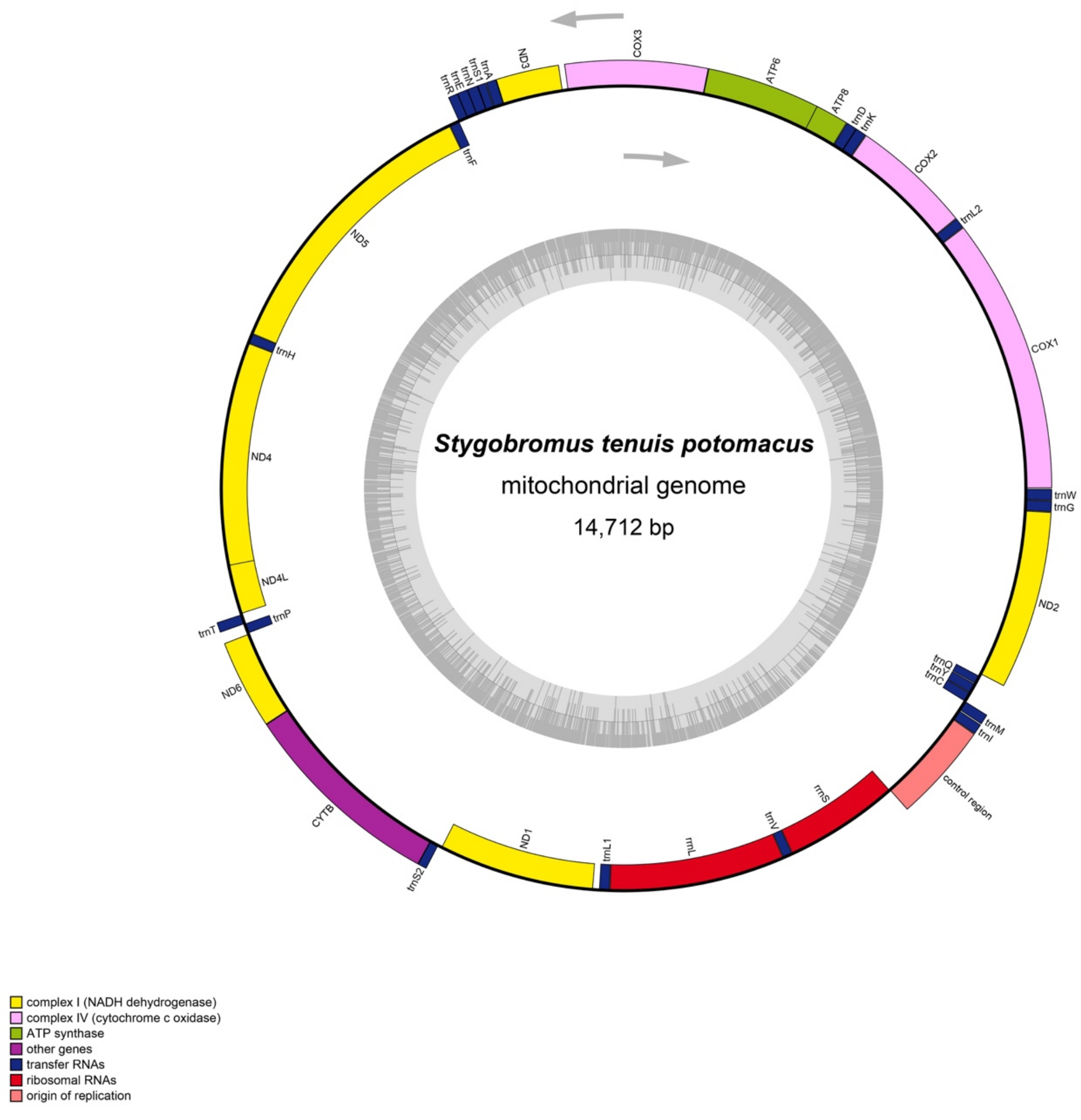

Stygobromus tenuis potomacus
mitochondrial genome
14,712 bp
COX3
ATP6
ATP8
trnD
trnK
COX2
trnL2
COX1
trnW
trnG
ND2
trnQ
trnY
trnC
trnM
trnI
control region
rrnS
trnV
rrnL
trnL1
ND1
trnS2
CYTB
ND6
trnT
trnP
ND4L
ND4
trnH
ND5
trnF
trnR
trnE
trnN
trnS1
trnA
ND3
complex I (NADH dehydrogenase)
complex IV (cytochrome c oxidase)
ATP synthase
other genes
transfer RNAs
ribosomal RNAs
origin of replication

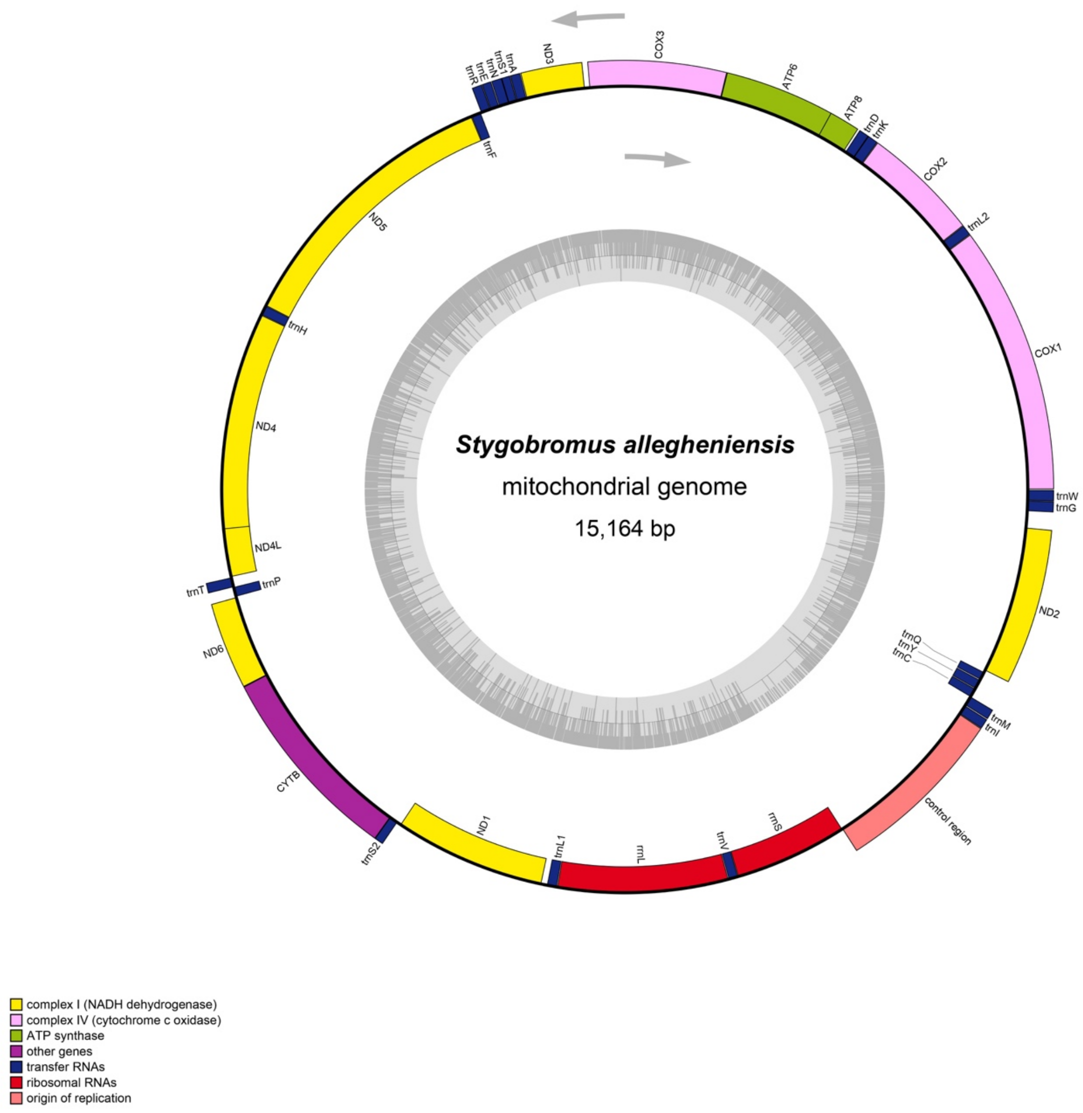

Stygobromus allegheniensis
mitochondrial genome
15,164 bp
COX3
ATP6
ATP8
trnD
trnK
COX2
trnL2
COX1
trnW
trnG
ND2
trnQ
trnY
trnC
trnM
trnI
control region
rrnS
trnV
rrnL
trnL1
ND1
trnS2
CYTB
ND6
trnT
trnP
ND4L
ND4
trnH
ND5
trnF
trnR
trnE
trnN
trnS1
trnA
ND3



complex I (NADH dehydrogenase)
complex IV (cytochrome c oxidase)
ATP synthase
other genes
transfer RNAs
ribosomal RNAs
origin of replication

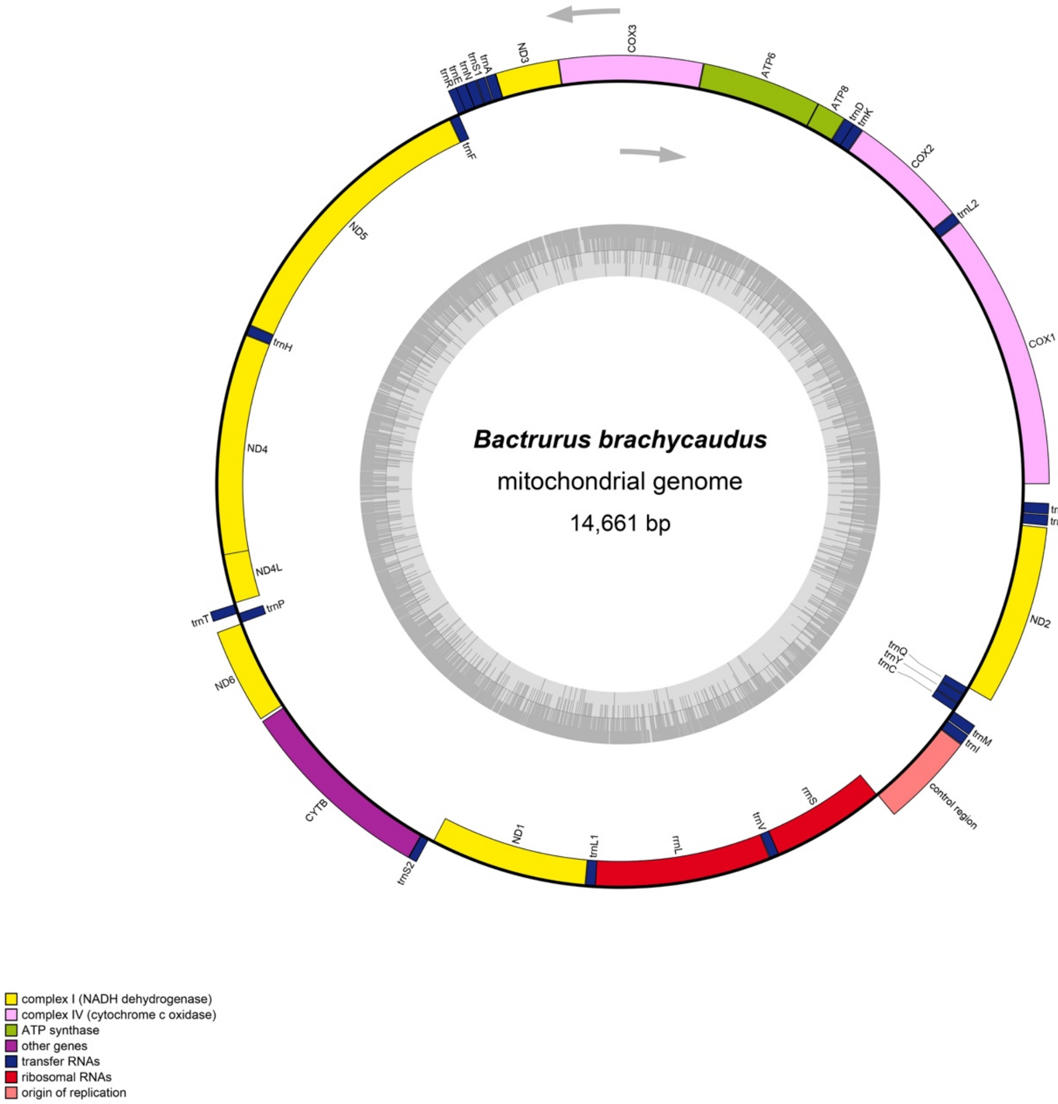
Bactrurus brachycaudus
mitochondrial genome
14,661 bp
COX3
ATP6
ATP8
trnD
trnK
COX2
trnL2
COX1
trnG
trnW
ND2
trnQ
trnY
trnC
trnM
trnI
control region
rrnS
trnV
rrnL
trnL1
ND1
trnS2
CYTB
ND6
trnT
trnP
ND4L
ND4
trnH
ND5
trnF
trnR
trnE
trnN
trnS1
trnA
ND3
complex I (NADH dehydrogenase)
complex IV (cytochrome c oxidase)
ATP synthase
other genes
transfer RNAs
ribosomal RNAs
origin of replication

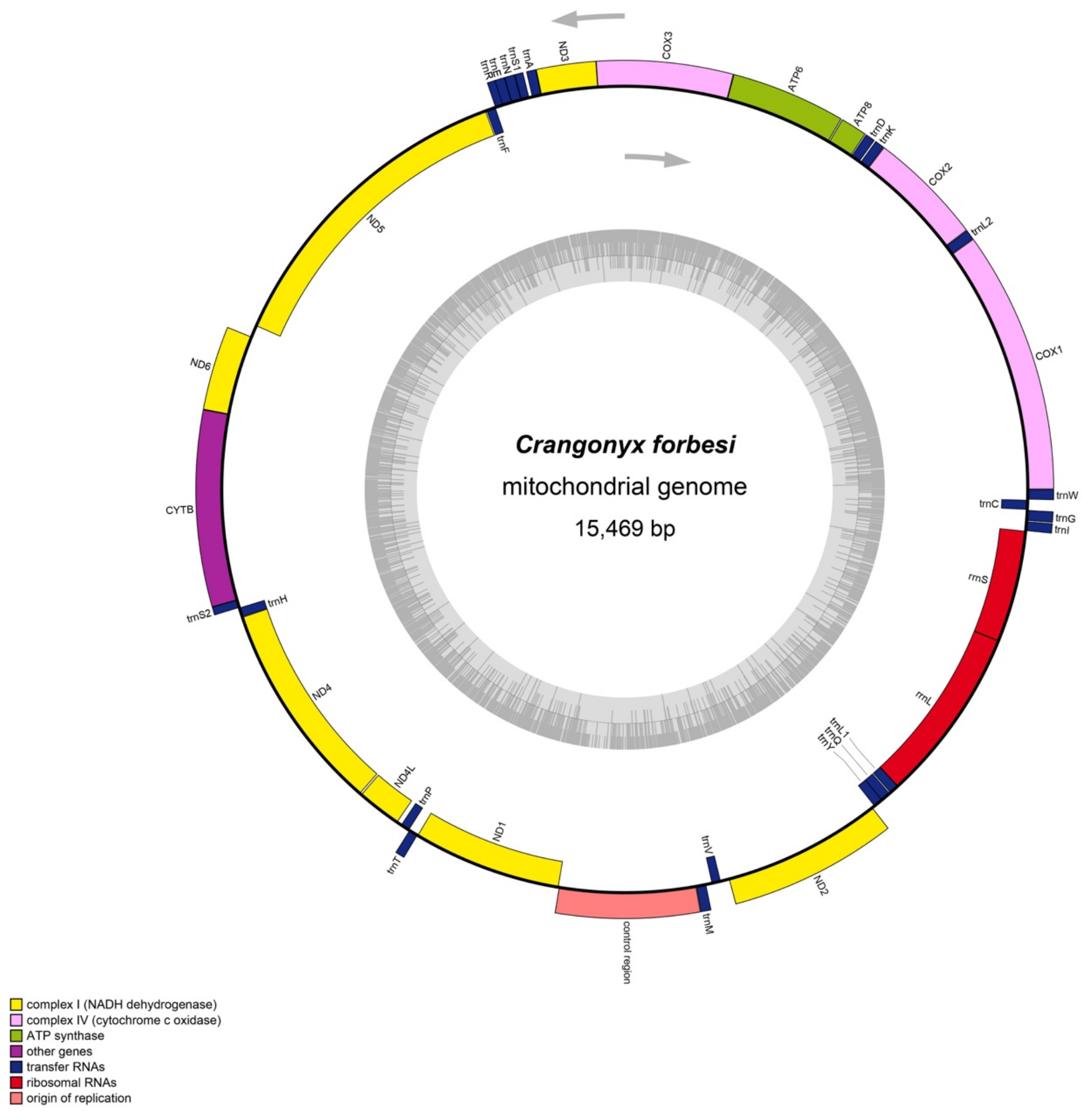


**Supplementary Figure S2**. Map of the mitochondrial genome of *S. pizzinii, S. tenuis potomacus, Stygobromus allegheniensis, Bactrurus brachycaudus,* and *Crangonyx forbesi*

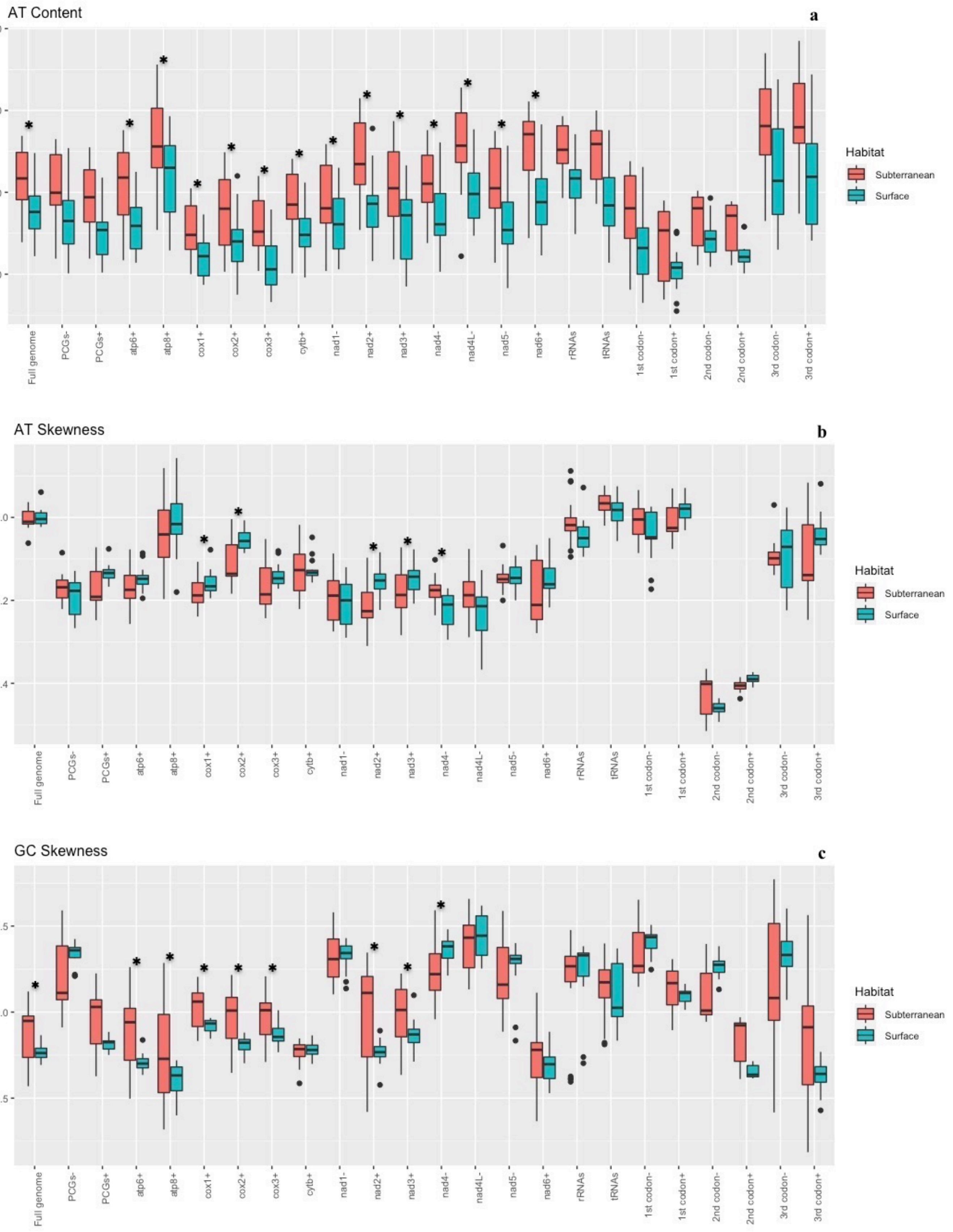
AT Content
a
90
80
70
60
Full genome
PCGs-
PCGs+
atp6+
atp8+
cox1+
cox2+
cox3+
cytb+
nad1-
nad2+
nad3+
nad4-
nad4L-
nad5-
nad6+
rRNAs
tRNAs
1st codon-
1st codon+
2nd codon-
2nd codon+
3rd codon-
3rd codon+
Habitat
Subterranean
Surface
AT Skewness
b
0.0
-0.2
-0.4
GC Skewness
c
0.5
0.0
-0.5

**Supplementary Figure S3.** Box plot showing AT percentage **(a)**, AT-skew **(b)**, and GC-skew **(c)** between subterranean and surface amphipods across mitogenomes, protein coding genes (PCG), ribosomal (rRNA) loci, and transfer ribosomal (tRNA) loci. The same features are shown for each protein-coding gene and pooled by codon position and coding strand. Genes coded on the (-) strand are represented by a "-" sign and genes coded on the (+) strand are represented by "+" sign at the end of the gene label. Mitogenome and PCG with significant P-value < 0.05 are indicated using *.

### CREx: comparison

back to distance matrix

**Pan-crustacean → Stygobromus_pizzinii_MN175620**

- family diagram for Pan-crustacean (e)

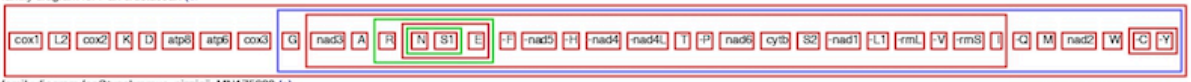


- family diagram for Stygobromus_pizzinii_MN175620 (e)

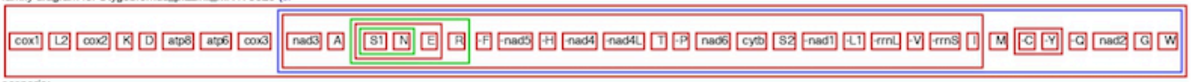


- scenario:
  - transposition

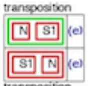


  - transposition

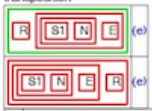


  - 

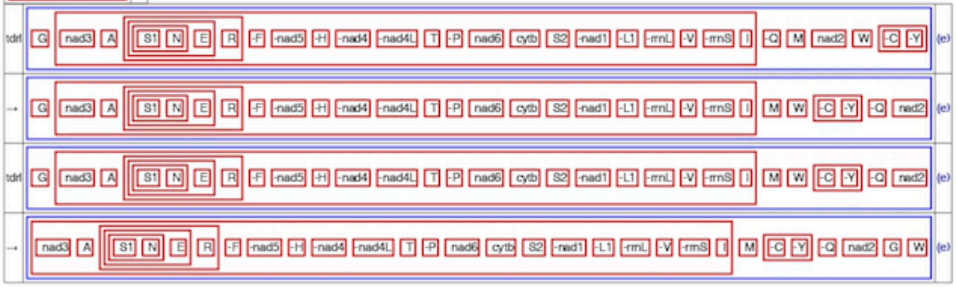


**Stygobromus_pizzinii_MN175620 → Pan-crustacean**

- family diagram for Stygobromus_pizzinii_MN175620 (e)

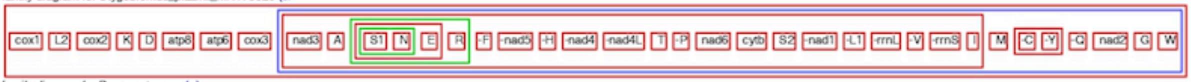


- family diagram for Pan-crustacean (e)

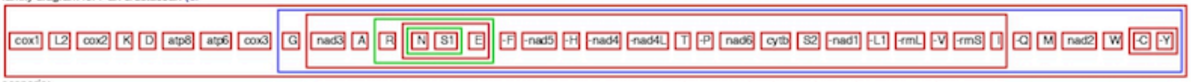


- scenario:
  - transposition

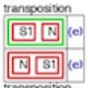


  - transposition

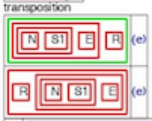


  - 

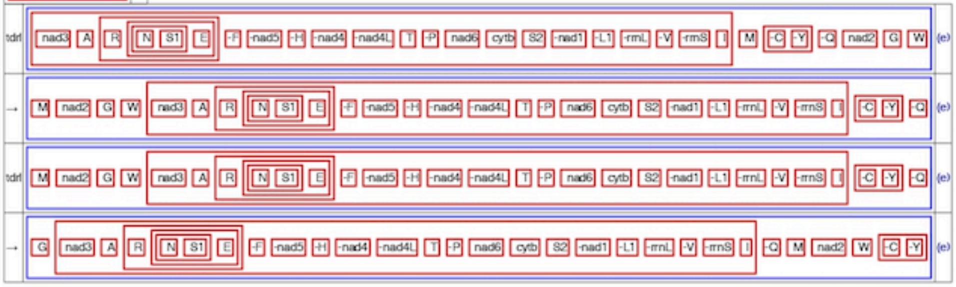

### CREx: comparison

back to distance matrix

#### Pan-crustacean → Bactrurus_brachycaudus_MN176619

- family diagram for Pan-crustacean (e)

  cox1 L2 cox2 K D atp8 atp6 cox3 G nad3 A R N S1 E -F -nad5 -H -nad4 -nad4L T -P nad6 cytb S2 -nad1 -L1 -rrnL -V -rrnS I -Q M nad2 W -C -Y

- family diagram for Bactrurus_brachycaudus_MN175619 (e)

  cox1 L2 cox2 K D atp8 atp6 cox3 nad3 A S1 N E R -F -nad5 -H -nad4 -nad4L T -P nad6 cytb S2 -nad1 -L1 -rrnL -V -rrnS I M -C -Y -Q nad2 W G

- scenario:
  - transposition

    N S1 (e)

    S1 N (e)
  - transposition

    R S1 N E (e)

    S1 N E R (e)
  - transposition

    G nad3 A S1 N E R -F -nad5 -H -nad4 -nad4L T -P nad6 cytb S2 -nad1 -L1 -rrnL -V -rrnS I -Q M nad2 W -C -Y (e)

    nad3 A S1 N E R -F -nad5 -H -nad4 -nad4L T -P nad6 cytb S2 -nad1 -L1 -rrnL -V -rrnS I -Q M nad2 W -C -Y G (e)
  - tdrl

    -Q M nad2 W -C -Y (e)

    M -C -Y -Q nad2 W (e)

#### Bactrurus_brachycaudus_MN176619 → Pan-crustacean

- family diagram for Bactrurus_brachycaudus_MN175619 (e)

  cox1 L2 cox2 K D atp8 atp6 cox3 nad3 A S1 N E R -F -nad5 -H -nad4 -nad4L T -P nad6 cytb S2 -nad1 -L1 -rrnL -V -rrnS I M -C -Y -Q nad2 W G

- family diagram for Pan-crustacean (e)

  cox1 L2 cox2 K D atp8 atp6 cox3 G nad3 A R N S1 E -F -nad5 -H -nad4 -nad4L T -P nad6 cytb S2 -nad1 -L1 -rrnL -V -rrnS I -Q M nad2 W -C -Y

- scenario:
  - transposition

    S1 N (e)

    N S1 (e)
  - transposition

    N S1 E R (e)

    R N S1 E (e)
  - transposition

    nad3 A R N S1 E -F -nad5 -H -nad4 -nad4L T -P nad6 cytb S2 -nad1 -L1 -rrnL -V -rrnS I M -C -Y -Q nad2 W G (e)

    G nad3 A R N S1 E -F -nad5 -H -nad4 -nad4L T -P nad6 cytb S2 -nad1 -L1 -rrnL -V -rrnS I M -C -Y -Q nad2 W (e)
  -

| | | |
|---|---|---|
| transposition | M -C -Y -Q nad2 W | (e) |
| → | -C -Y -Q M nad2 W | (e) |
| transposition | -C -Y -Q M nad2 W | (e) |
| → | -Q M nad2 W -C -Y | (e) |

### CREx: comparison

back to distance matrix

**Pan-crustacean → Crangonyx_forbesi_MN175623**

- family diagram for Pan-crustacean (e)

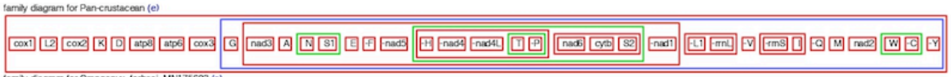


- family diagram for Crangonyx_forbesi_MN175623 (e)

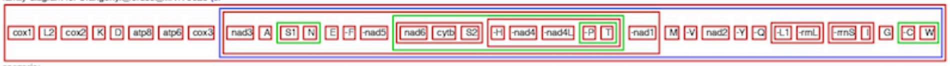


- scenario:
  - transposition

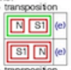


  - transposition

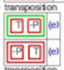


  - transposition

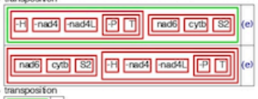


  - transposition

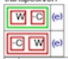


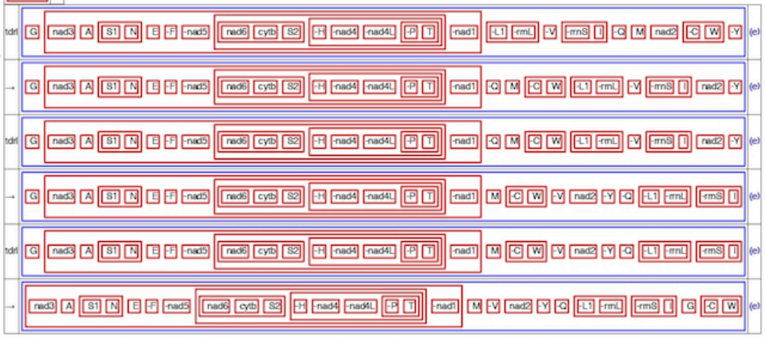

**Crangonyx_forbesi_MN176623 → Pan-crustacean**

- family diagram for Crangonyx_forbesi_MN175623 (e)

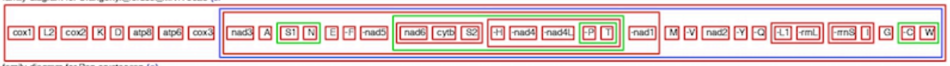


- family diagram for Pan-crustacean (e)

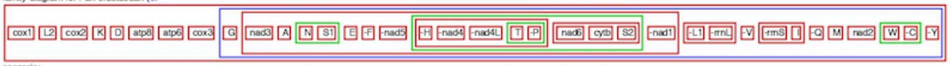


- scenario:
  - transposition

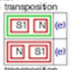


  - transposition

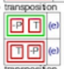


  - transposition

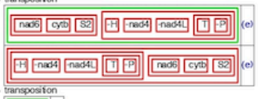


  - transposition

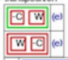


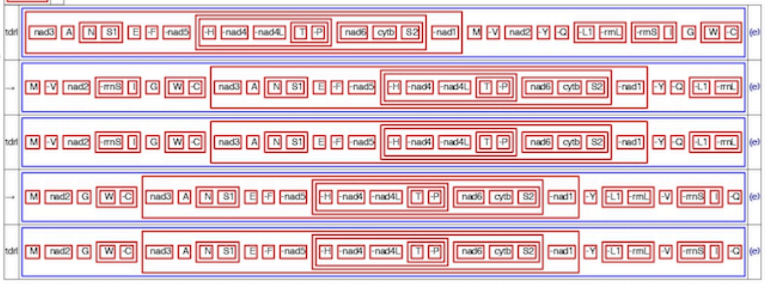

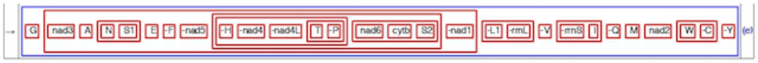

**Supplementary Figure S4**. CREx analysis showing the possible scenarios for the evolution of gene rearrangements in the crangonyctid amphipod genus *Stygobromus*, *Bactrurus*, and *Crangonyx* from the ancestral pan-crustacean pattern

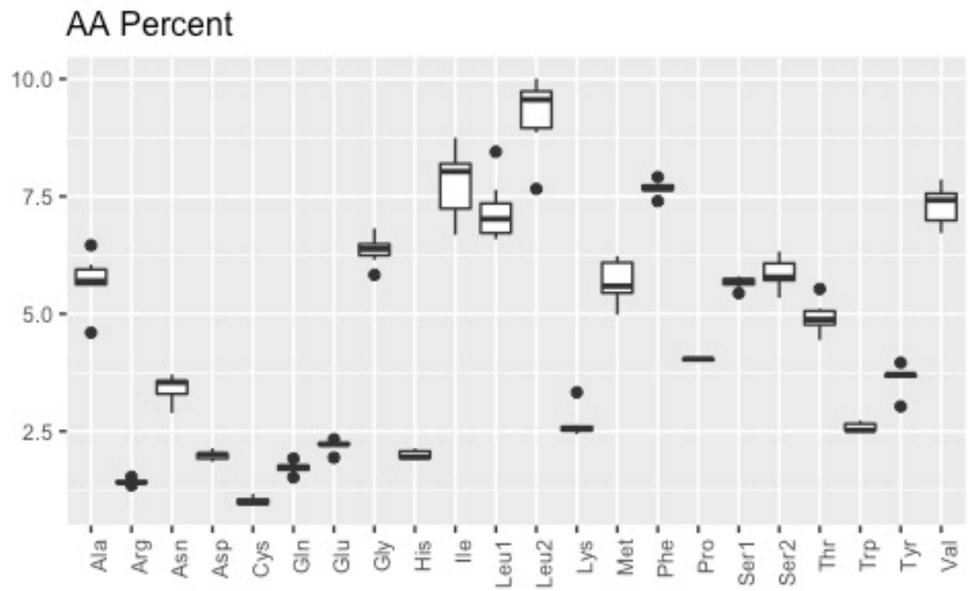


**Supplementary Figure S5.** Box plot showing amino acid composition for mitochondrial protein-coding gene across crangonyctid mitogenomes.

### *Stygobromus pizzinii*

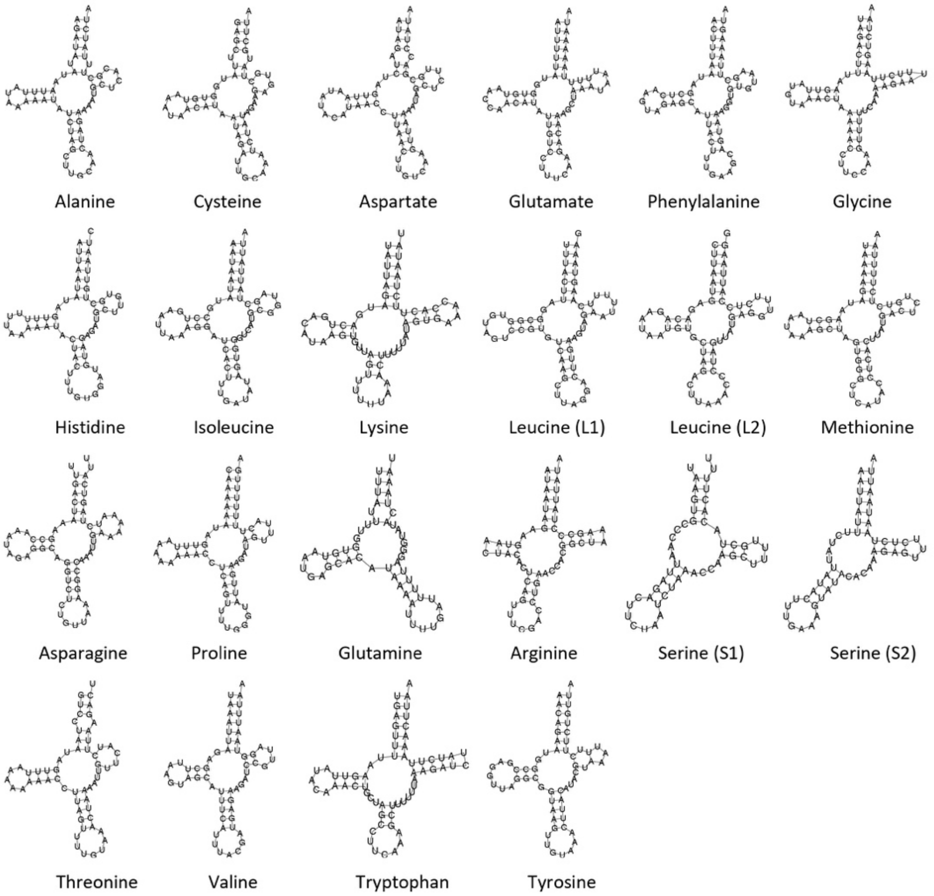


### *Stygobromus tenuis potomacus*

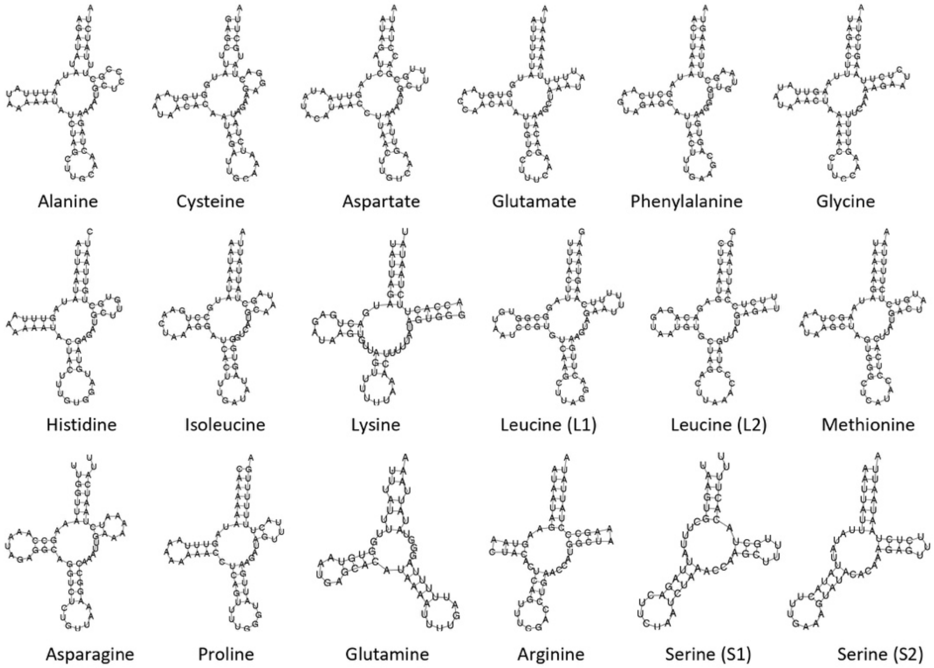

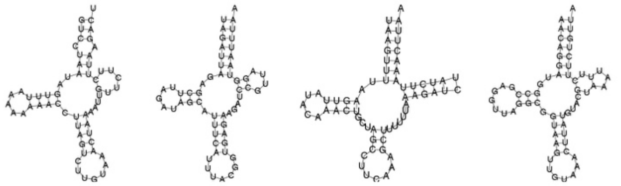

Threonine Valine Tryptophan Tyrosine

### ***Bactrurus brachycaudus***

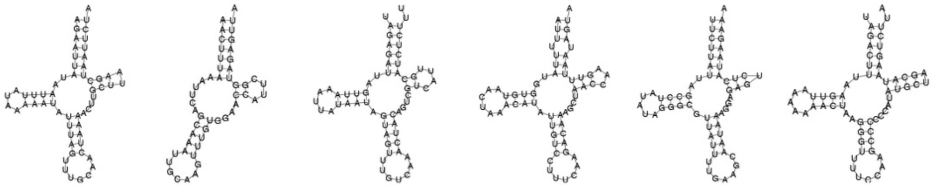

Alanine Cysteine Aspartate Glutamate Phenylalanine Glycine

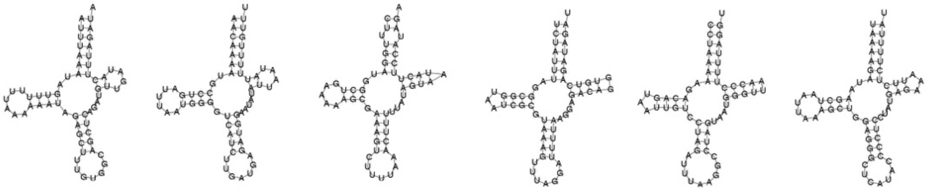

Histidine Isoleucine Lysine Leucine (L1) Leucine (L2) Methionine

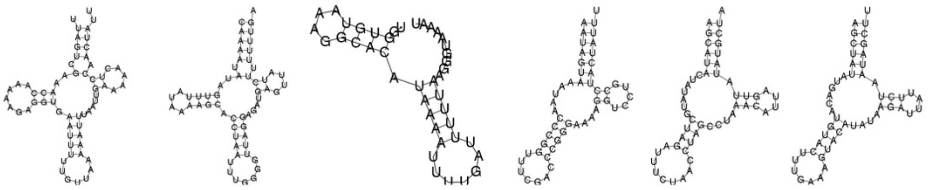

Asparagine Proline Glutamine Arginine Serine (S1) Serine (S2)

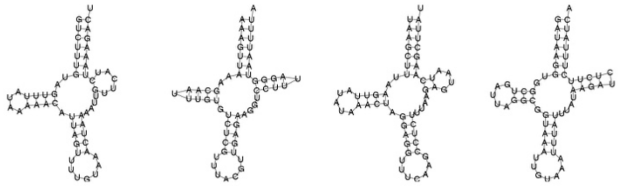

Threonine Valine Tryptophan Tyrosine

### ***Stygobromus allegheniensis***

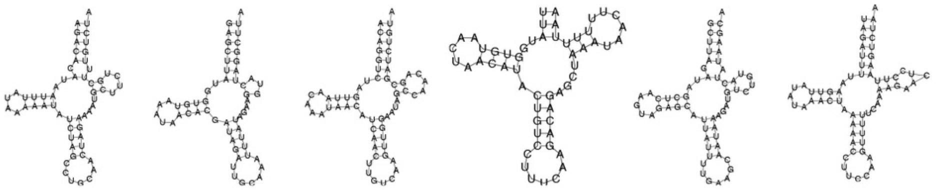

Alanine Cysteine Aspartate Glutamate Phenylalanine Glycine

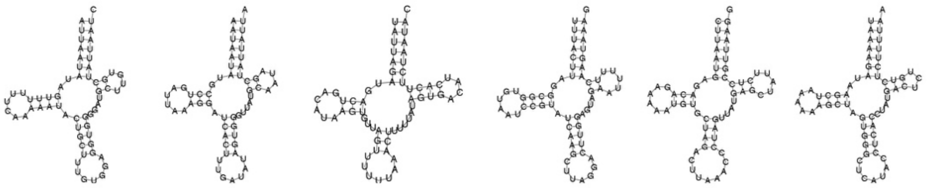

Histidine Isoleucine Lysine Leucine (L1) Leucine (L2) Methionine

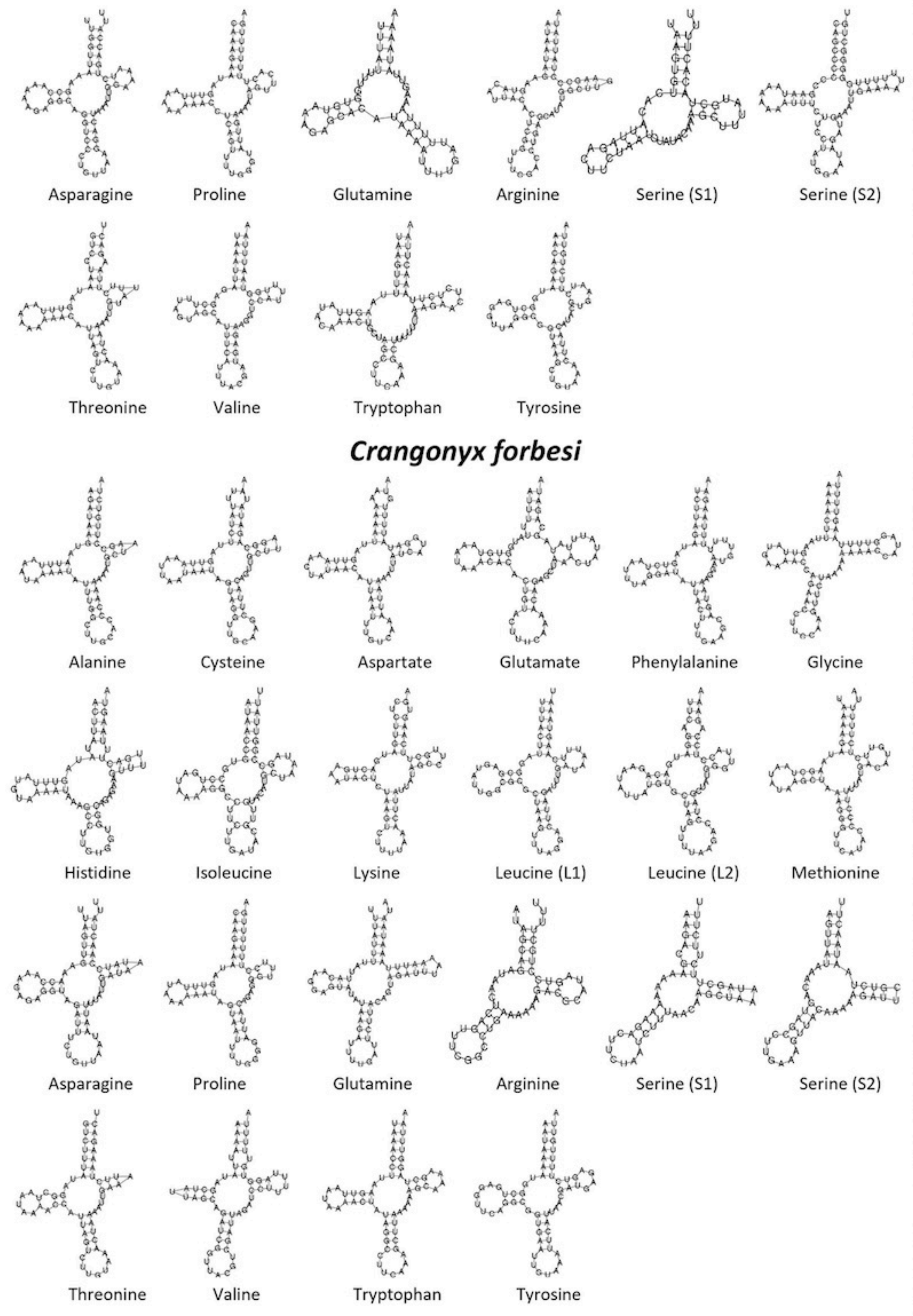


**Supplementary Figure S6.** The predicted mitochondrial tRNAs secondary structures of crangonyctid amphipods under study

**Supplementary Table S1.** Organization of the mitochondrial genomes of crangonyctid amphipods under study.

***Bactrurus brachycaudus***

| Gene | Position | | Size | Intergenic nucleotides | Codon | | |
|---|---|---|---|---|---|---|---|
| | From | To | | | Start | Stop | Strand |
| cox1 | 1 | 1534 | 1534 | | ATC | T | H |
| *trnL2* | 1535 | 1594 | 60 | | | | H |
| *cox2* | 1595 | 2264 | 670 | | ATG | T | H |
| trnK | 2265 | 2323 | 59 | | | | H |
| *trnD* | 2320 | 2379 | 60 | -4 | | | H |
| *atp8* | 2380 | 2544 | 165 | | ATC | TAA | H |
| *atp6* | 2544 | 3206 | 663 | -1 | ATG | TAA | H |
| *cox3* | 3207 | 4002 | 796 | | ATG | T | H |
| *nad3* | 4003 | 4353 | 351 | | ATC | TAG | H |
| *trnA* | 4352 | 4410 | 59 | -2 | | | H |
| *trnS1* | 4412 | 4463 | 52 | 1 | | | H |
| *trnN* | 4464 | 4525 | 62 | | | | H |
| trnE | 4523 | 4584 | 62 | -3 | | | H |
| *trnR* | 4578 | 4627 | 50 | -7 | | | H |
| *trnF* | 4625 | 4683 | 59 | -3 | | | L |
| *nad5* | 4684 | 6382 | 1699 | | ATT | T | L |
| trnH | 6383 | 6443 | 61 | | | | L |
| *nad4* | 6444 | 7755 | 1312 | | ATC | T | L |
| *nad4L* | 7740 | 8030 | 291 | -16 | ATG | TAG | L |
| *trnT* | 8035 | 8092 | 58 | 4 | | | H |
| trnP | 8092 | 8150 | 59 | -1 | | | L |
| *nad6* | 8153 | 8683 | 531 | 2 | ATG | TAG | H |
| *cytb* | 8697 | 9794 | 1098 | 13 | ATC | TAG | H |
| S_copy2 | 9793 | 9843 | 51 | -2 | | | H |
| *nad1* | 9871 | 10794 | 924 | 27 | TTG | TAA | L |
| *L_copy2* | 10795 | 10854 | 60 | | | | L |
| *rrnL* | 10851 | 11884 | 1034 | -4 | | | L |
| *trnV* | 11885 | 11941 | 57 | | | | L |
| rrnS | 11939 | 12609 | 671 | -3 | | | L |
| *control region* | 12610 | 13140 | 531 | | | | H |
| *trnI* | 13141 | 13202 | 62 | | | | H |
| trnM | 13206 | 13267 | 62 | 3 | | | H |

| | | | | | | | |
|---|---|---|---|---|---|---|---|
| *trnC* | 13267 | 13316 | 50 | -1 | | | L |
| trnY | 13317 | 13376 | 60 | | | | L |
| *trnQ* | 13377 | 13426 | 50 | | | | L |
| *nad2* | 13427 | 14425 | 999 | | ATT | TAA | H |
| trnW | 14435 | 14494 | 60 | 9 | | | H |
| *trnG* | 14497 | 14558 | 62 | 2 | | | H |
| *Overlap:* | 12 | gap: | 8 | | | | |

***Stygobromus pizzinii***

| Gene | Position | | Size | Intergenic nucleotides | Codon | | |
|---|---|---|---|---|---|---|---|
| | **From** | **To** | | | **Start** | **Stop** | **Strand** |
| *cox1* | 1 | 1534 | 1534 | | ATA | T | H |
| *trnL2* | 1535 | 1596 | 62 | | | | H |
| *cox2* | 1596 | 2270 | 675 | -1 | GTG | TAA | H |
| *trnK* | 2271 | 2333 | 63 | | | | H |
| *trnD* | 2332 | 2393 | 62 | -2 | | | H |
| *atp8* | 2394 | 2582 | 189 | | ATC | TAG | H |
| *atp6* | 2542 | 3210 | 669 | -41 | ATG | TAA | H |
| cox3 | 3210 | 4004 | 795 | -1 | ATG | TAA | H |
| *nad3* | 4036 | 4386 | 351 | 31 | ATT | TAG | H |
| *trnA* | 4385 | 4443 | 59 | -2 | | | H |
| *trnS1* | 4444 | 4493 | 50 | | | | H |
| trnN | 4492 | 4553 | 62 | -2 | | | H |
| *trnE* | 4551 | 4612 | 62 | -3 | | | H |
| *trnR* | 4610 | 4666 | 57 | -3 | | | H |
| *trnF* | 4665 | 4724 | 60 | -2 | | | L |
| *nad5* | 4725 | 6432 | 1708 | | ATT | T | L |
| *trnH* | 6433 | 6492 | 60 | | | | L |
| *nad4* | 6493 | 7813 | 1321 | | ATG | T | L |
| *nad4L* | 7807 | 8100 | 294 | -7 | ATG | TAA | L |
| *trnT* | 8104 | 8162 | 59 | 3 | | | H |
| trnP | 8162 | 8221 | 60 | -1 | | | L |
| *nad6* | 8226 | 8726 | 501 | 4 | ATG | TAA | H |
| *cytb* | 8726 | 9865 | 1140 | -1 | ATG | TAA | H |
| S_copy2 | 9864 | 9916 | 53 | -2 | | | H |
| *nad1* | 9937 | 10857 | 921 | 20 | TTG | TAA | L |
| *L_copy2* | 10895 | 10956 | 62 | 37 | | | L |

| | | | | | | | |
|---|---|---|---|---|---|---|---|
| *rrnL* | 10957 | 11993 | 1037 | | | | L |
| trnV | 11994 | 12051 | 58 | | | | L |
| *rrnS* | 12052 | 12729 | 678 | | | | L |
| *control region* | 12730 | 13750 | 1021 | | | | H |
| *trnI* | 13751 | 13811 | 61 | | | | H |
| *trnM* | 13815 | 13875 | 61 | 3 | | | H |
| *trnC* | 13873 | 13930 | 58 | -3 | | | L |
| *trnY* | 13931 | 13991 | 61 | | | | L |
| *trnQ* | 13993 | 14044 | 52 | 1 | | | L |
| *nad2* | 14053 | 15046 | 994 | 8 | ATA | T | H |
| trnG | 15047 | 15109 | 63 | | | | H |
| *trnW* | 15110 | 15173 | 64 | | | | H |
| Overlap: | 14 | gap: | 8 | | | | |

***Stygobromus tenuis potomacus***

| Gene | Position | | Size | Intergenic nucleotides | Codon | | |
|---|---|---|---|---|---|---|---|
| | From | To | | | Start | Stop | Strand |
| *cox1* | 1 | 1534 | 1534 | | ATG | T | H |
| *trnL2* | 1535 | 1595 | 61 | | | | H |
| *cox2* | 1596 | 2270 | 675 | | GTG | TAA | H |
| *trnK* | 2271 | 2333 | 63 | | | | H |
| *trnD* | 2332 | 2393 | 62 | -2 | | | H |
| *atp8* | 2394 | 2582 | 189 | | ATT | TAG | H |
| *atp6* | 2542 | 3210 | 669 | -41 | ATG | TAA | H |
| cox3 | **3210** | 4004 | 795 | -1 | ATG | TAA | H |
| *nad3* | 4034 | 4387 | 354 | 29 | ATT | TAG | H |
| *trnA* | 4386 | 4444 | 59 | -2 | | | H |
| *trnS1* | 4445 | 4494 | 50 | | | | H |
| trnN | **4493** | 4554 | 62 | -2 | | | H |
| *trnE* | 4552 | 4612 | 61 | -3 | | | H |
| trnR | **4610** | 4666 | 57 | -3 | | | H |
| *trnF* | 4665 | 4724 | 60 | -2 | | | L |
| *nad5* | 4725 | 6432 | 1708 | | ATG | T | L |
| *trnH* | 6433 | 6490 | 58 | | | | L |
| *nad4* | 6491 | 7811 | 1321 | | ATG | T | L |
| *nad4L* | 7805 | 8098 | 294 | -7 | ATG | TAA | L |
| *trnT* | 8102 | 8160 | 59 | 3 | | | H |

| | | | | | | | |
|---|---|---|---|---|---|---|---|
| *trnP* | 8160 | 8219 | 60 | -1 | | | L |
| *nad6* | 8223 | 8723 | 501 | 3 | ATG | TAA | H |
| *cytb* | 8723 | 9862 | 1140 | -1 | ATG | TAA | H |
| *S_copy2* | 9861 | 9913 | 53 | -2 | | | H |
| *nad1* | 9935 | 10855 | 921 | 21 | TTG | TAG | L |
| *L_copy2* | 10891 | 10953 | 63 | 35 | | | L |
| rrnL | **10954** | 11989 | 1036 | | | | L |
| *trnV* | 11990 | 12047 | 58 | | | | L |
| *rrnS* | 12048 | 12726 | 679 | | | | L |
| *control region* | 12727 | 13282 | 556 | | | | H |
| trnI | **13283** | 13343 | 61 | | | | H |
| *trnM* | 13347 | 13407 | 61 | 3 | | | H |
| *trnC* | 13405 | 13462 | 58 | -3 | | | L |
| *trnY* | 13463 | 13523 | 61 | | | | L |
| *trnQ* | 13525 | 13576 | 52 | 1 | | | L |
| *nad2* | 13589 | 14582 | 994 | 12 | ATA | T | H |
| *trnG* | 14583 | 14645 | 63 | | | | H |
| trnW | 14646 | 14709 | 64 | | | | H |
| *Overlap:* | 13 | gap: | 8 | | | | |

***Stygobromus allegheniensis***

| **Gene** | **Position** | | **Size** | **Intergenic nucleotides** | **Codon** | | |
|---|---|---|---|---|---|---|---|
| | **From** | **To** | | | **Start** | **Stop** | **Strand** |
| cox1 | 1 | 1534 | 1534 | | ATC | T | H |
| *trnL2* | 1535 | 1596 | 62 | | | | H |
| *cox2* | 1597 | 2271 | 675 | | ATG | TAG | H |
| *trnK* | 2272 | 2334 | 63 | | | | H |
| *trnD* | 2333 | 2393 | 61 | -2 | | | H |
| *atp8* | 2403 | 2582 | 180 | 9 | ATA | TAA | H |
| *atp6* | 2542 | 3210 | 669 | -41 | ATG | TAA | H |
| *cox3* | 3210 | 4002 | 793 | -1 | ATG | T | H |
| nad3 | 4031 | 4384 | 354 | 28 | ATT | TAA | H |
| *trnA* | 4386 | 4444 | 59 | 1 | | | H |
| trnS1 | 4445 | 4495 | 51 | | | | H |
| *trnN* | 4494 | 4554 | 61 | -2 | | | H |
| trnE | 4556 | 4610 | 55 | 1 | | | H |
| *trnR* | 4611 | 4670 | 60 | | | | H |

| | | | | | | | |
|---|---|---|---|---|---|---|---|
| trnF | 4673 | 4732 | 60 | 2 | | | L |
| *nad5* | 4733 | 6440 | 1708 | | GTG | T | L |
| *trnH* | 6441 | 6501 | 61 | | | | L |
| nad4 | 6502 | 7822 | 1321 | | ATG | T | L |
| *nad4L* | 7816 | 8109 | 294 | -7 | ATG | TAG | L |
| *trnT* | 8113 | 8172 | 60 | 3 | | | H |
| *trnP* | 8172 | 8231 | 60 | -1 | | | L |
| nad6 | 8235 | 8735 | 501 | 3 | ATG | TAA | H |
| *cytb* | 8735 | 9874 | 1140 | -1 | ATG | TAA | H |
| S_copy2 | 9873 | 9926 | 54 | -2 | | | H |
| *nad1* | 9949 | 10869 | 921 | 22 | TTG | TAG | L |
| L_copy2 | 10904 | 10965 | 62 | 34 | | | L |
| *rrnL* | 10967 | 12000 | 1034 | 1 | | | L |
| trnV | 12002 | 12060 | 59 | 1 | | | L |
| *rrnS* | 12061 | 12748 | 688 | | | | L |
| *control region* | 12749 | 13739 | 991 | | | | H |
| trnI | 13740 | 13799 | 60 | | | | H |
| *trnM* | 13802 | 13862 | 61 | 2 | | | H |
| trnC | 13860 | 13916 | 57 | -3 | | | L |
| *trnY* | 13918 | 13976 | 59 | 1 | | | L |
| trnQ | 13979 | 14029 | 51 | 2 | | | L |
| *nad2* | 14041 | 14935 | 895 | 11 | ATA | T | H |
| *trnG* | 15035 | 15097 | 63 | 99 | | | H |
| trnW | 15098 | 15161 | 64 | | | | H |
| *Overlap:* | 9 | gap: | 16 | | | | |

***Crangonyx forbesi***

| **Gene** | **Position** | | **Size** | **Intergenic nucleotides** | **Codon** | | |
|---|---|---|---|---|---|---|---|
| | **From** | **To** | | | **Start** | **Stop** | **Strand** |
| *cox1* | 1 | 1534 | 1534 | | ATT | T | H |
| trnL2 | 1535 | 1595 | 61 | | | | H |
| *cox2* | 1596 | 2271 | 676 | | ATT | T | H |
| trnK | 2272 | 2330 | 59 | | | | H |
| *trnD* | 2337 | 2397 | 61 | 6 | | | H |
| *atp8* | 2401 | 2559 | 159 | 3 | ATG | TAA | H |
| *atp6* | 2564 | 3235 | 672 | 4 | ATG | TAA | H |
| *cox3* | 3235 | 4033 | 799 | -1 | ATG | T | H |

| | | | | | | | |
|---|---|---|---|---|---|---|---|
| *nad3* | 4031 | 4381 | 351 | -3 | ATT | TAG | H |
| *trnA* | 4380 | 4438 | 59 | -2 | | | H |
| *trnS1* | 4456 | 4507 | 52 | 17 | | | H |
| *trnN* | 4507 | 4568 | 62 | -1 | | | H |
| trnE | 4566 | 4631 | 66 | -3 | | | H |
| *trnR* | 4622 | 4672 | 51 | -10 | | | H |
| *trnF* | 4671 | 4730 | 60 | -2 | | | L |
| nad5 | 4734 | 6707 | 1974 | 3 | ATT | TAA | L |
| *nad6* | 6779 | 7273 | 495 | 71 | ATG | TAA | H |
| *cytb* | 7273 | 8418 | 1146 | -1 | ATG | TAA | H |
| S_copy2 | 8418 | 8467 | 50 | -1 | | | H |
| *trnH* | 8467 | 8526 | 60 | -1 | | | L |
| *nad4* | 8527 | 9838 | 1312 | | ATG | T | L |
| nad4L | 9847 | 10125 | 279 | 8 | ATA | TAA | L |
| *trnP* | 10153 | 10211 | 59 | 27 | | | L |
| *trnT* | 10214 | 10272 | 59 | 2 | | | H |
| nad1 | 10272 | 11195 | 924 | -1 | GTG | TAA | L |
| *control region* | 11198 | 12040 | 843 | 2 | | | H |
| *trnM* | 12041 | 12102 | 62 | | | | H |
| trnV | 12137 | 12194 | 58 | 34 | | | L |
| *nad2* | 12245 | 13231 | 987 | 50 | ATA | TAA | H |
| *trnY* | 13245 | 13307 | 63 | 13 | | | L |
| trnQ | 13304 | 13368 | 65 | -4 | | | L |
| *L_copy2* | 13372 | 13433 | 62 | 3 | | | L |
| rrnL | 13433 | 14522 | 1090 | -1 | | | L |
| *rrnS* | 14521 | 15215 | 695 | -2 | | | L |
| trnI | 15216 | 15274 | 59 | | | | H |
| *trnG* | 15278 | 15343 | 66 | 3 | | | H |
| trnC | 15345 | 15405 | 61 | 1 | | | L |
| *trnW* | 15408 | 15469 | 62 | 2 | | | H |
| Overlap: | 14 | gap: | 17 | | | | |

**Supplementary Table S2.** Summary of putative start codons in PCG of mitochondrial genomes of all amphipods

| | | Putative start codon | | | | | | | | | | | | |
|---|---|---|---|---|---|---|---|---|---|---|---|---|---|---|
| **Species** | **Accession** | ***atp6*** | ***atp8*** | ***cox1*** | ***cox2*** | ***cox3*** | ***cytb*** | ***nad1*** | ***nad2*** | ***nad3*** | ***nad4*** | ***nad4L*** | ***nad5*** | ***nad6*** |
| *Stygobromus pizzinii* | MN175620 | ATG | ATC | ATA | GTG | ATG | ATG | TTG | ATA | ATT | ATG | ATG | ATT | ATG |
| *Stygobromus tenuis potomacus* | MN175621 | ATG | ATT | ATG | GTG | ATG | ATG | TTG | ATA | ATT | ATG | ATG | ATG | ATG |
| *Bactrurus brachycaudus* | MN175619 | ATG | ATC | ATC | ATG | ATG | ATC | TTG | ATT | ATC | ATC | ATG | ATT | ATG |
| *Stygobromus allegheniensis* | MN175622 | ATG | GTA | ATC | ATG | ATG | ATG | TTG | ATA | ATT | ATG | ATG | GTG | ATG |
| *Crangonyx forbesi* | MN175623 | ATA | ATG | ATT | ATT | ATG | ATG | GTG | ATT | ATT | ATG | ATA | ATT | ATG |
| *Gondogeneia antarctica* | JN827386.1 | ATG | ATT | ATG | ATA | ATG | ATG | ATT | ATT | ATA | ATG | ATG | ATT | ATG |
| *Gmelinoides fasciatus* | NC_033361.1 | ATG | ATC | ATA | TTG | ATG | ATG | ATA | TTG | ATG | GTG | ATG | ATG | ATG |
| *Brachyuropus grewingkii* | NC_026309.1 | ATG | GTG | ATT | TTG | ATG | ATG | TTG | TTG | ATG | ATG | ATG | ATT | GTG |
| *Gammarus fossarum* | NC_034937.1 | ATA | ATC | ATA | TTG | ATG | ATA | TTG | TTG | ATG | ATA | ATG | TTG | ATG |
| *Pallaseopsis kessleri* | NC_033362.1 | ATG | ATT | ATT | GTG | ATG | ATG | TTG | TTG | ATG | TTG | ATG | TTG | ATT |
| *Gammarus duebeni* | NC_017760.1 | ATG | ATA | ATA | TTG | ATG | ATG | ATA | TTG | ATG | ATA | ATG | GTG | ATG |
| *Eulimnogammarus vittatus* | NC_025564.1 | ATG | GTG | ATT | TTG | ATG | ATG | TTG | TTG | ATG | ATG | ATG | TTG | ATG |
| *Caprella mutica* | NC_014492.1 | ATG | ATA | ATT | ATA | ATA | ATG | ATA | ATT | ATG | ATG | TTG | ATA | ATT |
| *Eulimnogammarus verrucosus* | NC_023104.1 | ATG | GTG | ATT | TTG | ATG | ATG | ATT | TTG | ATG | ATA | ATA | ATA | ATG |
| *Pseudoniphargus daviui* | NC_019662.2 | ATG | ATA | ATT | TTG | ATG | ATG | ATA | ATC | TTG | ATA | ATG | TTG | AAT |
| *Caprella scaura* | NC_014687.1 | ATG | ATA | ATT | ATA | ATG | ATG | CTG | ATG | ATG | ATG | CTG | ATA | ATA |
| *Metacrangonyx spinicaudatus* | NC_019657.1 | ATG | ATA | ATT | ATC | ATG | ATG | ATA | ATT | ATT | ATG | ATG | TTG | ATT |
| *Stygobromus tenuis potomacus* | KU869712.1 | ATG | ATT | ATT | ATC | ATG | ATG | TTG | ATA | ATA | ATG | ATG | ATA | ATG |
| *Metacrangonyx remyi* | NC_019660.1 | ATG | GTG | ATC | ATT | ATG | ATG | ATA | ATA | ATT | ATG | ATG | CTG | ATT |

| | | | | | | | | | | | | | | |
|---|---|---|---|---|---|---|---|---|---|---|---|---|---|---|
| *Platorchestia parapacifica* | MG010371.1 | ATA | ATG | ATT | ATT | ATG | ATA | ATT | ATC | ATG | ATG | ATT | ATA | ATC |
| *Platorchestia japonica* | MG010370.1 | ATA | ATG | ATT | ATA | ATG | ATA | ATT | ATA | ATG | ATG | ATA | ATA | ATT |
| *Metacrangonyx ilvanus* | NC_019656.1 | ATG | ATC | ATT | ATT | ATG | ATG | ATA | ATT | ATT | ATG | ATG | TTG | ATT |
| *Onisimus nanseni* | NC_013819.1 | ATG | ATT | ATG | TTG | ATG | ATG | ATA | ATG | ATG | TTG | ATG | GTG | ATT |
| *Metacrangonyx longicaudus* | NC_019658.1 | ATG | ATT | ATT | ATC | ATG | ATG | ATA | ATT | ATT | ATG | ATG | TTG | ATT |
| *Bahadzia jaraguensis* | FR872382.1 | ATG | ATC | ATT | ATT | ATG | ATG | ATG | ATT | ATT | ATG | ATG | ATA | ATT |
| *Stygobromus indentatus* | NC_030261.1 | ATA | ATT | ATT | ATA | ATG | ATG | GTG | ATT | ATA | ATT | ATG | GTG | ATG |
| *Metacrangonyx dominicanus* | NC_019654.1 | ATG | ATC | ATT | ATT | ATG | ATG | ATA | ATT | ATC | ATG | ATG | TTG | ATT |
| *Metacrangonyx goulmimensis* | NC_019655.1 | ATG | ATC | ATT | ATC | ATG | ATG | ATA | ATT | ATC | ATG | ATG | TTG | ATT |
| *Metacrangonyx panousei* | NC_019659.1 | ATG | ATC | ATT | ATT | ATG | ATG | ATA | ATT | ATT | ATG | ATG | TTG | ATT |
| *Eulimnogammarus cyaneus* | NC_033360.1 | ATG | GTG | ATC | TTG | ATG | ATG | TTG | TTG | ATG | ATG | ATG | GTG | ATG |
| *Metacrangonyx repens* | NC_019653.1 | ATG | GTG | ATT | ATT | ATG | ATG | ATA | ATT | ATT | ATG | ATG | TTG | ATT |
| *Metacrangonyx longipes* | NC_013032.1 | ATG | ATC | ATT | ATT | ATG | ATG | ATA | ATT | ATT | ATA | ATA | ATT | ATT |
| *Metacrangonyx sp. 3 ssp. 1 MDMBR-2012* | HE860504.1 | ATG | ATC | ATT | ATT | ATG | ATG | ATA | ATT | ATT | ATG | ATG | TTG | ATT |
| *Metacrangonyx sp. 4 MDMBR-2012* | HE860498.1 | ATG | ATC | ATC | ATC | ATG | ATG | ATA | ATA | ATC | GTG | ATG | TTG | ATT |
| *Metacrangonyx sp. 1 MDMBR-2012* | HE860513.1 | ATG | ATC | ATT | ATC | ATG | ATG | ATA | TTT | ATT | ATG | ATG | TTG | ATT |

**Supplementary Table S3.** Summary of codon usage in PCG of mitochondrial genomes of all crangonyctid amphipods. Sign "#" indicates total number of certain codon in protein-coding sequences of every species; "%" indicates percent of certain codon in total coding sequence in every species; RSCU indicates the calculated RSCU value of certain codon in total coding sequence in every species

| AA | Codon | Stygobromus indentatus | | | Crangonyx forbesi | | | Stygobromus allegheniensis | | | Stygobromus tenuis potomacus | | | Stygobromus pizzinii | | | Bactrurus brachycaudus | | | Stygobromus tenuis potomacus | | |
|---|---|---|---|---|---|---|---|---|---|---|---|---|---|---|---|---|---|---|---|---|---|---|
| | | RSCU | # | % | RSCU | # | % | RSCU | # | % | RSCU | # | % | RSCU | # | % | RSCU | # | % | RSCU | # | % |
| Phe | UUU | 1.74 | 238 | 6.45 | 1.76 | 255 | 6.78 | 1.64 | 237 | 6.49 | 1.70 | 240 | 6.51 | 1.69 | 241 | 6.54 | 1.55 | 218 | 5.94 | 1.72 | 243 | 6.57 |
| | UUC | 0.26 | 35 | 0.95 | 0.24 | 35 | 0.93 | 0.36 | 52 | 1.42 | 0.30 | 42 | 1.14 | 0.31 | 45 | 1.22 | 0.45 | 63 | 1.72 | 0.28 | 39 | 1.05 |
| Leu2 | UUA | 2.89 | 286 | 7.75 | 2.58 | 267 | 7.10 | 2.56 | 259 | 7.09 | 2.95 | 298 | 8.08 | 2.96 | 297 | 8.06 | 2.10 | 207 | 5.64 | 3.04 | 314 | 8.49 |
| | UUG | 0.49 | 48 | 1.30 | 0.64 | 66 | 1.76 | 0.89 | 90 | 2.46 | 0.64 | 65 | 1.76 | 0.58 | 58 | 1.57 | 0.75 | 74 | 2.02 | 0.54 | 56 | 1.51 |
| Leu1 | CUU | 1.00 | 99 | 2.68 | 1.05 | 109 | 2.90 | 0.97 | 98 | 2.68 | 1.02 | 103 | 2.79 | 1.03 | 103 | 2.79 | 1.18 | 116 | 3.16 | 0.99 | 102 | 2.76 |
| | CUC | 0.33 | 33 | 0.89 | 0.32 | 33 | 0.88 | 0.27 | 27 | 0.74 | 0.26 | 26 | 0.70 | 0.24 | 24 | 0.65 | 0.46 | 45 | 1.23 | 0.27 | 28 | 0.76 |
| | CUA | 1.10 | 109 | 2.95 | 1.07 | 111 | 2.95 | 0.99 | 100 | 2.74 | 0.88 | 89 | 2.41 | 0.97 | 97 | 2.63 | 1.12 | 110 | 3.00 | 0.91 | 94 | 2.54 |
| | CUG | 0.18 | 18 | 0.49 | 0.33 | 34 | 0.90 | 0.33 | 33 | 0.90 | 0.25 | 25 | 0.68 | 0.23 | 23 | 0.62 | 0.40 | 39 | 1.06 | 0.24 | 25 | 0.68 |
| Ile | AUU | 1.57 | 253 | 6.85 | 1.47 | 185 | 4.92 | 1.45 | 188 | 5.15 | 1.62 | 240 | 6.51 | 1.61 | 245 | 6.64 | 1.48 | 201 | 5.48 | 1.53 | 230 | 6.22 |
| | AUC | 0.43 | 70 | 1.90 | 0.53 | 66 | 1.76 | 0.55 | 71 | 1.94 | 0.38 | 56 | 1.52 | 0.39 | 59 | 1.60 | 0.52 | 70 | 1.91 | 0.47 | 71 | 1.92 |
| Met | AUA | 1.71 | 197 | 5.34 | 1.61 | 183 | 4.87 | 1.66 | 151 | 4.13 | 1.64 | 169 | 4.58 | 1.66 | 169 | 4.58 | 1.37 | 153 | 4.17 | 1.70 | 168 | 4.54 |
| | AUG | 0.29 | 33 | 0.89 | 0.39 | 45 | 1.20 | 0.34 | 31 | 0.85 | 0.36 | 37 | 1.00 | 0.34 | 35 | 0.95 | 0.63 | 71 | 1.94 | 0.30 | 30 | 0.81 |
| Val | GUU | 1.71 | 106 | 2.87 | 1.61 | 112 | 2.98 | 1.55 | 111 | 3.04 | 1.72 | 118 | 3.20 | 1.57 | 101 | 2.74 | 1.36 | 96 | 2.62 | 1.60 | 103 | 2.79 |
| | GUC | 0.32 | 20 | 0.54 | 0.46 | 32 | 0.85 | 0.50 | 36 | 0.99 | 0.42 | 29 | 0.79 | 0.50 | 32 | 0.87 | 0.54 | 38 | 1.04 | 0.34 | 22 | 0.60 |
| | GUA | 1.24 | 77 | 2.09 | 1.38 | 96 | 2.55 | 1.17 | 84 | 2.30 | 1.39 | 95 | 2.58 | 1.40 | 90 | 2.44 | 1.52 | 107 | 2.92 | 1.61 | 104 | 2.81 |
| | GUG | 0.73 | 45 | 1.22 | 0.56 | 39 | 1.04 | 0.78 | 56 | 1.53 | 0.47 | 32 | 0.87 | 0.54 | 35 | 0.95 | 0.58 | 41 | 1.12 | 0.45 | 29 | 0.78 |
| Ser2 | UCU | 2.01 | 108 | 2.93 | 1.93 | 110 | 2.93 | 1.81 | 92 | 2.52 | 1.80 | 94 | 2.55 | 1.89 | 100 | 2.71 | 1.74 | 95 | 2.59 | 1.95 | 100 | 2.70 |
| | UCC | 0.56 | 30 | 0.81 | 0.47 | 27 | 0.72 | 0.47 | 24 | 0.66 | 0.63 | 33 | 0.89 | 0.43 | 23 | 0.62 | 0.75 | 41 | 1.12 | 0.25 | 13 | 0.35 |
| | UCA | 1.34 | 72 | 1.95 | 1.39 | 79 | 2.10 | 1.38 | 70 | 1.92 | 1.45 | 76 | 2.06 | 1.61 | 85 | 2.31 | 1.37 | 75 | 2.04 | 1.74 | 89 | 2.41 |
| | UCG | 0.17 | 9 | 0.24 | 0.39 | 22 | 0.59 | 0.18 | 9 | 0.25 | 0.19 | 10 | 0.27 | 0.09 | 5 | 0.14 | 0.31 | 17 | 0.46 | 0.14 | 7 | 0.19 |
| Pro | CCU | 1.88 | 69 | 1.87 | 1.87 | 71 | 1.89 | 1.95 | 71 | 1.94 | 1.66 | 62 | 1.68 | 1.72 | 64 | 1.74 | 1.76 | 65 | 1.77 | 1.79 | 67 | 1.81 |
| | CCC | 0.54 | 20 | 0.54 | 0.84 | 32 | 0.85 | 0.44 | 16 | 0.44 | 0.27 | 10 | 0.27 | 0.56 | 21 | 0.57 | 0.65 | 24 | 0.65 | 0.61 | 23 | 0.62 |

| | | | | | | | | | | | | | | | | | | | | | | |
|---|---|---|---|---|---|---|---|---|---|---|---|---|---|---|---|---|---|---|---|---|---|---|
| | CCA | 1.31 | 48 | 1.30 | 0.84 | 32 | 0.85 | 1.37 | 50 | 1.37 | 1.80 | 67 | 1.82 | 1.40 | 52 | 1.41 | 1.11 | 41 | 1.12 | 1.44 | 54 | 1.46 |
| | CCG | 0.27 | 10 | 0.27 | 0.45 | 17 | 0.45 | 0.25 | 9 | 0.25 | 0.27 | 10 | 0.27 | 0.32 | 12 | 0.33 | 0.49 | 18 | 0.49 | 0.16 | 6 | 0.16 |
| Thr | ACU | 1.59 | 70 | 1.90 | 1.42 | 74 | 1.97 | 1.81 | 83 | 2.27 | 1.70 | 75 | 2.03 | 1.76 | 79 | 2.14 | 1.37 | 56 | 1.53 | 1.67 | 79 | 2.14 |
| | ACC | 0.59 | 26 | 0.70 | 0.73 | 38 | 1.01 | 0.52 | 24 | 0.66 | 0.59 | 26 | 0.70 | 0.47 | 21 | 0.57 | 0.88 | 36 | 0.98 | 0.68 | 32 | 0.87 |
| | ACA | 1.73 | 76 | 2.06 | 1.52 | 79 | 2.10 | 1.51 | 69 | 1.89 | 1.61 | 71 | 1.93 | 1.67 | 75 | 2.03 | 1.37 | 56 | 1.53 | 1.54 | 73 | 1.97 |
| | ACG | 0.09 | 4 | 0.11 | 0.33 | 17 | 0.45 | 0.15 | 7 | 0.19 | 0.09 | 4 | 0.11 | 0.11 | 5 | 0.14 | 0.37 | 15 | 0.41 | 0.11 | 5 | 0.14 |
| Ala | GCU | 2.05 | 106 | 2.87 | 1.66 | 72 | 1.92 | 1.86 | 103 | 2.82 | 2.08 | 108 | 2.93 | 1.92 | 101 | 2.74 | 1.77 | 105 | 2.86 | 2.02 | 109 | 2.95 |
| | GCC | 0.77 | 40 | 1.08 | 0.90 | 39 | 1.04 | 0.89 | 49 | 1.34 | 0.67 | 35 | 0.95 | 0.86 | 45 | 1.22 | 1.08 | 64 | 1.74 | 0.72 | 39 | 1.05 |
| | GCA | 1.10 | 57 | 1.54 | 1.13 | 49 | 1.30 | 0.98 | 54 | 1.48 | 1.13 | 59 | 1.60 | 0.95 | 50 | 1.36 | 0.81 | 48 | 1.31 | 1.11 | 60 | 1.62 |
| | GCG | 0.08 | 4 | 0.11 | 0.30 | 13 | 0.35 | 0.27 | 15 | 0.41 | 0.12 | 6 | 0.16 | 0.27 | 14 | 0.38 | 0.34 | 20 | 0.55 | 0.15 | 8 | 0.22 |
| Tyr | UAU | 1.46 | 100 | 2.71 | 1.11 | 83 | 2.21 | 1.31 | 88 | 2.41 | 1.49 | 102 | 2.77 | 1.32 | 89 | 2.41 | 1.35 | 75 | 2.04 | 1.42 | 97 | 2.62 |
| | UAC | 0.54 | 37 | 1.00 | 0.89 | 66 | 1.76 | 0.69 | 46 | 1.26 | 0.51 | 35 | 0.95 | 0.68 | 46 | 1.25 | 0.65 | 36 | 0.98 | 0.58 | 40 | 1.08 |
| His | CAU | 1.37 | 54 | 1.46 | 0.99 | 36 | 0.96 | 1.21 | 47 | 1.29 | 1.30 | 46 | 1.25 | 1.32 | 49 | 1.33 | 0.94 | 33 | 0.90 | 1.28 | 46 | 1.24 |
| | CAC | 0.63 | 25 | 0.68 | 1.01 | 37 | 0.98 | 0.79 | 31 | 0.85 | 0.70 | 25 | 0.68 | 0.68 | 25 | 0.68 | 1.06 | 37 | 1.01 | 0.72 | 26 | 0.70 |
| Gln | CAA | 1.70 | 51 | 1.38 | 1.19 | 34 | 0.90 | 1.20 | 42 | 1.15 | 1.42 | 46 | 1.25 | 1.44 | 46 | 1.25 | 1.63 | 53 | 1.44 | 1.31 | 42 | 1.14 |
| | CAG | 0.30 | 9 | 0.24 | 0.81 | 23 | 0.61 | 0.80 | 28 | 0.77 | 0.58 | 19 | 0.52 | 0.56 | 18 | 0.49 | 0.37 | 12 | 0.33 | 0.69 | 22 | 0.60 |
| Asn | AAU | 1.31 | 81 | 2.19 | 1.19 | 80 | 2.13 | 1.25 | 74 | 2.03 | 1.40 | 96 | 2.60 | 1.37 | 90 | 2.44 | 1.15 | 61 | 1.66 | 1.39 | 91 | 2.46 |
| | AAC | 0.69 | 43 | 1.16 | 0.81 | 55 | 1.46 | 0.75 | 44 | 1.20 | 0.60 | 41 | 1.11 | 0.63 | 41 | 1.11 | 0.85 | 45 | 1.23 | 0.61 | 40 | 1.08 |
| Lys | AAA | 1.63 | 79 | 2.14 | 1.60 | 100 | 2.66 | 1.60 | 71 | 1.94 | 1.47 | 69 | 1.87 | 1.47 | 69 | 1.87 | 1.25 | 57 | 1.55 | 1.60 | 76 | 2.06 |
| | AAG | 0.37 | 18 | 0.49 | 0.40 | 25 | 0.67 | 0.40 | 18 | 0.49 | 0.53 | 25 | 0.68 | 0.53 | 25 | 0.68 | 0.75 | 34 | 0.93 | 0.40 | 19 | 0.51 |
| Asp | GAU | 1.24 | 44 | 1.19 | 1.17 | 42 | 1.12 | 1.13 | 44 | 1.20 | 1.41 | 52 | 1.41 | 1.30 | 48 | 1.30 | 1.32 | 45 | 1.23 | 1.39 | 52 | 1.41 |
| | GAC | 0.76 | 27 | 0.73 | 0.83 | 30 | 0.80 | 0.87 | 34 | 0.93 | 0.59 | 22 | 0.60 | 0.70 | 26 | 0.71 | 0.68 | 23 | 0.63 | 0.61 | 23 | 0.62 |
| Glu | GAA | 1.35 | 58 | 1.57 | 1.01 | 37 | 0.98 | 1.31 | 53 | 1.45 | 1.24 | 51 | 1.38 | 1.33 | 55 | 1.49 | 1.09 | 44 | 1.20 | 1.40 | 58 | 1.57 |
| | GAG | 0.65 | 28 | 0.76 | 0.99 | 36 | 0.96 | 0.69 | 28 | 0.77 | 0.76 | 31 | 0.84 | 0.67 | 28 | 0.76 | 0.91 | 37 | 1.01 | 0.60 | 25 | 0.68 |
| Cys | UGU | 1.62 | 30 | 0.81 | 1.59 | 35 | 0.93 | 1.25 | 25 | 0.68 | 1.49 | 26 | 0.70 | 1.60 | 28 | 0.76 | 1.33 | 24 | 0.65 | 1.56 | 28 | 0.76 |
| | UGC | 0.38 | 7 | 0.19 | 0.41 | 9 | 0.24 | 0.75 | 15 | 0.41 | 0.51 | 9 | 0.24 | 0.40 | 7 | 0.19 | 0.67 | 12 | 0.33 | 0.44 | 8 | 0.22 |
| Trp | UGA | 1.53 | 74 | 2.00 | 1.29 | 65 | 1.73 | 1.28 | 59 | 1.62 | 1.30 | 60 | 1.63 | 1.40 | 65 | 1.76 | 1.10 | 55 | 1.50 | 1.61 | 75 | 2.03 |
| | UGG | 0.47 | 23 | 0.62 | 0.71 | 36 | 0.96 | 0.72 | 33 | 0.90 | 0.70 | 32 | 0.87 | 0.60 | 28 | 0.76 | 0.90 | 45 | 1.23 | 0.39 | 18 | 0.49 |
| Arg | CGU | 1.36 | 17 | 0.46 | 1.11 | 15 | 0.40 | 1.46 | 19 | 0.52 | 1.54 | 20 | 0.54 | 1.23 | 16 | 0.43 | 0.64 | 9 | 0.25 | 1.23 | 16 | 0.43 |

| | | | | | | | | | | | | | | | | | | | | | | |
|---|---|---|---|---|---|---|---|---|---|---|---|---|---|---|---|---|---|---|---|---|---|---|
| | CGC | 0.32 | 4 | 0.11 | 0.44 | 6 | 0.16 | 0.54 | 7 | 0.19 | 0.38 | 5 | 0.14 | 0.69 | 9 | 0.24 | 1.00 | 14 | 0.38 | 0.54 | 7 | 0.19 |
| | CGA | 1.76 | 22 | 0.60 | 1.19 | 16 | 0.43 | 1.08 | 14 | 0.38 | 1.38 | 18 | 0.49 | 1.31 | 17 | 0.46 | 1.29 | 18 | 0.49 | 1.85 | 24 | 0.65 |
| | CGG | 0.56 | 7 | 0.19 | 1.26 | 17 | 0.45 | 0.92 | 12 | 0.33 | 0.69 | 9 | 0.24 | 0.77 | 10 | 0.27 | 1.07 | 15 | 0.41 | 0.38 | 5 | 0.14 |
| Ser1 | AGU | 1.10 | 59 | 1.60 | 1.18 | 67 | 1.78 | 1.08 | 55 | 1.51 | 1.09 | 57 | 1.55 | 1.10 | 58 | 1.57 | 0.75 | 41 | 1.12 | 0.96 | 49 | 1.33 |
| | AGC | 0.21 | 11 | 0.30 | 0.39 | 22 | 0.59 | 0.41 | 21 | 0.58 | 0.23 | 12 | 0.33 | 0.38 | 20 | 0.54 | 0.46 | 25 | 0.68 | 0.21 | 11 | 0.30 |
| | AGA | 1.86 | 100 | 2.71 | 1.47 | 84 | 2.23 | 1.97 | 100 | 2.74 | 2.03 | 106 | 2.87 | 1.87 | 99 | 2.69 | 1.59 | 87 | 2.37 | 1.85 | 95 | 2.57 |
| | AGG | 0.75 | 40 | 1.08 | 0.79 | 45 | 1.20 | 0.69 | 35 | 0.96 | 0.57 | 30 | 0.81 | 0.62 | 33 | 0.90 | 1.03 | 56 | 1.53 | 0.90 | 46 | 1.24 |
| Gly | GGU | 1.02 | 58 | 1.57 | 1.11 | 61 | 1.62 | 1.12 | 67 | 1.83 | 1.06 | 62 | 1.68 | 1.08 | 64 | 1.74 | 0.86 | 54 | 1.47 | 1.07 | 63 | 1.70 |
| | GGC | 0.53 | 30 | 0.81 | 0.82 | 45 | 1.20 | 0.43 | 26 | 0.71 | 0.43 | 25 | 0.68 | 0.54 | 32 | 0.87 | 0.83 | 52 | 1.42 | 0.39 | 23 | 0.62 |
| | GGA | 1.44 | 82 | 2.22 | 1.11 | 61 | 1.62 | 1.17 | 70 | 1.92 | 1.57 | 92 | 2.49 | 1.29 | 76 | 2.06 | 0.91 | 57 | 1.55 | 1.47 | 87 | 2.35 |
| | GGG | 1.00 | 57 | 1.54 | 0.95 | 52 | 1.38 | 1.28 | 77 | 2.11 | 0.94 | 55 | 1.49 | 1.08 | 64 | 1.74 | 1.39 | 87 | 2.37 | 1.07 | 63 | 1.70 |

**Supplementary Table S4.** Summary of amino acid composition in PCGs of mitochondrial genomes of all crangonyctid amphipods. Sign “#” indicates total number of certain amino acid in protein-coding sequences of every species; “%” indicates percent of certain amino acid in total coding sequence in every species

| AA | Stygobromus tenuis potomacus | | Stygobromus indentatus | | Stygobromus allegheniensis | | Bactrurus brachycaudus | | Crangonyx forbesi | | Stygobromus pizzinii | | Stygobromus tenuis potomacus | |
|---|---|---|---|---|---|---|---|---|---|---|---|---|---|---|
| | # | % | # | % | # | % | # | % | # | % | # | % | # | % |
| Phe | 282 | 7.63 | 273 | 7.40 | 289 | 7.91 | 281 | 7.66 | 290 | 7.71 | 286 | 7.76 | 282 | 7.65 |
| Leu2 | 370 | 10.01 | 334 | 9.05 | 349 | 9.56 | 281 | 7.66 | 333 | 8.86 | 355 | 9.63 | 363 | 9.84 |
| Leu1 | 249 | 6.74 | 259 | 7.02 | 258 | 7.06 | 310 | 8.45 | 287 | 7.64 | 247 | 6.70 | 243 | 6.59 |
| Ile | 301 | 8.14 | 323 | 8.75 | 259 | 7.09 | 271 | 7.39 | 251 | 6.68 | 304 | 8.25 | 296 | 8.03 |
| Met | 198 | 5.36 | 230 | 6.23 | 182 | 4.98 | 224 | 6.11 | 228 | 6.07 | 204 | 5.53 | 206 | 5.59 |
| Val | 258 | 6.98 | 248 | 6.72 | 287 | 7.86 | 282 | 7.69 | 279 | 7.42 | 258 | 7.00 | 274 | 7.43 |
| Ser2 | 209 | 5.65 | 219 | 5.93 | 195 | 5.34 | 228 | 6.22 | 238 | 6.33 | 213 | 5.78 | 213 | 5.78 |
| Pro | 150 | 4.06 | 147 | 3.98 | 146 | 4.00 | 148 | 4.03 | 152 | 4.04 | 149 | 4.04 | 149 | 4.04 |
| Thr | 189 | 5.11 | 176 | 4.77 | 183 | 5.01 | 163 | 4.44 | 208 | 5.53 | 180 | 4.88 | 176 | 4.77 |
| Ala | 216 | 5.84 | 207 | 5.61 | 221 | 6.05 | 237 | 6.46 | 173 | 4.60 | 210 | 5.70 | 208 | 5.64 |
| Tyr | 137 | 3.71 | 137 | 3.71 | 134 | 3.67 | 111 | 3.03 | 149 | 3.96 | 135 | 3.66 | 137 | 3.71 |

| | | | | | | | | | | | | | | |
|---|---|---|---|---|---|---|---|---|---|---|---|---|---|---|
| His | 72 | 1.95 | 79 | 2.14 | 78 | 2.14 | 70 | 1.91 | 73 | 1.94 | 74 | 2.01 | 71 | 1.93 |
| Gln | 64 | 1.73 | 60 | 1.63 | 70 | 1.92 | 65 | 1.77 | 57 | 1.52 | 64 | 1.74 | 65 | 1.76 |
| Asn | 131 | 3.54 | 124 | 3.36 | 118 | 3.23 | 106 | 2.89 | 135 | 3.59 | 131 | 3.55 | 137 | 3.71 |
| Lys | 95 | 2.57 | 97 | 2.63 | 89 | 2.44 | 91 | 2.48 | 125 | 3.33 | 94 | 2.55 | 94 | 2.55 |
| Asp | 75 | 2.03 | 71 | 1.92 | 78 | 2.14 | 68 | 1.85 | 72 | 1.92 | 74 | 2.01 | 74 | 2.01 |
| Glu | 83 | 2.25 | 86 | 2.33 | 81 | 2.22 | 81 | 2.21 | 73 | 1.94 | 83 | 2.25 | 82 | 2.22 |
| Cys | 36 | 0.97 | 37 | 1.00 | 40 | 1.10 | 36 | 0.98 | 44 | 1.17 | 35 | 0.95 | 35 | 0.95 |
| Trp | 93 | 2.52 | 97 | 2.63 | 92 | 2.52 | 100 | 2.73 | 101 | 2.69 | 93 | 2.52 | 92 | 2.49 |
| Arg | 52 | 1.41 | 50 | 1.35 | 52 | 1.42 | 56 | 1.53 | 54 | 1.44 | 52 | 1.41 | 52 | 1.41 |
| Ser1 | 201 | 5.44 | 210 | 5.69 | 211 | 5.78 | 209 | 5.70 | 218 | 5.80 | 210 | 5.70 | 205 | 5.56 |
| Gly | 236 | 6.38 | 227 | 6.15 | 240 | 6.57 | 250 | 6.82 | 219 | 5.83 | 236 | 6.40 | 234 | 6.34 |

**Supplementary Table S5.** Nucleotide composition statistics for ribosomal RNA genes in mitochondrial genomes of all crangonyctid amphipods.

| Species | Gene | Size (bp) | AT % | AT-skew | GC-skew |
|---|---|---|---|---|---|
| | *rrnL-* | 1034 | 67.8 | -0.030 | 0.381 |
| *Bactrurus brachycaudus* | *rrnS-* | 671 | 71.5 | -0.029 | 0.361 |
| | *rrnL-* | 1090 | 72.7 | -0.064 | 0.259 |
| *Crangonyx forbesi* | *rrnS-* | 695 | 73.6 | -0.084 | 0.359 |
| | *rrnL-* | 1034 | 70.4 | -0.115 | 0.412 |
| *Stygobromus allegheniensis* | *rrnS-* | 688 | 74.0 | -0.108 | 0.352 |
| | *rrnL-* | 1035 | 72.7 | -0.089 | 0.383 |
| *Stygobromus indentatus* | *rrnS-* | 669 | 77.2 | -0.070 | 0.307 |
| | *rrnL-* | 1037 | 72.3 | -0.093 | 0.415 |
| *Stygobromus pizzinii* | *rrnS-* | 678 | 74.6 | -0.075 | 0.337 |
| | *rrnL-* | 1036 | 72.8 | -0.074 | 0.426 |
| *Stygobromus tenuis potomacus* | *rrnS-* | 679 | 76.0 | -0.109 | 0.370 |
| | *rrnL-* | 1034 | 72.3 | -0.099 | 0.420 |
| *Stygobromus tenuis potomacus* | *rrnS-* | 683 | 74.6 | -0.088 | 0.322 |

**Supplementary Table S6.** Results of selective pressure (ω ratio) analyses of mitochondrial PCGs with LRT P-value < 0.05 in subterranean and surface lineages of amphipods based on 2 vs. 1 ratio model.

| Model | np | Ln L | Estimates of parameters | | | Model compared | LRT P-value | Omega for Foreground Branch | Gene | Species |
|---|---|---|---|---|---|---|---|---|---|---|
| Two ratio Model 2 | 70 | -24608.883248 | ω: | ω0=0.02037 | ω1=0.05379 | Model 0 vs. Two ratio Model 2 | 0.000131407 | ω1=0.05379 | *cox1* | *C. forbesi* |
| Model 0 | 69 | -24616.193996 | ω= | 0.02099 | | | | | | |
| Two ratio Model 2 | 70 | -15283.383385 | ω: | ω0=0.05091 | ω1=0.18347 | Model 0 vs. Two ratio Model 2 | 0.037178941 | ω1=0.18347 | *atp6* | *P. daviui* |
| Model 0 | 69 | -15285.554483 | ω= | 0.05174 | | | | | | |
| Two ratio Model 2 | 70 | -24611.896377 | ω: | ω0=0.02052 | ω1=0.04222 | Model 0 vs. Two ratio Model 2 | 0.003370432 | ω1=0.04222 | *cox1* | |
| Model 0 | 69 | -24616.193996 | ω= | 0.02099 | | | | | | |
| Two ratio Model 2 | 70 | -19414.614776 | ω: | ω0=0.03225 | ω1=0.19046 | Model 0 vs. Two ratio Model 2 | 0.002904709 | ω1=0.19046 | *nad1* | |
| Model 0 | 69 | -19419.047965 | ω= | 0.03239 | | | | | | |
| Two ratio Model 2 | 70 | -24599.610909 | ω: | ω0=0.02012 | ω1=0.07488 | Model 0 vs. Two ratio Model 2 | 0.000000008 | ω1=0.07488 | *cox1* | *G. fasciatus* |
| Model 0 | 69 | -24616.193996 | ω= | 0.02099 | | | | | | |
| Two ratio Model 2 | 70 | -12953.332620 | ω: | ω0=0.03009 | ω1=0.21013 | Model 0 vs. Two ratio Model 2 | 0.000072299 | ω1=0.21013 | *cox2* | |
| Model 0 | 69 | -12961.207473 | ω= | 0.03033 | | | | | | |
| Two ratio Model 2 | 70 | -16204.441186 | ω: | ω0=0.04169 | ω1=0.09747 | Model 0 vs. Two ratio Model 2 | 0.012180256 | ω1=0.09747 | *cox3* | |
| Model 0 | 69 | -16207.583406 | ω= | 0.04274 | | | | | | |
| Two ratio Model 2 | 70 | -29486.407012 | ω: | ω0=0.04262 | ω1=0.12215 | Model 0 vs. Two ratio Model 2 | 0.002680018 | ω1=0.12215 | *nad4* | |
| Model 0 | 69 | -29490.913731 | ω= | 0.04296 | | | | | | |
| Two ratio Model 2 | 70 | -16205.223658 | ω: | ω0=0.04182 | ω1=0.10051 | Model 0 vs. Two ratio Model 2 | 0.029822483 | ω1=0.10051 | *cox3* | *O. nanseni* |
| Model 0 | 69 | -16207.583406 | ω= | 0.04274 | | | | | | |
| Two ratio Model 2 | 70 | -24619.041265 | ω: | ω0=0.01766 | ω1=0.04082 | Model 0 vs. Two ratio Model 2 | 0.017017792 | ω1=0.04082 | *cox1* | *G. fossarum* |
| Model 0 | 69 | -24616.193996 | ω= | 0.02099 | | | | | | |
| Two ratio Model 2 | 70 | -23400.542887 | ω: | ω0=0.03254 | ω1=0.08246 | Model 0 vs. Two ratio Model 2 | 0.004931290 | ω1=0.08246 | *cytb* | |
| Model 0 | 69 | -23404.495120 | ω= | 0.03337 | | | | | | |
| Two ratio Model 2 | 70 | -19414.218221 | ω: | ω0=0.03259 | ω1=0.13042 | Model 0 vs. Two ratio Model 2 | 0.001883761 | ω1=0.13042 | *nad1* | |
| Model 0 | 69 | -19419.047965 | ω= | 0.03239 | | | | | | |

| | | | | | | | | | | |
|---|---|---|---|---|---|---|---|---|---|---|
| Two ratio Model 2 | 70 | -29487.580618 | ω: | ω0=0.04235 | ω1=0.09448 | Model 0 vs. Two ratio Model 2 | 0.009825704 | ω1=0.09448 | *nad4* | |
| Model 0 | 69 | -29490.913731 | ω= | 0.04296 | | | | | | |
| Two ratio Model 2 | 70 | -6864.685694 | ω: | ω0=0.03733 | ω1=0.19416 | Model 0 vs. Two ratio Model 2 | 0.030276845 | ω1=0.19416 | *nad4l* | |
| Model 0 | 69 | -6867.032446 | ω= | 0.03788 | | | | | | |
| Two ratio Model 2 | 70 | -24613.676186 | ω: | ω0=0.02143 | ω1=0.01086 | Model 0 vs. Two ratio Model 2 | 0.024831197 | ω1=0.01086 | *cox1* | *B. jaraguensis* |
| Model 0 | 69 | -24616.193996 | ω= | 0.02099 | | | | | | |
| Two ratio Model 2 | 70 | -42645.830655 | ω: | ω0=0.06114 | ω1=0.02138 | Model 0 vs. Two ratio Model 2 | 0.035443809 | ω1=0.02138 | *nad5* | |
| Model 0 | 69 | -42648.042488 | ω= | 0.06077 | | | | | | |
| Two ratio Model 2 | 70 | -24610.442091 | ω: | ω0=0.02042 | ω1=0.04738 | Model 0 vs. Two ratio Model 2 | 0.000694537 | ω1=0.04738 | *cox1* | *B. brachycaudus* |
| Model 0 | 69 | -24616.193996 | ω= | 0.02099 | | | | | | |
| Two ratio Model 2 | 70 | -16205.427054 | ω: | ω0=0.04211 | ω1=0.09064 | Model 0 vs. Two ratio Model 2 | 0.037828788 | ω1=0.09064 | *cox3* | |
| Model 0 | 69 | -16207.583406 | ω= | 0.04274 | | | | | | |
| Two ratio Model 2 | 70 | -19416.309168 | ω: | ω0=0.03286 | ω1=0.00207 | Model 0 vs. Two ratio Model 2 | 0.019261754 | ω1=0.00207 | *nad1* | |
| Model 0 | 69 | -19419.047965 | ω= | 0.03239 | | | | | | |
| Two ratio Model 2 | 70 | -24607.788487 | ω: | ω0=0.02132 | ω1=0.00010 | Model 0 vs. Two ratio Model 2 | 0.000041293 | ω1=0.00010 | *cox1* | *S. tenuis_MN* |
| Model 0 | 69 | -24616.193996 | ω= | 0.02099 | | | | | | |
| Two ratio Model 2 | 70 | -12967.565600 | ω: | ω0=0.02987 | ω1=2.22100 | Model 0 vs. Two ratio Model 2 | 0.000362491 | ω1=2.22100 | *cox2* | |
| Model 0 | 69 | -12961.207473 | ω= | 0.03033 | | | | | | |
| Two ratio Model 2 | 70 | -19417.045819 | ω: | ω0=0.03302 | ω1=0.00531 | Model 0 vs. Two ratio Model 2 | 0.045384555 | ω1=0.00531 | *nad1* | |
| Model 0 | 69 | -19419.047965 | ω= | 0.03239 | | | | | | |
| Two ratio Model 2 | 70 | -6864.477274 | ω: | ω0=0.03988 | ω1=0.00010 | Model 0 vs. Two ratio Model 2 | 0.023783604 | ω1=0.00010 | *nad4l* | |
| Model 0 | 69 | -6867.032446 | ω= | 0.03788 | | | | | | |
| Two ratio Model 2 | 70 | -42645.097762 | ω: | ω0=0.06190 | ω1=0.02375 | Model 0 vs. Two ratio Model 2 | 0.015231840 | ω1=0.02375 | *nad5* | |
| Model 0 | 69 | -42648.042488 | ω= | 0.06077 | | | | | | |
| Two ratio Model 2 | 70 | -24643.013443 | ω: | ω0=0.02059 | ω1=37.72058 | Model 0 vs. Two ratio Model 2 | 0.000000000 | ω1=37.72058 | *cox1* | *S. tenuis_KU* |
| Model 0 | 69 | -24616.193996 | ω= | 0.02099 | | | | | | |
| Two ratio Model 2 | 70 | -16204.089657 | ω: | ω0=0.04368 | ω1=0.01396 | Model 0 vs. Two ratio Model 2 | 0.008208101 | ω1=0.01396 | *cox3* | |
| Model 0 | 69 | -16207.583406 | ω= | 0.04274 | | | | | | |
| Two ratio Model 2 | 70 | -42643.975206 | ω: | ω0=0.06250 | ω1=0.02993 | Model 0 vs. Two ratio Model 2 | 0.004342929 | ω1=0.02993 | *nad5* | |

| | | | | | | | | | | |
|---|---|---|---|---|---|---|---|---|---|---|
| Model 0 | 69 | -42648.042488 | ω= | 0.06077 | | | | | | |
| Two ratio Model 2 | 70 | -11161.910962 | ω: | ω0=0.04608 | ω1=0.01015 | Model 0 vs. Two ratio Model 2 | 0.045732562 | ω1=0.01015 | *nad6* | |
| Model 0 | 69 | -11163.906671 | ω= | 0.04373 | | | | | | |
| Two ratio Model 2 | 70 | -24609.090352 | ω: | ω0=0.02152 | ω1=0.00488 | Model 0 vs. Two ratio Model 2 | 0.000163735 | ω1=0.00488 | *cox1* | *S. allegheniensis* |
| Model 0 | 69 | -24616.193996 | ω= | 0.02099 | | | | | | |
| Two ratio Model 2 | 70 | -16205.109405 | ω: | ω0=0.04350 | ω1=0.01645 | Model 0 vs. Two ratio Model 2 | 0.026120839 | ω1=0.01645 | *cox3* | |
| Model 0 | 69 | -16207.583406 | ω= | 0.04274 | | | | | | |
| Two ratio Model 2 | 70 | -24613.689587 | ω: | ω0=0.02117 | ω1=0.00359 | Model 0 vs. Two ratio Model 2 | 0.025218520 | ω1=0.00359 | *cox1* | *S. pizzinii* |
| Model 0 | 69 | -24616.193996 | ω= | 0.02099 | | | | | | |
| Two ratio Model 2 | 70 | -12958.490043 | ω: | ω0=0.03112 | ω1=0.00477 | Model 0 vs. Two ratio Model 2 | 0.019738673 | ω1=0.00477 | *cox2* | |
| Model 0 | 69 | -12961.207473 | ω= | 0.03033 | | | | | | |
| Two ratio Model 2 | 70 | -23400.497996 | ω: | ω0=0.03415 | ω1=0.00760 | Model 0 vs. Two ratio Model 2 | 0.004692619 | ω1=0.00760 | *cytb* | |
| Model 0 | 69 | -23404.495120 | ω= | 0.03337 | | | | | | |
| Two ratio Model 2 | 70 | -29487.040979 | ω: | ω0=0.04390 | ω1=0.01007 | Model 0 vs. Two ratio Model 2 | 0.005384643 | ω1=0.01007 | *nad4* | |
| Model 0 | 69 | -29490.913731 | ω= | 0.04296 | | | | | | |
| Two ratio Model 2 | 70 | -24606.402409 | ω: | ω0=0.02171 | ω1=0.00333 | Model 0 vs. Two ratio Model 2 | 0.000009631 | ω1=0.00333 | *cox1* | *S. indentatus* |
| Model 0 | 69 | -24616.193996 | ω= | 0.02099 | | | | | | |
| Two ratio Model 2 | 70 | -19831.073084 | ω: | ω0=0.04595 | ω1=0.00102 | Model 0 vs. Two ratio Model 2 | 0.000440467 | ω1=0.00102 | *nad2* | |
| Model 0 | 69 | -19837.249185 | ω= | 0.04445 | | | | | | |

**Supplementary Table S7.** Evidence of positive selection on the mitochondrial PCGs with LRT P-value < 0.05 and positively selected site (BEB: P≥95%) in subterranean and surface dwelling lineages of amphipods based on branch-site models.

| Species | Gene | Model | np | Ln L | Estimates of parameters | | | | | Model compared | LRT P-value | Positive sites (BEB: P≥95%) |
|---|---|---|---|---|---|---|---|---|---|---|---|---|
| *P. kessleri* | *atp8* | Model A | 72 | -3843.85 | Site class | 0 | 1 | 2a | 2b | Model A vs.Model A null | 0.0007 | 4 S 1.000**,5 N 0.978*,6 W 1.000**,8 F 1.000**,14 L 0.969*,16 I 0.981*,20 M 0.976*,21 N 0.952*,24 L 0.999**,28 S 0.988*,33 L 0.951*,34 N 0.958*,35 N 0.993**,37 A 0.986* |
| | | | | | f | 0.23149 | 0.32042 | 0.18795 | 0.26014 | | | |
| | | | | | ω0 | 0.12034 | 1.00000 | 0.12034 | 1.00000 | | | |

| | | | | | | | | | | | | |
|---|---|---|---|---|---|---|---|---|---|---|---|---|
| | | | | | ω1 | 0.12034 | 1.00000 | 999.00000 | 999.00000 | | | |
| | | Model A null | 71 | -3849.58 | 1 | | | | | | | Not Allowed |
| | *nad5* | Model A | 72 | -42215.62 | Site class | 0 | 1 | 2a | 2b | Model A vs.Model A null | 0.0468 | 12 G 0.992**,34 W 0.993**,60 T 0.964*,91 M 0.966*,358 L 0.971*,457 K 0.964*,477 S 0.979*,511 S 0.960* |
| | | | | | f | 0.76401 | 0.11614 | 0.10404 | 0.01581 | | | |
| | | | | | ω0 | 0.06917 | 1.00000 | 0.06917 | 1.00000 | | | |
| | | | | | ω1 | 0.06917 | 1.00000 | 2.17571 | 2.17571 | | | |
| | | Model A null | 71 | -42217.60 | 1 | | | | | | | Not Allowed |
| *G. antarctica* | *nad5* | Model A | 72 | -42207.04 | Site class | 0 | 1 | 2a | 2b | Model A vs.Model A null | 0.0045 | 35 G 0.996**,40 N 0.971*,46 F 0.994**,47 S 0.957*,59 S 0.985*,131 S 0.986*,171 S 0.953*,192 S 0.989*,239 I 0.965*,283 E 0.993**,381 V 0.980*,389 M 0.993**,429 W 0.953*,472 G 0.987*,473 L 0.991**,477 S 0.976*,488 K 0.962* |
| | | | | | f | 0.69413 | 0.11323 | 0.16562 | 0.02702 | | | |
| | | | | | ω0 | 0.07063 | 1.00000 | 0.07063 | 1.00000 | | | |
| | | | | | ω1 | 0.07063 | 1.00000 | 42.86420 | 42.86420 | | | |
| | | Model A null | 71 | -42211.07 | 1 | | | | | | | Not Allowed |
| | *nad6* | Model A | 72 | -11115.29 | Site class | 0 | 1 | 2a | 2b | Model A vs.Model A null | 0.0019 | 51 G 0.989*,86 N 0.982*,100 S 0.964*,108 L 0.954*,128 G 0.974* |
| | | | | | f | 0.60982 | 0.15741 | 0.18502 | 0.04776 | | | |
| | | | | | ω0 | 0.07302 | 1.00000 | 0.07302 | 1.00000 | | | |
| | | | | | ω1 | 0.07302 | 1.00000 | 1.50615 | 1.50615 | | | |
| | | Model A null | 71 | -11110.47 | 1 | | | | | | | Not Allowed |
| *P. japonica* | *nad2* | Model A | 72 | -19775.06 | Site class | 0 | 1 | 2a | 2b | Model A vs.Model A null | 0.0172 | 116 D 0.979*, 232 S 0.970* |
| | | | | | f | 0.92964 | 0.04229 | 0.02685 | 0.00122 | | | |
| | | | | | ω0 | 0.04908 | 1.00000 | 0.04908 | 1.00000 | | | |
| | | | | | ω1 | 0.04908 | 1.00000 | 855.94976 | 855.94976 | | | |
| | | Model A null | 71 | -19777.90 | 1 | | | | | | | Not Allowed |
| | *nad6* | Model A | 72 | -11114.94 | Site class | 0 | 1 | 2a | 2b | Model A vs.Model A null | 0.0420 | 86 N 0.982* |
| | | | | | f | 0.80724 | 0.14323 | 0.04207 | 0.00746 | | | |
| | | | | | ω0 | 0.06706 | 1.00000 | 0.06706 | 1.00000 | | | |
| | | | | | ω1 | 0.06706 | 1.00000 | 998.99536 | 998.99536 | | | |
| | | Model A null | 71 | -11117.00 | 1 | | | | | | | Not Allowed |

| | | | | | | | | | | | | |
|---|---|---|---|---|---|---|---|---|---|---|---|---|
| *C. mutica* | *nad1* | Model A | 72 | -19367.63 | Site class | 0 | 1 | 2a | 2b | Model A vs.Model A null | 0.0007 | 163 I 0.997**,243 S 0.984*,249 V 0.989* |
| | | | | | f | 0.94079 | 0.02466 | 0.03367 | 0.00088 | | | |
| | | | | | ω0 | 0.03489 | 1.00000 | 0.03489 | 1.00000 | | | |
| | | | | | ω1 | 0.03489 | 1.00000 | 14.28437 | 14.28437 | | | |
| | | Model A null | 71 | -19373.41 | 1 | | | | | | | Not Allowed |
| | *nad5* | Model A | 72 | -42225.68 | Site class | 0 | 1 | 2a | 2b | Model A vs.Model A null | 0.0030 | 280 G 0.975*,326 G 0.975*,400 S 0.989* |
| | | | | | f | 0.79382 | 0.13139 | 0.06417 | 0.01062 | | | |
| | | | | | ω0 | 0.07070 | 1.00000 | 0.07070 | 1.00000 | | | |
| | | | | | ω1 | 0.07070 | 1.00000 | 9.66169 | 9.66169 | | | |
| | | Model A null | 71 | -42230.09 | 1 | | | | | | | Not Allowed |
| | *cox2* | Model A | 72 | -12930.83 | Site class | 0 | 1 | 2a | 2b | Model A vs.Model A null | 0.0115 | 142 K 0.986* |
| | | | | | f | 0.97663 | 0.00901 | 0.01423 | 0.00013 | | | |
| | | | | | ω0 | 0.03012 | 1.00000 | 0.03012 | 1.00000 | | | |
| | | | | | ω1 | 0.03012 | 1.00000 | 268.52413 | 268.52413 | | | |
| | | Model A null | 71 | -12934.02 | 1 | | | | | | | Not Allowed |
| *G. fossarum* | *nad3* | Model A | 72 | -7315.91 | Site class | 0 | 1 | 2a | 2b | Model A vs.Model A null | 0.0439 | 6 F 0.991**,17 L 0.999**,22 H 0.979*,23 S 0.999**,25 P 0.969*,26 S 1.000**,76 P 0.978*,82 T 0.972*,84 L 0.998**,91 V 0.974*,93 L 0.966*,94 I 0.999** |
| | | | | | f | 0.59041 | 0.08730 | 0.28077 | 0.04152 | | | |
| | | | | | ω0 | 0.03895 | 1.00000 | 0.03895 | 1.00000 | | | |
| | | | | | ω1 | 0.03895 | 1.00000 | 11.02113 | 11.02113 | | | |
| | | Model A null | 71 | -7317.94 | 1 | | | | | | | Not Allowed |
| *O. nanseni* | *atp6* | Model A | 72 | -15167.07 | Site class | 0 | 1 | 2a | 2b | Model A vs.Model A null | 0.0478 | 69 M 0.986*, 146 N 0.981*, 172 S 0.974*, 186 A 0.969*, 188 G 0.981*, 189 L 0.996** |
| | | | | | f | 0.76666 | 0.07611 | 0.14304 | 0.01420 | | | |
| | | | | | ω0 | 0.05002 | 1.00000 | 0.05002 | 1.00000 | | | |
| | | | | | ω1 | 0.05002 | 1.00000 | 18.13180 | 18.13180 | | | |
| | | Model A null | 71 | -15169.03 | 1 | | | | | | | Not Allowed |
| | *nad5* | Model A | 72 | -42207.40 | Site class | 0 | 1 | 2a | 2b | Model A vs.Model A null | 0.0004 | 3 S 0.991**, 6 A 0.988*, 73 S 0.985*, 98 F 0.972*, 130 S 0.968*, 162 W 0.961*, 179 I 0.960*, 242 M 0.955*, 325 W 0.976* , 341 C 0.995**, 367 L 0.990**, 373 V 0.953*, 377 G 0.974*, 427 M 0.987*, 465 N 0.989*, 479 W |
| | | | | | f | 0.72687 | 0.12217 | 0.12924 | 0.02172 | | | |
| | | | | | ω0 | 0.07034 | 1.00000 | 0.07034 | 1.00000 | | | |

| | | | | | | | | | | | | |
|---|---|---|---|---|---|---|---|---|---|---|---|---|
| | | | | | ω1 | 0.07034 | 1.00000 | 62.60443 | 62.60443 | | | 0.977*, 514 L 0.969*, 515 Q 0.989*, 518 Q 0.999**, 519 S 0.985*, 523 S 0.959* |
| | | Model A null | 71 | -42213.74 | 1 | | | | | | | Not Allowed |
| *G. fasciatus* | *atp8* | Model A | 72 | -4021.06 | Site class | 0 | 1 | 2a | 2b | Model A vs.Model A null | 0.0000 | 7 S 0.999**, 12 F 0.976* |
| | | | | | f | 0.25262 | 0.42481 | 0.12029 | 0.20228 | | | |
| | | | | | ω0 | 0.09609 | 1.00000 | 0.09609 | 1.00000 | | | |
| | | | | | ω1 | 0.09609 | 1.00000 | 327.33424 | 327.33424 | | | |
| | | Model A null | 71 | -3851.18 | 1 | | | | | | | Not Allowed |
| | *nad2* | Model A | 72 | -19783.73 | Site class | 0 | 1 | 2a | 2b | Model A vs.Model A null | 0.0000 | 117 M 0.960*, 191 K 0.967* |
| | | | | | f | 0.80648 | 0.03554 | 0.15131 | 0.00667 | | | |
| | | | | | ω0 | 0.04922 | 1.00000 | 0.04922 | 1.00000 | | | |
| | | | | | ω1 | 0.04922 | 1.00000 | 1.00000 | 1.00000 | | | |
| | | Model A null | 71 | -19775.46 | 1 | | | | | | | Not Allowed |
| | *nad5* | Model A | 72 | -42209.09 | Site class | 0 | 1 | 2a | 2b | Model A vs.Model A null | 0.0001 | 17 V 0.993**, 27 N 0.977*, 72 S 0.978*, 159 S 0.972*, 161 G 0.994**, 183 V 0.961*, 242 M 0.976*, 243 M 0.978*, 313 S 0.966*, 323 G 0.985*, 368 C 0.960*, 376 S 0.962*, 388 S 0.996**, 389 M 0.852, 391 V 0.984*, 474 A 0.997**, 514 L 0.976* |
| | | | | | f | 0.75033 | 0.12088 | 0.11092 | 0.01787 | | | |
| | | | | | ω0 | 0.06986 | 1.00000 | 0.06986 | 1.00000 | | | |
| | | | | | ω1 | 0.06986 | 1.00000 | 94.94243 | 94.94243 | | | |
| | | Model A null | 71 | -42216.31 | 1 | | | | | | | Not Allowed |
| *C. forbesi* | *nad5* | Model A | 72 | -42180.92 | Site class | 0 | 1 | 2a | 2b | Model A vs.Model A null | 0.0000 | 2 F 0.982*, 20 S 0.986*, 23 V 0.994**, 40 N 0.978*, 48 L 0.979*, 56 V 0.970*, 78 G 0.991**, 82 I 0.993**, 88 I 0.958*, 121 A 0.997**, 131 S 0.992**, 132 Q 0.999**, 152 S 0.981*, 157 S 0.979*, 159 S 0.989*, 165 V 0.965*, 178 I 0.962*, 181 A 0.990*, 192 S 0.998**, 200 A 0.997**, 250 L 0.989*, 253 N 0.980*, 254 Y 0.991**, 271 G 0.995**, 278 S 0.989*, 283 E 0.989*, 323 G 0.978*, 333 F 0.966*, 336 C 0.991**, 337 N 0.999**, 344 P 0.957*, 347 S 0.996**, 354 L 0.983*, 355 V 0.956*, 359 M 0.725, 360 L 0.956*, 361 S 0.996**, 373 V 0.989*, 375 S 0.988*, 384 V 0.989*, 411 G 0.999**, 412 G 0.956*, 417 F 0.984*, 444 S 0.981*, 448 L 0.984*, 465 N 0.986*, 472 G 0.996**, 473 L 0.994**, 474 A 0.998**, 495 L 0.997**, 502 G 0.970*, 503 G 0.992**, 507 F 0.991**, 511 S 0.994**, 519 S 0.982*, 523 S 0.996**, 538 V 0.977* |
| | | | | | f | 0.58855 | 0.10272 | 0.26286 | 0.04588 | | | |
| | | | | | ω0 | 0.07187 | 1.00000 | 0.07187 | 1.00000 | | | |
| | | | | | ω1 | 0.07187 | 1.00000 | 63.07884 | 63.07884 | | | |

| | | | | | | | | | | | | |
|---|---|---|---|---|---|---|---|---|---|---|---|---|
| | | Model A null | 71 | -42191.63 | 1 | | | | | | | Not Allowed |
| | *nad6* | Model A | 72 | -11102.90 | Site class | 0 | 1 | 2a | 2b | Model A vs.Model A null | 0.0077 | 3 I 1.000**, 7 L 1.000**, 12 L 0.997**, 20 L 0.997**, 23 T 0.984*, 28 V 0.958*, 29 A 0.996**, 35 W 0.996**, 42 A 0.996**, 49 F 0.964*, 50 L 0.970*, 58 T 0.992**, 61 T 0.999**, 69 T 1.000**, 85 N 0.994**, 87 L 0.990*, 95 H 1.000**, 96 K 0.995**, 97 I 0.956*, 100 S 0.996**, 101 G 0.982*, 102 T 0.956*, 103 E 0.995**, 106 T 0.977*, 129 S 1.000**, 133 S 0.986* |
| | | | | | f | 0.47135 | 0.07130 | 0.39725 | 0.06009 | | | |
| | | | | | ω0 | 0.06479 | 1.00000 | 0.06479 | 1.00000 | | | |
| | | | | | ω1 | 0.06479 | 1.00000 | 61.36780 | 61.36780 | | | |
| | | Model A null | 71 | -11106.46 | 1 | | | | | | | Not Allowed |
| | *cox1* | Model A | 72 | -24421.90 | Site class | 0 | 1 | 2a | 2b | Model A vs.Model A null | 0.0385 | 155 E 0.972* |
| | | | | | f | 0.90038 | 0.02854 | 0.06889 | 0.00218 | | | |
| | | | | | ω0 | 0.01865 | 1.00000 | 0.01865 | 1.00000 | | | |
| | | | | | ω1 | 0.01865 | 1.00000 | 74.69725 | 74.69725 | | | |
| | | Model A null | 71 | -24424.04 | 1 | | | | | | | Not Allowed |
| | *cytb* | Model A | 72 | -23302.46 | Site class | 0 | 1 | 2a | 2b | Model A vs.Model A null | 0.0000 | 4 S 1.000**, 24 L 0.961*, 50 S 0.999**, 52 T 0.985*, 53 L 0.997**, 81 C 1.000**, 89 G 0.999**, 96 L 0.997**, 97 Q 1.000**, 99 H 0.999**, 111 T 0.997**, 147 D 0.997**, 150 K 0.958*, 179 A 0.999**, 181 A 0.966*, 197 L 0.998**, 237 L 0.998**, 244 T 0.984*, 250 T 0.979*, 253 I 0.999**, 273 N 0.984*, 279 L 0.968*, 282 L 0.993**, 283 L 0.998**, 294 T 0.998**, 298 K 0.999**, 307 N 1.000**, 324 L 0.967*, 335 F 0.997**, 356 T 0.998** |
| | | | | | f | 0.00003 | 0.00000 | 0.96156 | 0.03841 | | | |
| | | | | | ω0 | 0.03404 | 1.00000 | 0.03404 | 1.00000 | | | |
| | | | | | ω1 | 0.03404 | 1.00000 | 1.00000 | 1.00000 | | | |
| | | Model A null | 71 | -23269.10 | 1 | | | | | | | Not Allowed |
| *P. daviui* | *nad4* | Model A | 72 | -29345.91 | Site class | 0 | 1 | 2a | 2b | Model A vs.Model A null | 0.0134 | 23 S 0.951*, 37 S 0.970*, 48 S 0.974*, 60 A 0.989*, 84 L 0.983*, 260 V 0.971*, 284 S 0.996**, 319 W 0.959*, 379 E 0.973*, 390 G 0.990**, 411 S 0.958*, 413 I 0.956*, 414 F 0.989* |
| | | | | | f | 0.78376 | 0.04107 | 0.16645 | 0.00872 | | | |
| | | | | | ω0 | 0.04528 | 1.00000 | 0.04528 | 1.00000 | | | |
| | | | | | ω1 | 0.04528 | 1.00000 | 13.69295 | 13.69295 | | | |
| | | Model A null | 71 | -29348.97 | 1 | | | | | | | Not Allowed |
| | *nad5* | Model A | 72 | -42222.23 | Site class | 0 | 1 | 2a | 2b | Model A vs.Model A null | 0.0362 | 6 A 0.970*, 20 S 0.955*, 23 V 0.982*, 132 Q 0.992**, 148 G 0.953*, 157 S 0.953*, 232 S 0.951*, 282 I 0.976*, 285 A 0.970*, 332 I 0.956*, 461 T 0.958*, 486 K 0.990* |
| | | | | | f | 0.73648 | 0.11827 | 0.12515 | 0.02010 | | | |
| | | | | | ω0 | 0.07092 | 1.00000 | 0.07092 | 1.00000 | | | |
| | | | | | ω1 | 0.07092 | 1.00000 | 43.91410 | 43.91410 | | | |

| | | | | | | | | | | | | |
|---|---|---|---|---|---|---|---|---|---|---|---|---|
| | | Model A null | 71 | -42224.43 | 1 | | | | | | | Not Allowed |
| *B. jaraguensis* | *nad2* | Model A | 72 | -19779.55 | Site class | 0 | 1 | 2a | 2b | Model A vs.Model A null | 0.0001 | 162 M 0.959*, 233 F 0.973* |
| | | | | | f | 0.79491 | 0.03611 | 0.16164 | 0.00734 | | | |
| | | | | | ω0 | 0.04900 | 1.00000 | 0.04900 | 1.00000 | | | |
| | | | | | ω1 | 0.04900 | 1.00000 | 1.28390 | 1.28390 | | | |
| | | Model A null | 71 | -19771.49 | 1 | | | | | | | Not Allowed |
| *M. dominicanus* | *nad3* | Model A | 72 | -7323.42 | Site class | 0 | 1 | 2a | 2b | Model A vs.Model A null | 0.0190 | 81 N 0.995** |
| | | | | | f | 0.85321 | 0.12507 | 0.01895 | 0.00278 | | | |
| | | | | | ω0 | 0.03891 | 1.00000 | 0.03891 | 1.00000 | | | |
| | | | | | ω1 | 0.03891 | 1.00000 | 161.98168 | 161.98168 | | | |
| | | Model A null | 71 | -7326.17 | 1 | | | | | | | Not Allowed |
| *B. brachycaudus* | *atp6* | Model A | 72 | -15165.74 | Site class | 0 | 1 | 2a | 2b | Model A vs.Model A null | 0.0206 | 10 S 0.992**, 66 S 0.969*, 79 S 0.987*, 119 N 0.971*, 128 I 0.990**, 147 N 0.974*, 177 Y 0.982*, 184 G 0.990* |
| | | | | | f | 0.70794 | 0.06884 | 0.20344 | 0.01978 | | | |
| | | | | | ω0 | 0.05148 | 1.00000 | 0.05148 | 1.00000 | | | |
| | | | | | ω1 | 0.05148 | 1.00000 | 44.53036 | 44.53036 | | | |
| | | Model A null | 71 | -15168.42 | 1 | | | | | | | Not Allowed |
| | *nad5* | Model A | 72 | -42197.28 | Site class | 0 | 1 | 2a | 2b | Model A vs.Model A null | 0.0001 | 1 S 0.977*, 12 G 0.959*, 13 S 0.991**, 45 S 0.980*, 73 S 0.972*, 74 S 0.953*, 97 G 0.956*, 130 S 0.972*, 132 Q 0.999**, 163 G 0.984*, 229 S 0.987*, 312 I 0.986*, 330 L 0.992**, 336 C 0.986*, 365 S 0.981*, 375 S 0.982*, 388 S 0.983*, 391 V 0.976*, 409 S 0.966*, 411 G 0.990**, 423 L 0.988*, 448 L 0.985*, 469 Y 0.983*, 472 G 0.991**, 479 W 0.991**, 485 Y 0.957*, 493 S 0.989*, 497 E 0.975*, 498 V 0.993**, 500 P 0.991**, 519 S 0.990* |
| | | | | | f | 0.65345 | 0.10879 | 0.20383 | 0.03393 | | | |
| | | | | | ω0 | 0.07113 | 1.00000 | 0.07113 | 1.00000 | | | |
| | | | | | ω1 | 0.07113 | 1.00000 | 54.77953 | 54.77953 | | | |
| | | Model A null | 71 | -42204.57 | 1 | | | | | | | Not Allowed |
| *S. allegheniensis* | *nad3* | Model A | 72 | -7325.19 | Site class | 0 | 1 | 2a | 2b | Model A vs.Model A null | 0.0375 | 77 V 0.990* |
| | | | | | f | 0.85787 | 0.12736 | 0.01286 | 0.00191 | | | |
| | | | | | ω0 | 0.03868 | 1.00000 | 0.03868 | 1.00000 | | | |
| | | | | | ω1 | 0.03868 | 1.00000 | 108.89135 | 108.89135 | | | |
| | | Model A null | 71 | -7327.36 | 1 | | | | | | | Not Allowed |

| | | | | | | | | | | | | |
|---|---|---|---|---|---|---|---|---|---|---|---|---|
| *S. indentatus* | *atp8* | Model A | 72 | -4020.49 | Site class | 0 | 1 | 2a | 2b | Model A vs.Model A null | 0.0028 | 12 F 0.994**, 14 I 0.981* |
| | | | | | f | 0.30650 | 0.45936 | 0.09370 | 0.14043 | | | |
| | | | | | ω0 | 0.09877 | 1.00000 | 0.09877 | 1.00000 | | | |
| | | | | | ω1 | 0.09877 | 1.00000 | 999.00000 | 999.00000 | | | |
| | | Model A null | 71 | -4024.97 | 1 | | | | | | | Not Allowed |
| | *nad5* | Model A | 72 | -42219.00 | Site class | 0 | 1 | 2a | 2b | Model A vs.Model A null | 0.0001 | 3 S 0.974*, 59 S 0.991**, 171 S 0.967*, 310 S 0.989*, 321 V 0.970*, 419 L 0.982*, 468 A 0.985*, 469 Y 0.968*, 486 K 0.986* |
| | | | | | f | 0.79435 | 0.13501 | 0.06038 | 0.01026 | | | |
| | | | | | ω0 | 0.07100 | 1.00000 | 0.07100 | 1.00000 | | | |
| | | | | | ω1 | 0.07100 | 1.00000 | 13.75919 | 13.75919 | | | |
| | | Model A null | 71 | -42227.16 | 1 | | | | | | | Not Allowed |
| | *cox3* | Model A | 72 | -16038.31 | Site class | 0 | 1 | 2a | 2b | Model A vs.Model A null | 0.0150 | 118 T 0.989* |
| | | | | | f | 0.92090 | 0.06266 | 0.01539 | 0.00105 | | | |
| | | | | | ω0 | 0.03997 | 1.00000 | 0.03997 | 1.00000 | | | |
| | | | | | ω1 | 0.03997 | 1.00000 | 29.92053 | 29.92053 | | | |
| | | Model A null | 71 | -16041.27 | 1 | | | | | | | Not Allowed |
